# INSIHGT: An accessible multi-scale, multi-modal 3D spatial biology platform

**DOI:** 10.1101/2024.05.24.595771

**Authors:** Chun Ngo Yau, Jacky Tin Shing Hung, Robert A. A. Campbell, Thomas Chun Yip Wong, Bei Huang, Ben Tin Yan Wong, Nick King Ngai Chow, Lichun Zhang, Eldric Pui Lam Tsoi, Yuqi Tan, Joshua Jing Xi Li, Yun Kwok Wing, Hei Ming Lai

## Abstract

Biological systems are complex, encompassing intertwined spatial, molecular and functional features. However, methodological constraints always limit the completeness of information that can be extracted. Here, we report the development of INSIHGT, a minimally perturbative, accessible and cost-efficient three-dimensional (3D) spatial biology method utilizing superchaotropes and host-guest chemistry. This allows highly multiplexed and multi-modal readout of tissue biomolecules in biological systems up to centimeter scales, permitting radio-histological correlation of phosphorylated alpha-synuclein pathologies in human hemi-brainstem. The homogeneous penetration permits reliable semi-quantitative signals in 3D compared to reference signals. Diverse antigens, mRNA transcripts, neurotransmitters, and post-translational and epigenetic modifications, are well-preserved and visualized. INSIHGT also allows multi-round molecular probing for high-dimensional spatial biology and compatibility with downstream traditional histology. With INSIHGT, we mapped previously undescribed podocyte-to-parietal epithelial cell microfilaments and demonstrated their geodesic clustering in mouse glomeruli, and catalogued sparsely located neurofilament-intensive inclusion bodies in the human cerebellum, and identified NPY-proximal cell types defined by spatial morpho-proteomics in mouse hypothalamus. We anticipate INSIHGT can form the foundations for 3D spatial multi-omics technology development and holistic systems biology studies.

## Main

The complexity of biological systems mandates high-dimensional measurements to obtain an integrative understanding. However, measurements are inevitably perturbative, affecting the authenticity of the retrieved information. Spatially resolved transcriptomics ^1^ and highly multiplexed immunohistochemistry (IHC) ^2^ have proven to be powerful approaches to extract spatial molecular insights from tissue slices, but the two-dimensional (2D) readout limits the representativeness of the information extracted. Meanwhile, three-dimensional (3D) multiplexed visualization of tissue structural and molecular features can reveal previously unknown organization principles ^3,4^. Optical tissue clearing technologies promises to reveal the authentic 3D nature of tissue architecture and molecular distributions ^5^. Despite its significant advancements, the achievable depths of probe penetration limits the depth of analysis ^6^. The limited penetration of antibodies in 3D immunohistochemistry (IHC) represents one of the most significant barrier to 3D spatial biology ^6^.

In recent years, multiple creative solutions have been proposed for deep immunohistochemistry ^7–10^. However, an accessible technology that balances the authenticity and volume of data extracted is still lacking. For example, signal homogeneity across penetration depth is suboptimal with most methods, where probes preferentially deposit near the tissue surface and complicates downstream quantitative protein expression determination ^6,11^. The homogeneous penetration can only be attained either through complicated operations or equipment ^11–13^, or extensive tissue permeabilization ^14,15^ or incubation times measuring in weeks ^8^. These shortcomings hinder the wide adoption of 3D tissue analysis in research and renders them unsatisfactory for clinical translation.

To address these technical bottlenecks, we report the development of *<u>In</u> <u>s</u>*<u>i</u>tu *<u>H</u>*ost-*<u>G</u>*uest Chemistry for *<u>T</u>*hree-dimensional Histology (INSIHGT). INSIHGT is a user-friendly 3D histochemistry method, featuring (1) homogeneous probe penetration up to centimeter depths, (2) producing quantitative, highly specific immunostaining signals, (3) a fast and affordable workflow to accommodate different tissue sizes and shapes, (4) simple immersion-based staining at room temperature, thus easily adopted in any laboratory and ready for scaling and automate, and (5) uses off-the-shelf antibodies or probes and is directly applicable to otherwise unlabelled mouse and human tissues fixed with paraformaldehyde only. INSIHGT was developed based on the manipulation of chemical potential gradients that drive diffusiophoresis using an *in situ* supramolecular reaction system. Specifically, a boron cluster compound *closo*-dodecahydrododecaborate [B_12_H_12_]^2–16^ and a γ-cyclodextrin derivative were used to achieve switchable modulation of probe motions throughout the tissue and attain homogeneous, deeply penetrating 3D histochemistry.

## Results

The limited penetration of macromolecular probes in complex biological systems belongs to the broader subject of transport phenomena, where diffusion and advections respectively drive the dissipation and directional drift of mass, energy and momentum. When biomolecules such as proteins are involved, the (bio)molecular fluxes are additionally determined by binding reactions, which can significantly deplete biomolecules due to their high binding affinities and low concentrations employed - a “reaction barrier” to deep antibody penetration. This is first described and postulated by Renier et al. ^17^ (as in immunolabeling-enabled three-dimensional imaging of solvent-cleared organs, iDISCO) and Murray et al. ^14^ (as in system-wide control of interaction time and kinetics of chemicals, SWITCH), and the latter further showed that the modulation of antibody-antigen (Ab-Ag) binding affinity (SWITCH labelling) can lead to homogeneous penetration of up to 1 millimetre for an anti-Histone H3 antibody using low concentrations of sodium dodecyl sulphate (SDS). Other techniques similarly utilises urea ^8^, sodium deoxycholate ^12^, and heat ^9^ to modulate antibody-antigen binding.

However, others and we observed a general compromise between antibody labelling quality, penetration depth and uniformity, and duration of incubation. Deep penetration invariably requires long incubation times with inhomogeneous signal across depth, while faster methods leads to weak or nonspecific staining, as well as non-uniform penetration ^8,9,17^. Specifically, the use of SDS for deep labelling with SWITCH labelling has only been demonstrated for a handful of antigens (e.g., Histone H3 ^14^, NeuN ^18^, ColIV , αSMA , and TubIII ^19^). It was found that deep staining with SDS was not universally applicable ^18^, resulting in weak calbindin staining ^20^, insufficient staining depth for β-amyloid plaques ^21^, and often requires tailored refinement of buffer concentration ^22^. In our validation data, we similarly observed the variable performance when SDS is co-applied with antibodies (**Fig. S1**). Furthermore, although adding antibodies or probes theoretically improves penetration via steep concentration gradients, either the cost becomes prohibitive or it produces a biased representation of rimmed surface staining pattern ^6,8^, especially for densely expressed binding targets. In the most extreme cases, the superficial staining signal would saturate microscope detectors while the core remains unstained (**Fig. S2**).

Nonetheless, the conception of modulating antibody-antigen binding kinetics as a means to control probe flux through tissues is highly attractive ^12,14^, given the simplicity, scalability, and affordability should the method be robust and generalizable. We postulated that the reason for the highly variable performance of SDS-assisted deep immunostaining is two-fold: the denaturation of antibodies beyond reparability, and the ineffective reinstatement of binding reactions. This prompted us to search for alternative approaches that can tune biomolecular binding affinities while preserving both macromolecular probe mobility and stability. In addition, the negation of modulatory effect should be efficient and robust to reinstate biomolecular reactions within the complex tissue environment. Therefore, here we aim to develop a fast, equipment-free, deep and uniform multiplexed immunostaining method, which will help bring 3D histology to any basic laboratories.

### Boron cluster host-guest chemistry for in situ macromolecular probe mobility control

Our initial attempts by using heat and the GroEL-GroES system to denature and refold antibodies *in situ* respectively has proved unsuccessful (**Fig. S1**). We thus switched from the natural molecular chaperones to artificial ones using milder detergents (e.g., sodium deoxycholate (SDC) and 3-([3-Cholamidopropyl]dimethylammonio)-2-hydroxy-1-propanesulfonate i.e., CHAPSO) and their charge-complementary, size-matched host-complexing agents (e.g., β-cyclodextrins and their derivatives such as heptakis-(6-amino-6-deoxy)-beta-Cyclodextrin, i.e., 6NβCD), which improved antibody penetration and staining success rate (**Fig. S3**). However, despite extensive optimization on the structure and derivatization on the detergents and their size- and charge-complementary cyclodextrins, they still have limited generality for a panel of antibodies tested (**Fig. S3**), producing non-specific vascular precipitates or nuclear stainings. We then explored the use of chaotropes, which are known to solubilize proteins with enhanced antibody penetration ^8^. However, these approaches require long incubation times with extensive tissue pre-processing. Furthermore, higher concentrations of chaotropes often denature proteins as they directly interact with various protein residues and backbone ^23,24^ (**Fig. 1A, B**).

**Figure 1.**
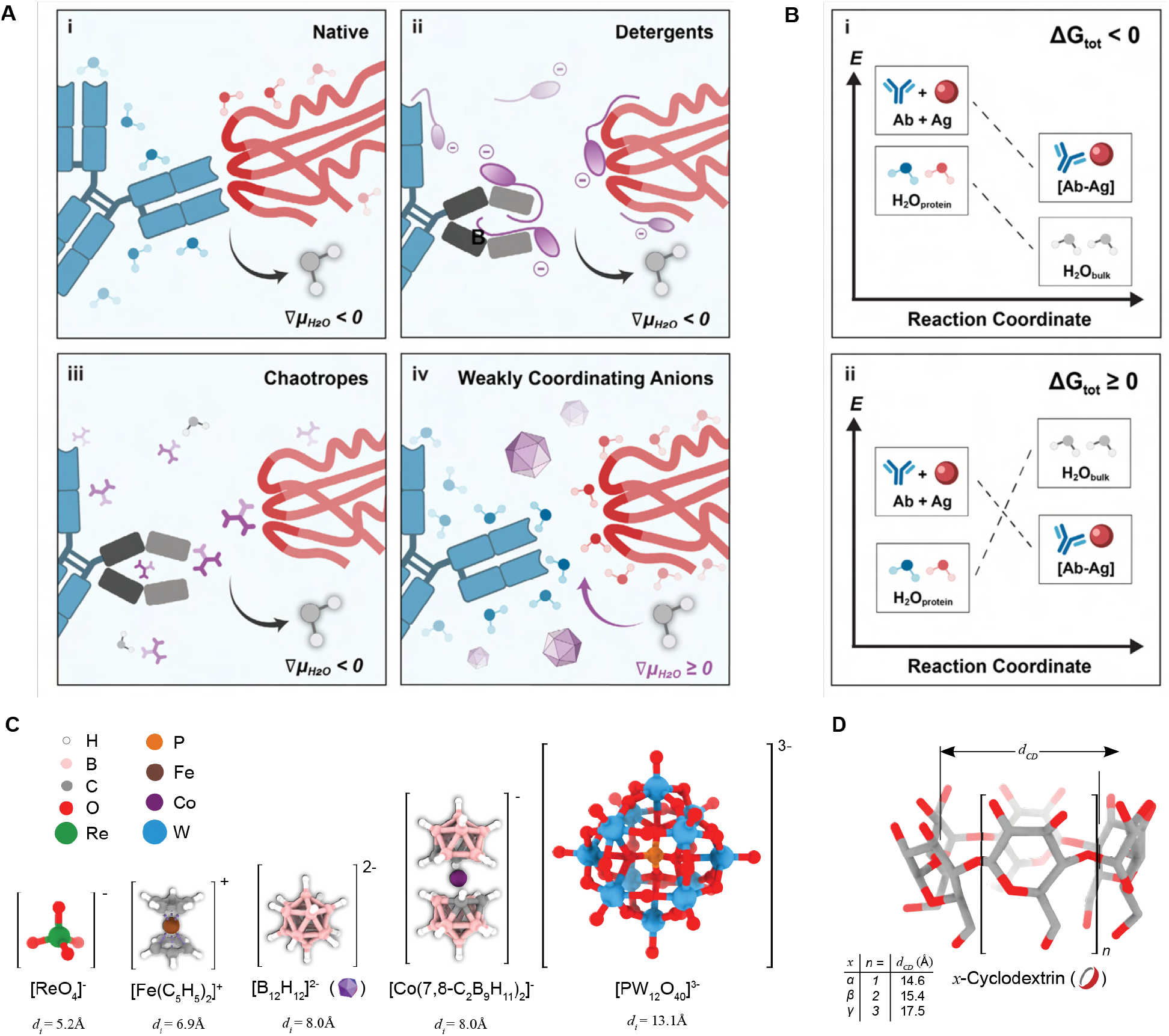
INSIHGT conceptualization and key components. (**A-B**) Reconceptualization of the biomolecular binding phenomena. (**A**) Microscopic view. (***i***) Native antibody-antigen binding (in an aqueous medium) requires the desolvation of their solvation shells (∇μ_H2O_< 0). (***ii, iii***) Detergents and chaotropes solubilize proteins by masking binding sites (and displacing the solvent shell), but they may lead to protein denaturation in high concentrations (black arm of the antibody). (***iv***) Weakly coordinating superchaotropes (WCS, e.g., [B_12_H_12_]^2-^) instead solubilizes proteins by favoring solvent-protein interactions (∇μ_H2O_ ≥ 0), striking a balance between protein solubilization and stabilization. (**B**) Energetic view. Antibody-antigen binding (***i***) without WCS, occurs spontaneously with ΔG_tot_ < 0, while (***ii***) with WCA, is unfavourable with ΔG_tot_ ≥ 0. (**C**) Structures and ionic diameters (*d*_i_) of weakly coordinating ions tested. (**D**) General structure of cyclodextrins and their cavity opening diameter (d_CD_).

We hence focus on testing weakly coordinating superchaotropes (WCS), a class of chemicals that we hypothesized to inhibit antibody-antigen interactions while preserving their structure and hence functions (**Fig. 1A, B**). We searched for weakly coordinating ions based on their utility in isolating extremely electrophilic species for X-ray crystallography, or as conjugate bases of superacids. After antibodies and WCS have been homogeneously distributed throughout the tissue matrix, measures must be taken to negate the superchaotropicity to reinstate inter-biomolecular interactions in a bio-orthogonal and system-wide manner. To do so, we took advantage of the enthalpy-driven chaotropic assembly reaction, where the activities of superchaotropes can be effectively negated with supramolecular hosts in situ, reactivating interactions between the macromolecular probes and their tissue targets.

Based on the above analysis, we designed a scalable deep molecular phenotyping method, performed in two stages: a first infiltrative stage where macromolecular probes co-diffuse homogeneously with WCS with minimized reaction barriers, followed by the addition of macrocyclic compounds for in situ host-guest reactions to reinstate antibody-antigen binding. With a much-narrowed list of chemicals to screen, we first benchmarked the performances of several putative WCS host-guest systems using a standard protocol as previously published ^6,8,9^ (**Fig. S4**). These include perrhenate/α-cyclodextrin (ReO_4_^-^/αCD), ferrocenium/βCD ([Fe(C_5_H_5_)_2_]^+^/βCD), *closo*-dodecaborate ions ([B_12_X_12_]^2-^/γCD (where X = H, Cl, Br, or I)), metallacarborane ([Co(7,8-C_2_B_9_H_11_)_2_]^-^/γCD), and polyoxometalates ([PM_12_O_40_]^3-^/γCD (where M = Mo, or W)) (**Fig. 1C-D**). Group 5 and 6 halide clusters and rhenium chalcogenide clusters such as [Ta_6_Br_12_]^2+^, [Mo_6_Cl_14_]^2-^ and {Re_6_Se_8_}^2+^ derivatives were excluded due to instability in aqueous environments. Only ReO_4_^-^, [B_12_H_12_]^2-^, and [Co(7,8-C_2_B_9_H_11_)_2_]^-^ proved compatible with immunostaining conditions without causing tissue destruction or precipitation. [B_12_H_12_]^2-^/γCD produced the best staining sensitivity, specificity and signal homogeneity across depth (**Fig. S5**), while the effect of derivatizing γCD was negligible (**Fig. S5**). Finally, we chose the more soluble 2-hydroxypropylated derivative (2HPγCD) for its higher water solubility in our applications. We term our new method INSIHGT, for <u>In si</u>tu <u>h</u>ost-<u>g</u>uest chemistry for three-dimensional histology.

### In situ host-guest chemistry for three-dimensional histology (INSIHGT)

INSIHGT was designed to be a minimally perturbative, deeply and homogeneously penetrating staining method for 3D histology. Designed for affordability and scalability, INSIHGT involves simply incubating the conventional formaldehyde-fixed tissues in [B_12_H_12_]^2-^/PBS with antibodies, then in 2HPγCD/PBS (**Fig. 2A**) - both at room temperature with no specialized equipment. We compared INSIHGT with other 3D IHC techniques using a stringent benchmarking experiment as previously published (see **Methods, Fig. S4**) to compare their penetration depths and homogeneity ^6,9^. We found that INSIHGT achieved the deepest immunolabeling penetration with the best signal homogeneity throughout the penetration depth (**Fig. 2B**). To quantitatively compare the signal, we segmented the labelled cells and compared the ratio between the deep immunolabelling signal and the reference signal against their penetration depths. Exponential decay curve fitting showed that the signal homogeneity was near-ideal (**Fig. 2C, Table S1**) - where there was negligible decay in deep immunolabelling signals across the penetration depth. We repeated our benchmarking experiment with different markers, and by correlating INSIHGT signal with the reference signal, we found INSIHGT provides reliable relative quantification of cellular marker expression levels throughout an entire mouse hemi-brain stained for 1 day (**Fig. 2D**). We supplemented our comparison with the binding kinetics modulating buffers employed in eFLASH and SWITCH-pumping of mELAST tissue-hydrogel, as we lacked the specialized equipment to provide the external force fields and mechanical compressions, respectively (**Fig. S6**). For SWITCH-pumping of mELAST tissue-hydrogel, we utilized the latest protocol and buffer recipe ^13^. Our results also showed the use of binding kinetics modulating buffers alone from eFLASH and SWITCH-pumping of mELAST tissue-hydrogel lead to shallower staining penetration than INSIHGT, confirming the deep penetration of these methods is mainly contributed by the added external force fields and mechanical compressions, respectively. Hence, with excellent penetration homogeneity with a simple operating protocol, INSIHGT can be the ideal method for mapping whole organs with cellular resolution. It is also the fastest deep immunolabelling from tissue harvesting to image (**Fig. 2E**).

**Figure 2.**
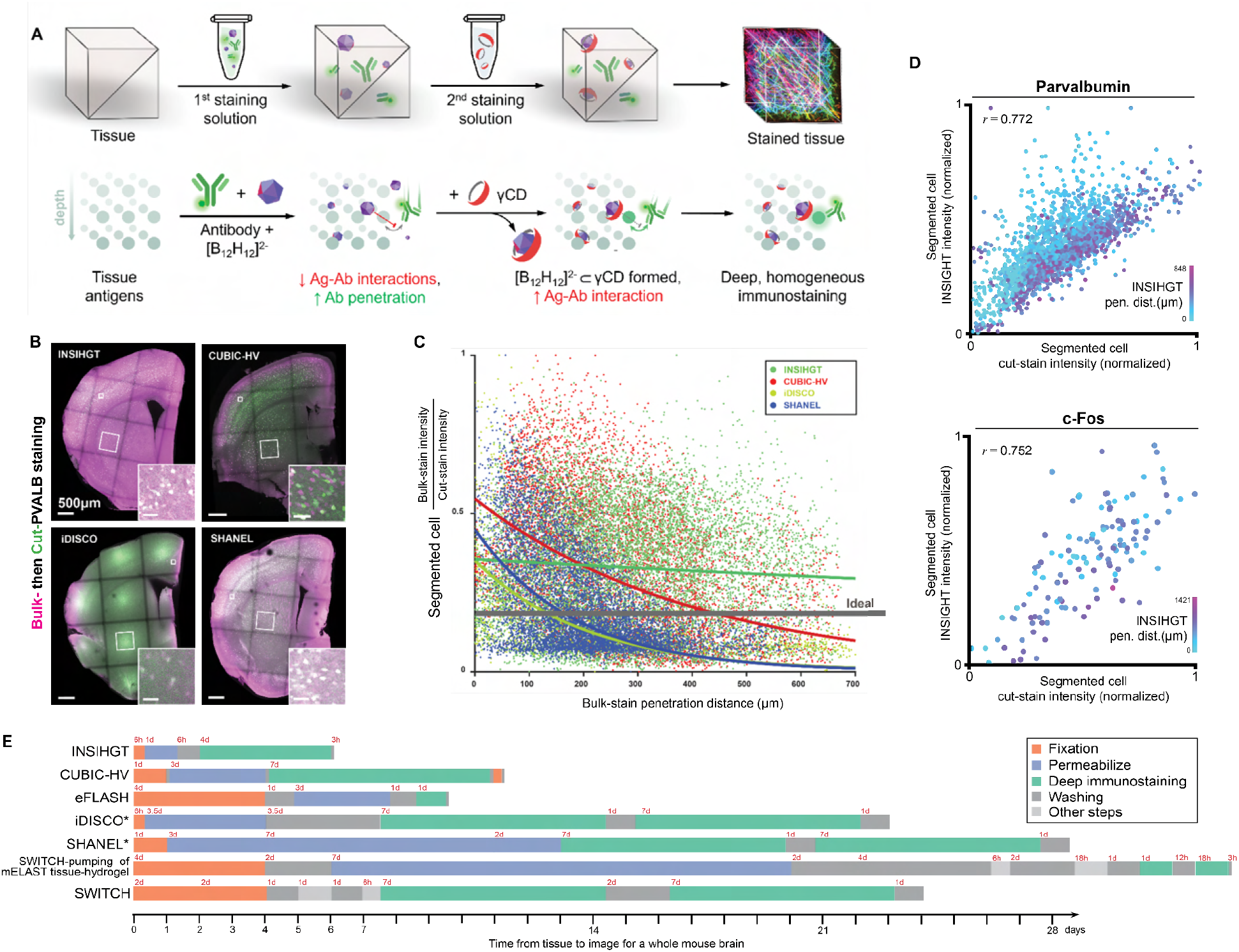
Homogeneous and deep staining with INSIHGT. (**A**) Experimental steps and principle of INSIHGT for immunostaining. Top row: Tissue is infiltrated with antibodies and a weakly coordinating superchaotrope ([B_12_H_12_]^2-^, purple dodecahedron) in the 1st staining solution and then transferred into the 2nd solution containing a complexation agent (CD, red ring). Bottom row: The molecular principles of INSIHGT. Weakly coordinating anion prevents antibody-antigen interactions, removing penetration obstacles. After homogeneous infiltration, subsequent γCD infiltration complexes the [B_12_H_12_]^2-^ ions, allowing deep tissue immunostaining. (**B**) Benchmark results of six buffers used in deep immunostaining. Enlarged views of smaller areas are shown in insets. Parvalbumin (PVALB) immunostaining signals on cut surface: magenta, bulk-staining signal; green: cut-staining PVALB signal (refer to **Fig. S1**). (**C**) Quantification of bulk:cut-staining signal ratio against penetration distance for segmented cells. Each dot represents a cell. Lines are single-term exponential decay regression curves. The signal decay distance constants (*#x03C4;*) are shown in **Table S1**. Hypothetical ideal method performance is shown as a gray line (*#x03C4;*→0+). (**D**) Correlation of INSIHGT signal with reference (cut-staining intensity) signal, illustrating 3D quantitative immunostaining. *r*: Pearson correlation coefficient. (**E**) Timeline illustration for a whole mouse brain processing experiment with the different benchmarked methods (drawn to scale). *indicates methods where in principle the use of secondary antibody Fab fragments can lead to as short as half the immunostaining time.

Notably, after washing, only a negligible effect of [B_12_H_12_]^2-^-treatment will remain within the tissue. This is evident as the cut-staining intensity profile of INSIHGT was similar to that of iDISCO (**Fig. S7**) which has identical tissue pre-processing steps. Upon the addition of 2HPγCD and washing off the so-formed complexes, the penetration enhancement effect was completely abolished. This suggests that [B_12_H_12_]^2-^ and cyclodextrins do not further permeabilize or disrupt the delipidated tissue.

### High-throughput, multiplexed, dense whole organ mapping

After confirming INSIHGT can achieve uniform, deeply penetrating immunostaining, we next applied INSIHGT to address the challenges in whole organ multiplexed immunostaining, where the limited penetration of macromolecular probes hinders the scale, speed, or choice of antigens that can be reliably mapped. Due to the operational simplicity, scaling up the sample size in organ mapping experiments with INSIHGT is straightforward and can be done using multiwell cell culture plates (**Fig. 3A**). For example, we demonstrated our case by mapping 14 mouse kidneys in parallel (**Fig. 3B**) within 6 days of tissue harvesting using a standard 24-well cell culture plate.

**Figure 3.**
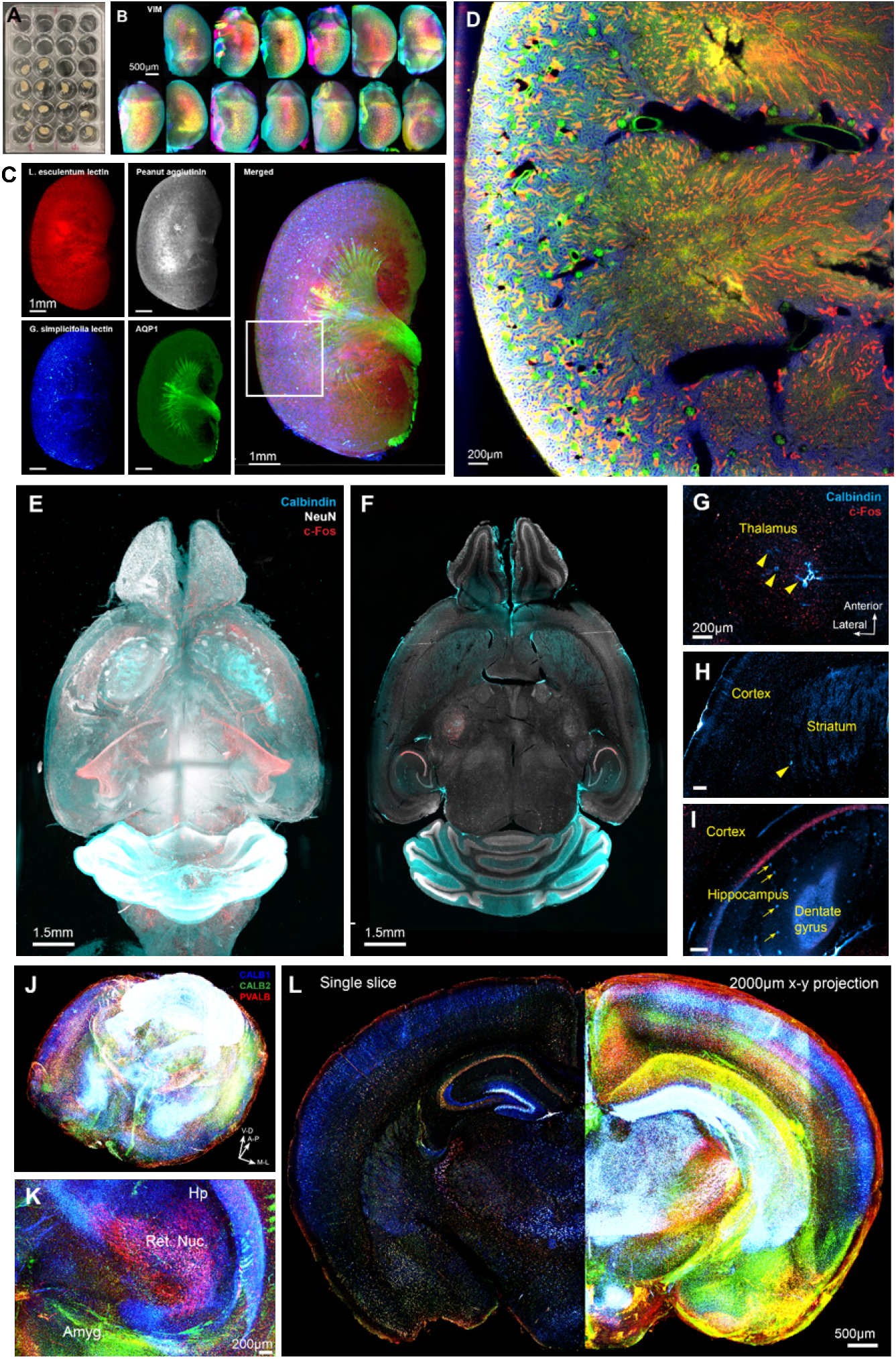
High-throughput whole-organ deep immunostaining with dense mapping using INSIHGT. (**A**) Parallelized sample processing with INSIHGT, exemplified with whole mouse kidneys. (**B**) Whole organ imaging results with parallelized INSIHGT for the samples shown in (**A**), showing vimentin INSIHGT signals color-coded by z-depth. One sample was dropped due to manual errors. (**C-D**) Whole mouse kidney densely multiplexed visualization with *Lycopersicon esculentum* lectin (red), Peanut agglutinin (grey), *Griffonia simplicifolia* lectin I (blue), and AQP-1 (green). (**D**) Enlarged 2D view of the white boxed area in (**C**). (**E-F**) Rendered view of whole mouse brain multiplexed Calbindin, NeuN, and c-Fos mapping of a 3-year-old mouse. (**G-I**) Age-related structural and molecular changes in the thalamus (**G**) and striatum (**H**) with cavitations (indicated by yellow arrowheads), and the hippocampus (**I**) with CALB1-positive deposits (indicated by yellow arrows). (**J-L**) Whole brain multiplexed staining of the calcium binding proteins calbindin (CALB1), calretinin (CALB2), and parvalbumin (PVALB) with 3 days of INSIHGT staining. **(K)** Zoomed in 3D rendering view on the hippocampus (Hp), reticular nucleus of thalamus (Ret. Nuc.) and amygdala (Amyg.). **(L)** A single coronal slice view (left) and a 2mm-thick anteroposterior projection (right) of the same sample.

We then exemplify the capability of INSIHGT to simultaneously map densely expressed targets in whole organs (**Fig. 3C-I**). We first performed multiplexed staining on mouse kidney with 3 days of incubation for *Lycopersicon esculentum* lectin (LEL), Peanut agglutinin (PNA), *Griffonia simplicifolia* lectin (GSL), and AQP-1, which are targets associated with poor probe penetration due to their binding targets’ dense expession (**Fig S2, Fig. 3C-D**). With the use of INSIHGT, the dense tubules and vascular structures can be reliably visualized and traced (**Fig. S8**).

We then proceeded to map the whole brain of a 3-year old mouse at the time of euthanasia. We utilized INSIHGT with 3 days of staining for Calbindin (CALB1), NeuN, and c-Fos, providing cell type and activity information across the aged organ (**Fig. 3E-I**). With whole organ sampling, we identified regions where ageing related changes were prominent, these include cavitations in the bilateral thalamus and striatum (**Fig. 3G-H**), as well as calbindin-positive deposits in the stratum radiatum of hippocampus (**Fig. 3I**). Interestingly, there seems to be an increased c-Fos expression level among the neurons surrounding thalamic cavitations (**Fig. 3G**) which are located deep within the brain tissue and thus cannot be explained by preferential antibody penetration, suggesting these cavitations may affect baseline neuronal activities. Similar 1-step multiplexed mapping of calcium-binding proteins across a whole adult mouse brain can also be performed with 3 days of staining (with a fixed tissue-to-image time of 6 days) (**Fig. 3J-L, Movie S1**). Similarly, whole adult mouse brain mapping and statistics can be obtained for ∼35 million NeuN+ cells, their GABA quantities and c-Fos expression levels using the same protocol (**Fig. S9**), allowing structure, neurotransmitter and activity markers to be analyzed simultaneously.

Overall, INSIHGT overcomes technical, operational, and cost bottlenecks towards accessible organ mapping for every basic molecular biology laboratory, providing rapid workflows to qualitatively evaluate organ-wide structural, molecular, and functional changes in health and disease, regardless of the spatial density of the visualization target.

### Boron cluster-based supramolecular histochemistry as a foundation for spatial multi-omics

With the maturation of single-cell omics technologies, integrating these high-dimensional datasets becomes problematic. Embedding these data in their native 3D spatial contexts is the most biologically informative approach. Hence, we next tested whether our boron cluster supramolecular chemistry allows the retention and detection of multiple classes of biomolecules and their features, based on which 3D spatial multi-omics technologies can be developed.

With identical tissue processing steps and INSIHGT conditions, we tested 357 antibodies and found 323 of them (90.5%) produced the expected immunostaining patterns as manually validated with reference to the human protein atlas and/ or existing literature (**Fig. 4A, Fig. S10-12, Table S2**). This was at least six times the number of compatible antibodies demonstrated by any other deep immunostaining method (**Fig. 4A**), demonstrating the robustness and scalability of INSIHGT. Antigens ranging from small molecules (e.g., neurotransmitters), epigenetic modifications, peptides to proteins and their phosphorylated forms were detectable using INSIHGT (**Fig. 4B-C**). The specificity of immunostaining even allowed the degree of lysine methylations (i.e., mono-, di- and tri-methylation) and the symmetricity of arginine dimethylations to be distinguished from one another (**Fig. 4B-C**). We further tested 21 lectins to detect complex glycosylations, proving that [B_12_H_12_]^2-^ do not sequester divalent metal ions essential for their carbohydrate recognition (**Fig. 4D, Fig. S13**).

**Figure 4.**
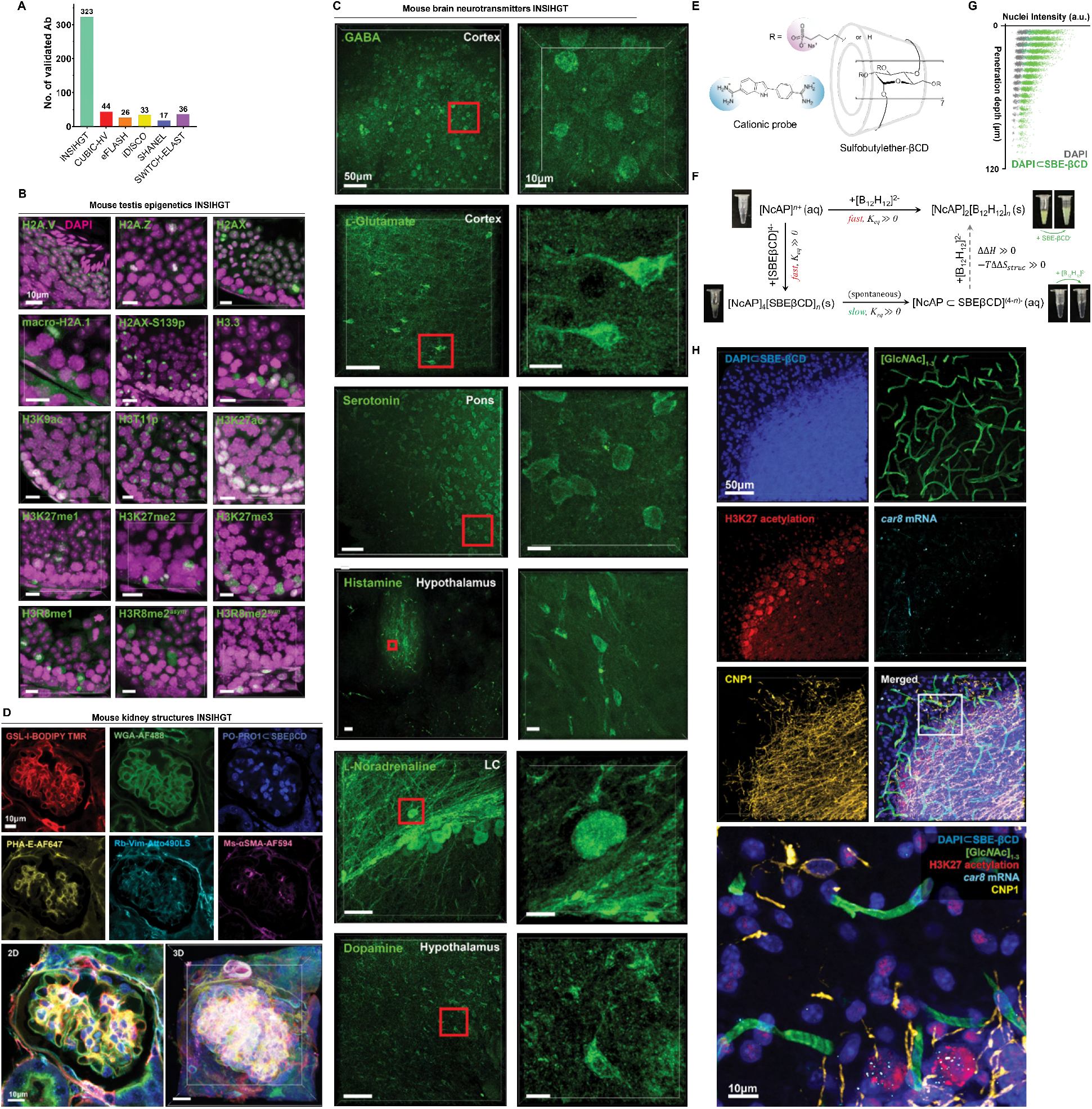
Multi-modality INSIHGT for generality. (**A**) Number of validated antibodies compatible with each benchmarked method. (**B–D**) INSIHGT’s compatibility in revealing the 3D location of various features. (**B**) Epigenetic markers based on histone post-translational modifications and isoforms. (**C**) Neurotransmitters. LC: locus ceruleus. (**D**) Structural features, using one-step multiplexed supramolecular histochemistry, multiplexed supramolecular lectin histochemistry, and the supramolecular dye complex [PO-PRO1⊂SBEβCD]. (**E**) Nucleic acid probe (NAP, DAPI structure shown) complexation by SBEβCD for improved tissue penetration. (**F**) Thermodynamic scheme of NAP’s complexation reaction with SBEβCD. SBEβCD neither redissolves DAPI/[B_12_H_12_]^2-^ precipitates nor precipitates DAPI out from the [DAPI⊂SBEβCD] complex, suggesting kinetic stabilization. (**G**) Quantification of penetration depths of [DAPI⊂SBEβCD] compared to traditional DAPI staining. (**H**) Multimodal 3D molecular phenotyping in a 1mm-thick mouse cerebellum slice for proteins (CNP1), nucleic acids (*car8*, DAPI), epigenetic modifications (H3K27 acetylation), and glycosylations ([Glc*N*ac]_1-3_) with INSIHGT. Imaging was limited to 200 μm due to working distance constraints.

Small molecule dyes such as nucleic acid probes, which are mostly positively charged, presents a separate challenge as they precipitate with *closo*-dodecaborates, forming [probe]^n+^/[B_12_H_12_]^2-^ precipitates when co-applied with INSIHGT. We found size-matched and charge-complementing cyclodextrin derivatives as cost-effective supramolecular host agents for non-destructive deep tissue penetration and preventing precipitation. For example, sulfobutylether-βCD (SBEβCD) (**Fig. 4E**) can react with nucleic acid probes to form [probe⊂SBEβCD], which exhibits penetration enhancement during INSIHGT (**Fig. 4F, G**) without precipitation problems.

We also performed RNA integrity number (RIN) and whole genome DNA extraction analyses on INSIHGT-treated samples (**Fig. S14**). We found each step of the INSIHGT protocol did not result in a significant decrease in RNA integrity number (RIN) (**Fig. S14A**). The total RNA extracted after undergoing the whole INSIHGT protocol has an RIN of 7.2, compared with an RIN of 9 from a treatment-naive control sample. For whole genome DNA, both control and INSIHGT-protocol-treated samples have similar sample integrity and total DNA yield per mm^3^ sample (14.6μg versus 10.12μg, **Fig. S14B**). Hence, unsurprisingly, we found single-molecule fluorescent in situ hybridization (FISH) is also applicable for co-detection of protein antigens and RNAs with INSIHGT. Combining all the above probes, simultaneous 3D visualization of protein antigens, RNA transcripts, protein glycosylations, epigenetic modifications, and nuclear DNA is possible using a mixed supramolecular system in conventionally formalin-fixed intact tissue (**Fig. 4H**). Taken together, our results suggest *in situ* boron cluster supramolecular histochemistry can form the foundation for volumetric spatial multi-omics method development.

### Centimeter-scale 3D histochemistry by isolated diffusional propagation

Since antibody penetration remains the most challenging obstacle, we focus the remainder of our investigation on larger-scale 3D immunophenotyping. We thus applied INSIHGT to visualize centimeter-scale human brain samples, without using any external force fields to drive the penetration of macromolecular probes. These large, pigmented samples were sliced in the middle of the tissues’ smallest dimensions to allow imaging of the deepest areas with tiling confocal microscopy. We show that INSIHGT can process a 1.5 cm × 1.5 cm × 3 cm human cortex block for parvalbumin (PV) (**Fig. 5A–C**), with excellent homogeneity and demonstration of parvalbumin neurons predominantly in layer 4 of the human cortex.

**Figure 5.**
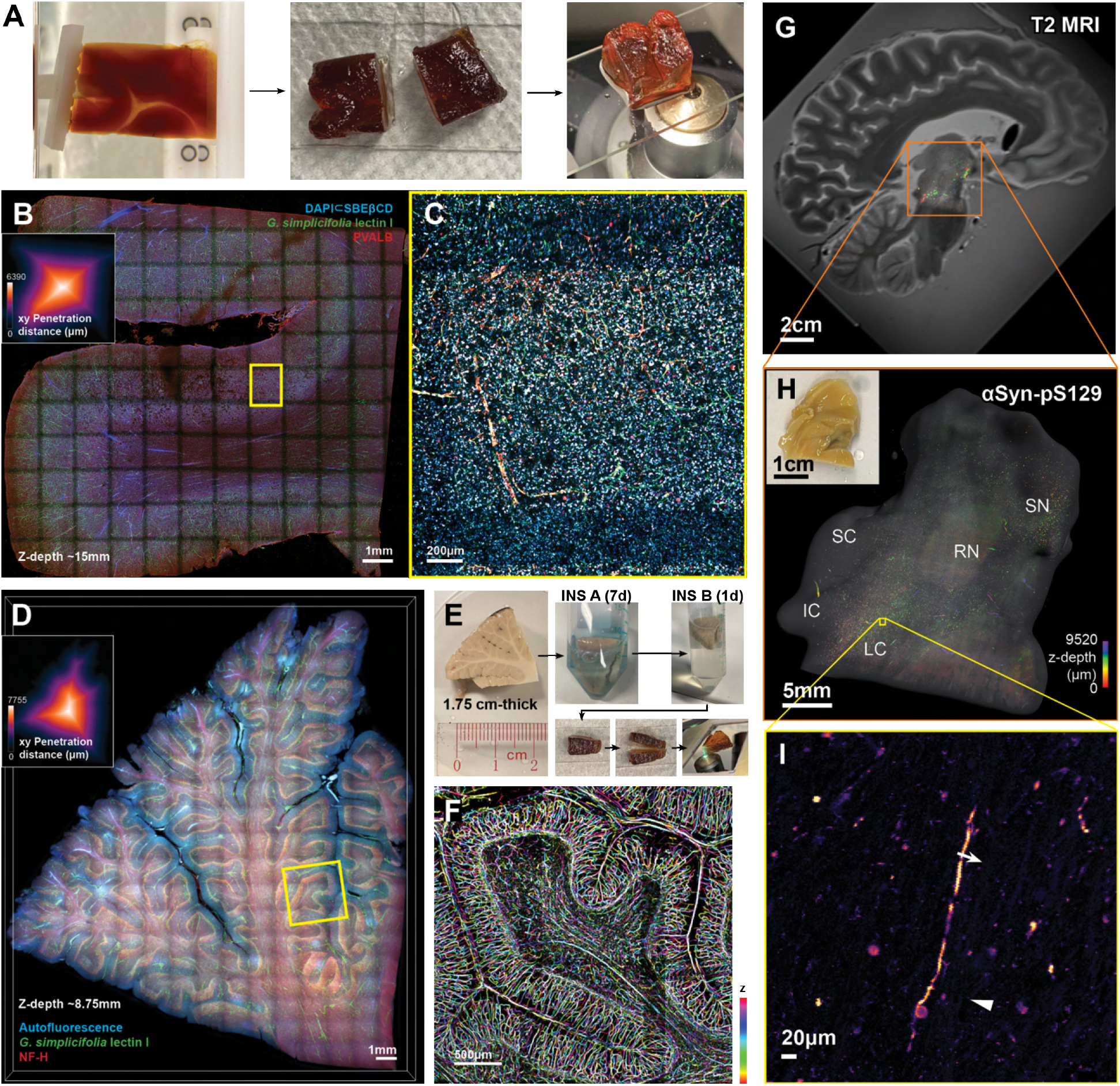
Centimeter-scale INSIHGT. (**A**) A human cortex block (1.5 cm × 1.5 cm × 3 cm), processed with INSIHGT and cut in the middle for confocal imaging to confirm penetration depth. (**B**) Confocal tiled image of the cut face from (**A**), stained with [DAPI⊂SBEβCD], *G. simplicifolia* lectin I (GSL-I), and for parvalbumin (PVALB). Inset: penetration depth over the imaging surface. (**C**) Enlarged view of the yellow boxed area in (**B**). (**D**) A human cerebellum tissue block (1.75 cm × 2.0 cm × 2.3 cm) processed with INSIHGT, stained with GSL-I lectin and NF-H. Inset: penetration depth over the imaging surface. (**E**) Illustrated tissue processing steps: tissue is stained for seven days in INSIHGT buffer A (with [B_12_H_12_]^2-^), washed in INSIHGT buffer B (with 2HPγCD), and sliced perpendicular to the thinnest dimension at the midpoint. (**F**) Enlarged view of the yellow boxed area in (**D**).

We then scaled INSIHGT to a 1.75 cm × 2.0 cm × 2.2 cm human cerebellum block for blood vessels (using *Griffonia simplicifolia lectin I, GSL-I*) (**Fig. 5D–F**), again revealing excellent homogeneity with no decay of signal across the centimeter of penetration depth. This shows that the use of boron cluster-based host-guest chemistry remains applicable for highly complex environments at the centimetre scale. The results further show that macromolecular transport within a dense biological matrix can remain unrestricted in a non-denaturing manner by globally adjusting inter-biomolecular interactions.

We further applied INSIHGT to a 1.0 cm × 1.4 cm × 1.4 cm human brainstem with dementia with Lewy bodies (DLB) for phosphorylated alpha-synuclein at serine 129 (αSyn-pS129) (**Fig. 5G-I, Fig. S15**). The large scale of imaging enabled registration and hence correlation with mesoscale imaging modalities such as magnetic resonance imaging (MRI) (**Fig. 5G, Movie S2**). With this, we confirmed the localization of Lewy body pathologies to the subcerulean nuclei ^25^ and substantia nigra, in keeping with the prominent rapid eye movement sleep behavior disorder (RBD) symptoms of this patient. Such a radio-histopathology approach towards mapping neurodegenerative aggregates may be useful for the development of gross imaging-based diagnostics for predicting underlying neurodegeneration. Overall, the capability of INSIHGT in achieving centimeter-sized tissue staining bridges the microscopic and mesoscopic imaging modalities, providing a general approach to correlative magnetic resonance-molecular imaging.

### Volumetric spatial morpho-proteomic cartography for cell type identification and neuropeptide proximity analysis

We next extended along the molecular dimension on conventionally fixed tissues, where highly multiplexed immunostaining-based molecular profiling in 3D had not been accomplished previously. A single round of INSIHGT-based indirect immunofluorescence plus lectin histochemistry can simultaneously map up to 6 antigens (**Fig. S16**), tolerating a total protein concentration at >0.5 μg/μl in the staining buffer, and is limited only by spectral overlap and species compatibility. To achieve higher multiplexing, antibodies can be stripped off with 0.1 M sodium sulfite in the [B_12_H_12_]^2-^-containing buffer after overnight incubation at 37°C (**Fig. 6A, Fig. S17**). Since [B_12_H_12_]^2-^ does not significantly disrupt intramolecular and intermolecular noncovalent protein interactions, the approach can be directly applied to routine formaldehyde-fixed tissues, we observed no tissue damage and little distortion, obviating the need for additional or specialist fixation methods.

**Figure 6.**
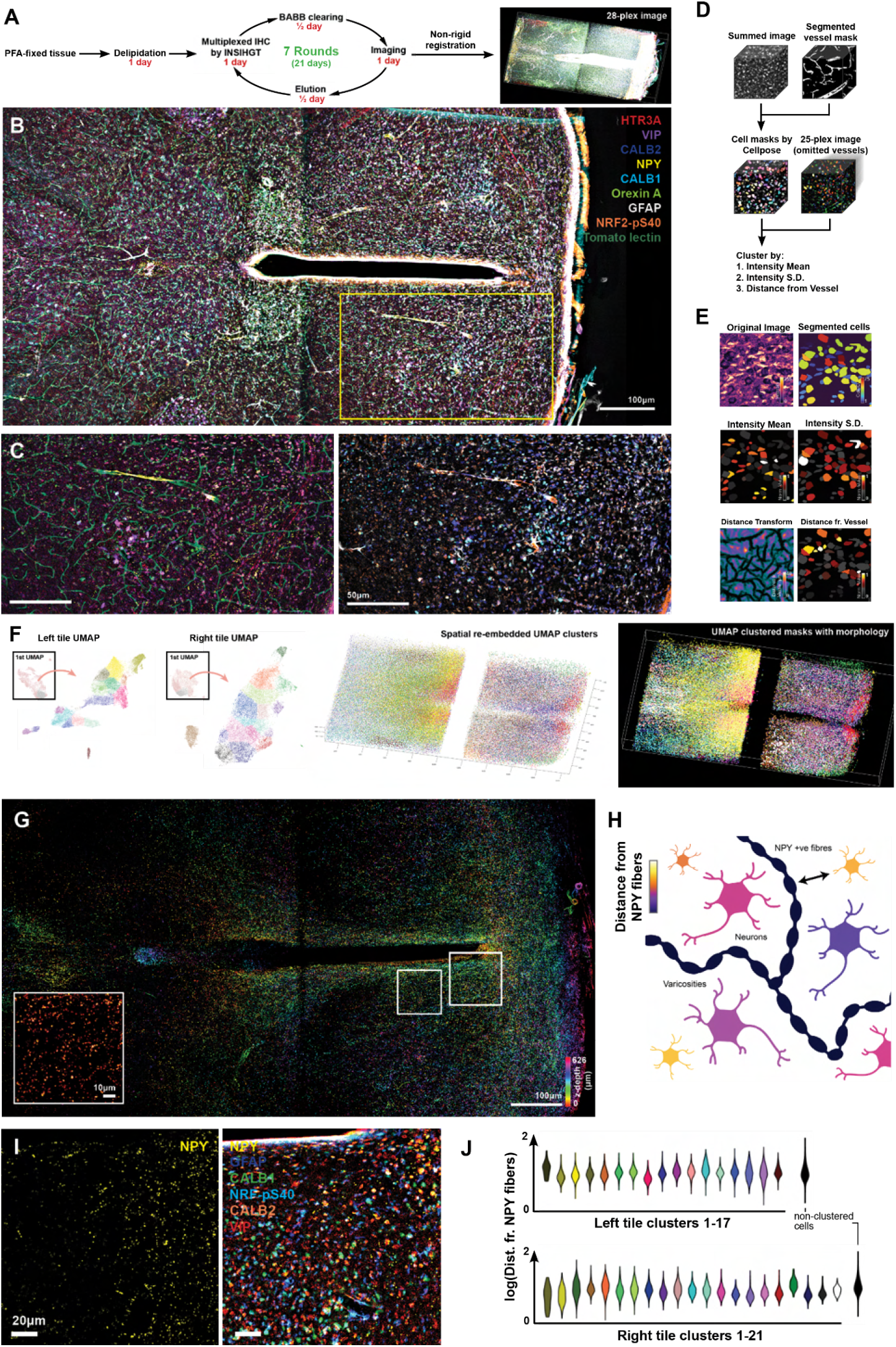
Multi-round multiplexed INSIHGT. (**A**) Schematics of the processing steps for a 1mm-thick mouse hypothalamus sample. (**B**) A selection of multi-round immunostaining signals (for nine targets) displayed for the multi-round multiplexed INSIHGT-processed sample. (**C**) Enlarged view of the yellow boxed area in (**B**) with complementary panel of markers. (**D**) Schematics of the image analysis process. (**E**) Illustrated results of a segmented image subset. The images below show corresponding cell segmentation and quantification results. (**F**) Left panel: Nested UMAP embedding of all segmented cells from both tiles of the image stack. Middle panel: The spatial locations of the different color-coded clusters. Right panel: Similar to the middle panel but with cellular morphology. (**G**) Color-coded *z*-projection of neuropeptide Y (NPY) staining signals. A higher magnification view for the left white boxed area is shown in the inset. (**H**) Schematic representation of distance measurement from NPY fibers to cell bodies via distance transformation. (**I**) NPY signal in a periventricular region (right white boxed area in (**G**)) is shown in the left panel, and with selected markers staining in the right panel. (**J**) Quantification of distance from NPY-expressing fibers for each cell type shown in violin plots, based on the 3D spatial locations of somas and NPY fibers.

We exemplified this approach by mapping 28 neuronal marker expression levels in a 2 mm-thick hypothalamus slice over 7 imaging rounds (**Fig. 6A–C**). With each iterative round taking 48 hours (including imaging, retrieval and elution), the whole manual process from tissue preparation to the 28-plex image took 16 days. After registration and segmentation using Cellpose 2.0 ^26^ (**Fig. 6D, E**), we obtained 192,075 cells and their differentially expressed proteins (DEPs) based on immunostaining signals. Uniform manifold approximation and projection (UMAP) analysis of a subset of 84,139 cells based on 50 markers (omitting 3 blood vessels channels and separately accounting for signal mean intensity and standard deviations, **Fig. 6F, Fig. S18-19**) plus their distance to the nearest vessels revealed 42 cell type clusters, allowing their 3D spatial interrelationships to be determined (**Fig. S20**).

3D spatial proteomics can blend morphology and molecular information to infer functional relationships, which is inaccessible to spatial transcriptomics or single-cell multi-omics. Recent characterizations of neuronal network activities based on the diffusional spread of neuropeptides highlight the need for 3D spatial mapping of protein antigens. To obtain these morphological-molecular relationships using INSIHGT, we segmented the neuropeptide Y (NPY)-positive fibers and computed the 3D distance to each UMAP-clustered cell types’ somatic membranes (**Fig. 6G–J**). While most clusters have a similar distance from NPY fibers, certain clustered cells (notably right tile clusters 1 and 2) are more proximally associated with NPY fibers, suggesting these cell clusters are differentially modulated by NPY when isotropic diffusion is assumed in the local brain parenchyma. Nonetheless, our dataset and analysis demonstrated it is possible to estimate the likely modulatory influence for a given cell-neuropeptide pair, providing a new approach to discovering novel neuronal dynamics paradigms.

### Fine-scale 3D imaging reveals novel intercellular contacts traversing the Bowman space in mouse kidneys

We found that the process of INSIHGT from fixation to completion preserve delicate structures such as free-hanging filaments and podia, enabling fine-scale analysis of compact structures such as the renal glomeruli. We found novel intercellular contacts traversing the Bowman space, which to the best of our knowledge has not been previously described even in extensive reviews on parietal epithelial cells (PECs) ^27,28^ and electron microscopy images ^29–31^ (**Fig. 7A, B**). These filaments are mostly originated from the podocytic surface, although some were also seen to emerge from PECs. They were numerous and found around the glomerular globe (**Fig. 7C**), with varied in their length, distance from each other, and morphologies (**Fig. 7D, Fig S21**).

**Figure 7.**
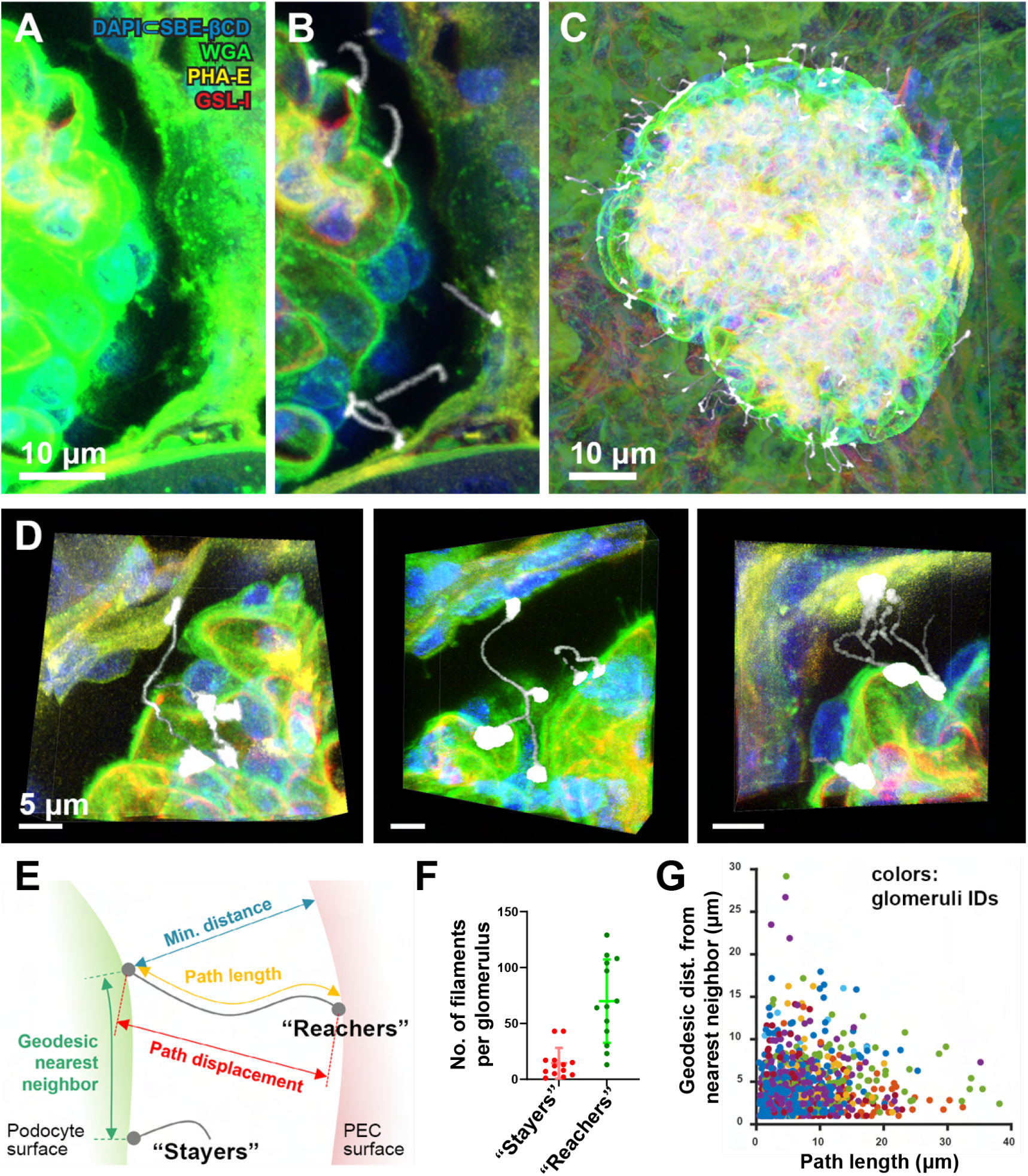
INSIHGT reveals novel intercellular filaments traversing the Bowman space in mouse kidneys. (**A**) Original image of multiplexed image of whole mouse glomerulus with full Bowman capsule. WGA: Wheat germ agglutinin, PHA-E: *Phaseolus vulgaris* hemagglutinin, GSL-I: *Griffonia simplicifolia* lectin I. (**B**) Segmentation of microfilaments with en bloc preservation of native morphologies and spatial relationships in 3D Euclidean space. (**C**) Global representation on the 3D spatial distribution of microfilaments across the entire Bowman space. (**D**) Distinct and protean morphologies of the podocyte-to-parietal epithelial cell (PEC) microfilaments. (**E**) Schematic representation of reachers (contacting PEC surface) and stayers (remaining in the Bowman space) originating from podocyte surfaces with the related physical parameters. (**F**) Descriptive statistics of microfilament subtypes per analyzed glomeruli. N = 14 glomeruli analyzed across 4 mice. (**G**) Correlation between the path length of each microfilament and the geodesic distance between its podocytic attachment point and its nearest neighbor. The data points were color-coded based on their glomerulus of origin in the dataset.

We classified these podocyte-to-PEC microfilaments into “reachers” and “stayers”, depending on whether they reached the PEC surface or not (**Fig. 7E**). Microfilaments of the reachers-type were more numerous than the stayers-type per glomerulus (**Fig. 7F**). Visually, we noted the emergence of these filaments tended to cluster together, especially for the reachers-type. To quantify such spatial clustering, we calculated the glomerular surface geodesic distances between the podocytic attachment points for each microfilament, which showed an inverse relationship with their path lengths (**Fig. 7G**), and reachers-type filament are geodesically located nearer to each other than the stayers type (**Fig. S22**). This suggests that the emergence of long, projecting microfilaments that reach across the Bowman space is localized on a few hotspots of the glomerular surface. Whether these hotspots of long-reaching microfilaments are driven by signals originated from the podocyte, the glomerular environment underneath, or the overarching PECs may reveal previously unsuspected podocyte physiological responses within their microenvironments.

### Sparsely distributed neurofilament inclusions unique to the human cerebellum

We next completely mapped a 3 mm-thick (post-dehydration dimensions) human cerebellar folium for NF-H, GFAP and blood vessels (**Fig 8A, Fig. S23-24, Movie S3**), with preserved details down to the Bergmann glia fibers, perivascular astrocytic endfeet, and Purkinje cell axons that makes the amenable to 3D orientation analysis and visualization (**Fig. 8B-D, Fig. S23-24**). The detailed visualization of filamentous structures throughout the 3 mm-thickness is in stark contrast to our previous attempts with similar specimens employing various methods, which showed weak NF-H signal in cerebellar sulci and barely visible GFAP signal in cerebellar white matter due to poor antibody penetration (data not shown).

**Figure 8.**
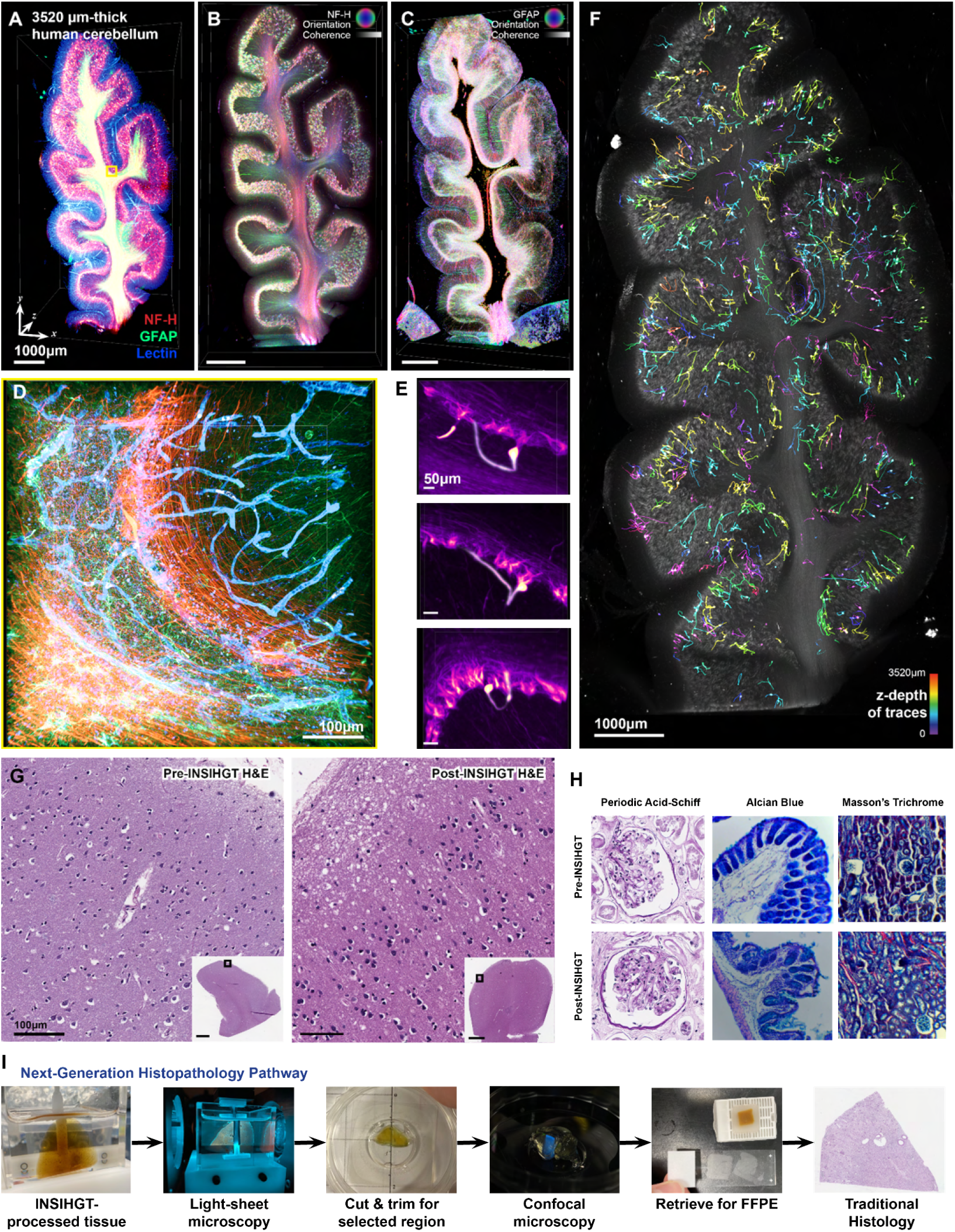
INSIHGT enables non-destructive characterization and analysis of human clinical samples. (**A**) A 3.5mm-thick human cerebellum triplex-stained for glial filaments (GFAP), neurofilament (NF-H) and blood vessels (*G. simplicifolia* lectin I). (**B-C**) Orientation and coherence (or fractional anisotropy) visualization of neurofilament (**B**, NF-H) and glial filament (**C**, GFAP) via structure tensor analysis. (**D**) Enlarged view of the boxed area in (**A**) via post-hoc confocal microscopy. (**E**) Prototypical morphology of human cerebellar neurofilament inclusions, where their extensions may loop back to the Purkinje layer and occasionally to another inclusion body (lowest image). (**F**) Overview of the 1078 manually traced neurofilament inclusions across the cerebellar sample, color-coded by z-depth. (**G-H**) Traditional 2D histology with special stains on pre-INSIHGT and post-INSIHGT processed samples. (**G**) H&E staining of human brain, (**H**) Left to right: Periodic acid-Schiff (PAS), Alcian blue, and Masson’s trichrome staining of human kidney, mouse colon, and mouse kidney sections respectively. (**I**) The Next-Generation Histopathology Pathway. INSIHGT is compatible to traditional histological pipelines, empowering a multi-pronged approach to maximizing the information extracted from clinical samples.

We discovered sparsely distributed NF-H-intense inclusions that are easily missed in 2D sectioning and thus remain poorly characterised. We manually traced and identified 1,078 inclusions throughout the entire imaged volume (**Fig. 8E-F**), where they were found in all of the three basic layers of the cerebellar cortex. A typical morphology of one type of these inclusion is a single bright globular inclusion at the sub-Purkinje layer radial location, with an elongated thick fiber extension that coils back and project to the adjacent molecular layer (**Fig. 8E**). However, much more protean morphologies also exist (**Fig. 8E-F, Fig S25**). To capture the morphological and spatial diversities of these inclusions, we obtained their spatial-morphometric statistics (**Fig. S25A**), followed by principal component analysis of the compiled morphometrics such as Sholl analysis and Horton-Strahler number. The results reveal most of these inclusions to be morphologically homogeneous with variations explained largely by their path lengths, with a small subset characterized by much higher branching of the NF-H-intense filaments (**Fig. S25B**).

### INSIHGT bridges the gap between 3D histology and traditional 2D pathology in current clinical practice

The bio-orthogonal nature of the INSIHGT chemical system underlies its non-destructiveness. To highlight the clinical impact of INSIHGT in addition to 3D imaging of human samples, we found that INSIHGT-processed samples can be retrieved and processed as naïve tissues for traditional 2D histology via paraffin wax embedding and sectioning. Notably, staining qualities of routine hematoxylin and eosin (H&E) and various special stains on the post-INSIHGT processed slides were indistinguishable from the pre-INSIHGT processed slides even by a senior pathologist (**Fig. 8G-H**). In addition to not interfering with downstream clinical processes, the preserved quality of special staining allows for multi-modal cross-validation of 3D fluorescent imaging findings, making INSIHGT the ideal platform choice for next-generation histopathology (**Fig. 8I**). We envision with INSIHGT, quantitative 3D information within clinical specimens can be maximally extracted with high authenticity, and its fast speed promises compatibility with current clinical workflows and constraints, allowing digital pathology and precision medicine to benefit from 3D analysis.

## Discussion

The convergence of multiple technological advances has paved the way for the acquisition of large-scale molecular phenotyping datasets at single-cell resolution, most notably single-cell transcriptomics ^32^. With a large number of newly discovered cell states, the quest to extend towards spatially resolved cell phenotyping based on translated and post-translationally expressed biomolecular signatures is paramount to understanding their structural and functional properties in biology ^33^.

Scalable, high-resolution 3D tissue mapping provides a powerful approach to further our understanding of these newly identified cell types. Clinically, 3D histology has been shown to improve diagnosis in bladder cancer ^34^, predict biochemical recurrence in prostate cancer ^35^, and evaluate response to chemotherapy in ovarian carcinoma ^34^, By sampling across whole intact samples, 3D histology can deliver unbiased, quantitative, ground-truth data on the spatial distributions of molecules and cell types in their native tissue contexts ^36^. However, 3D tissue imaging is yet to be widely adopted despite the increasing accessibility of tissue clearing, optical sectioning microscopy, and coding-free image processing software. This is in large part due to the limited penetration of probes that plague the field regardless of the combinations of these technologies employed ^6,10^, yielding variable, surface-biased data with questionable representativeness. Creative approaches have provided solutions to the penetration problem but are limited in their scalability and accessibility ^6^.

Constrained by the requirements of non-advective approaches and compatibility with off-the-shelf reagents, the development of INSIHGT involved re-examining biomolecular transport and protein stability from the first principles, which led us to identify weakly coordinating superchaotrope and its chemical activity modulation by in situ host-guest reactions to implement our theoretical formulation. With the use of *closo*-dodecaborate and cyclodextrin as an additive in PBS, we solved the bottleneck of 3D histology by providing a cost-efficient, scalable, and affordable approach to quantitatively map multiple molecules in centimeter-sized tissues. With an equivalent tissue processing pipeline to iDISCO ^17^, INSIHGT shares the same affordability and scalability while providing much faster processing and greatly improved image quality, due to enhanced antibody penetration depth and homogeneity. Mapping tissue blocks simultaneously in multi-well dishes is easily accomplished in any basic molecular biology laboratory. Such simplicity in operation makes it highly accessible and automatable, as it requires no specialized equipment or skills. Furthermore, cocktails of off-the-shelf antibodies can be directly added to PBS supplemented with [B_12_H_12_]^2-^. Finally, we note that both [B_12_H_12_]^2-^ salts and cyclodextrins are non-hazardous and stable indefinitely at ambient temperatures ^16^.

With the affordability and accessibility of INSIHGT, we anticipate its diverse applications in 2D and 3D histology applications. Meanwhile, boron cluster-based supramolecular histochemistry can form the backbone for 3D spatial molecular-structural-functional profiling methods and studies, as well as atlas mapping efforts. The high-depth, quantitative readout of well-preserved tissue biomolecules offered by INSIHGT forms the foundation for multiplexed, multi-modal, and multi-scale 3D spatial biology. By making non-destructive 3D tissue molecular probing accessible, INSIHGT can empower researchers to bridge molecular-structural inferences from subcellular to the organ-wide level, even up to clinical radiological imaging scales for radio-histopathological correlations. Finally, the compatibility of INSIHGT with downstream traditional 2D histology methods indicates its non-interference with subsequent clinical decision-making. This paves the way for the translation and development of 3D histology-based tissue diagnostics, promising rapid and accurate generation of groundtruth data across entire tissue specimens.

We recognize that INSIHGT still has room for further improvements. Immunostaining penetration homogeneities for larger tissues and denser antigens can be further enhanced, and the penetration of small molecule dyes and lectins were still suboptimal for millimeter-scale tissues. In multi-round immunostaining, we noticed that the staining specificity and sensitivity deteriorated with each round of antibody elution with sulphite or β-mercaptoethanol, calling for a better 3D immunostaining elution method. Alternatively, hyperspectral imaging ^37^, nonlinear optics ^38^, time-resolved fluorescence techniques ^39^, and same-species antibody multiplexing ^40^ could be explored to extend the multiplexing capabilities of INSIHGT.

Our discovery of boron clusters’ capabilities to solubilize proteins globally in a titratable manner, combined with their bio-orthogonal removal with supramolecular click chemistry, can reach beyond histology applications. Given the surprisingly robust performance of INSIHGT in complex tissue environments, we envision they can be applied in simpler *in vitro* settings to control intermolecular interactions – particularly when involving proteins – in a spatiotemporally precise manner.

## Acknowledgements

We would like to express our deepest gratitude to the tissue donor and his family for their generosity and wisdom in supporting scientific studies. We thank William Wu for access to confocal microscopy; Cathy Shuk Ling Chan for manual data annotation and analysis; Ka Wai Chan for administrative support to the project. Figure 1 was partly created with BioRender.com.

## Author Contributions

Conceptualization: HML. Methodology: HML, CNY, JTSH, RAAC, TCYW, BTYW, NKNC, LZ, EPLT. Investigation: HML, CNY, JTSH, RAAC, TCYW, BH, BTYW, NKNC, LZ, EPLT, YT, JJXL, YKW. Visualization: HML, CNY, JTSH, RAAC. Funding acquisition: YKW, HML. Project administration: HML. Supervision: HML. Writing: HML, CNY.

## Data and Materials Availability

Additional experimental data and the custom codes used in this study will be made publicly available upon the acceptance of the manuscript.

## Methods

### Materials

#### Chemicals and reagents

The antibodies utilized in this study were listed in **Table S2**. Secondary Fab fragments or nanobodies were acquired from Jackson ImmunoResearch or Synaptic Systems, and all lectins were sourced from VectorLabs. Conjugation of secondary antibodies and lectins with fluorophores was achieved through *N*-hydroxysuccinimidyl (NHS) chemistry. The process was conducted at room temperature for a duration exceeding 16 hours at antibody concentrations >3mg/ml, using a ten-fold molar excess of the reactive dye-NHS ester. Dodecahydro-*closo*-dodecaborate salts and other boron cluster compounds were procured from Katchem, while cyclodextrin derivatives were obtained from Cyclolab, Cyclodextrin Shop, or Sigma Aldrich.

We noticed occasionally the chemicals involved in the INSIHGT process requires purification. Specifically, for Na_2_[B_12_H_12_], if insoluble flakes were noticed after dissolution in PBS, the solution was then acidified to pH 1 with concentrated hydrochloric acid, extracted with diethyl ether (Sigma Aldrich), and the organic solvent removed and distilled off with a warm water bath. The residual H_2_B_12_H_12_ was then dissolved in minimal amount of water, neutralized with 1M Na_2_CO_3_ solution until pH 7 is reached with no further evanescence. The solution was then concentrated by distillation under vacuum and dried in an oven.

For 2-hydroxypropyl-γ-cyclodextrin and sulfobutylether-β-cyclodextrin, if insoluble specks or dusts were noticed after dissolution in PBS, the solution was vacuum filtered through 0.22μm hydrophilic cellulose membrane filters (GSWP14250) using a Buchner funnel before use. A slight brownish-yellow discoloration of the resulting solution would not interfering with the INSIHGT results.

For benzyl benzoate, if the solution is yellowish (possibly due to the impurities of fluorenone present), the solvent is poured into a metal bowl or glass crystallization dish and refrigerated to 4^०^C until crystallization begins. If no crystallization occurs, a small crystal seed of benzyl benzoate obtained by freezing the solvent at -20^०^C in a microcentrifuge tube can be put into the cooled solvent to kick start the process. The crystals were then collected by vacuum filtration with air continuously drawn at room temperature until the crystals are white, which were warmed to 37^०^C to result in clear, colourless benzyl benzoate. If the resulting colourless benzyl benzoate is cloudy, 3Å molecular sieves were added to the solvent to absorb the admixed water from condensation, before filtering off to result in a clear colourless benzyl benzoate. This purified benzyl benzoate is ready for constituting BABB clearing solution for imaging.

#### Human and animal tissues

All experimental procedures were approved by the Animal Research Ethics Committee of the Chinese University of Hong Kong (CUHK) and were performed in accordance with the Guide for the Care and Use of Laboratory Animals. Adult male C57BL/6 were utilized. These mice were housed in a controlled environment (22-23°C) with a 12-hour light-dark cycle, provided by the Laboratory Animal Service Center of CUHK. Unrestricted access to a standard mouse diet and water was ensured, and the environment was maintained at <70% relative humidity. Tissues were perfusion formaldehyde-fixed and collected by post-mortem dissection. In the case of immunostaining for neurotransmitters where Immusmol antibodies were used, the tissues were perfusion-fixed with the STAINperfect™ immunostaining kit A (Immusmol) with the antibody staining steps replaced with those in our INSIHGT method.

For human tissues, brain and kidney tissues were donated post-mortem by a patient were used in this study. Prior ethics approvals have been obtained and approved by the Institutional Review Board with consent from the donor and his family. Human dissection was performed by an anatomist (HML) after perfusion fixation with 4% paraformaldehyde via the femoral artery. The post-mortem delay to fixation and tissue harvesting was 4 weeks at -18°C refrigeration.

### Methods

#### Screening deep staining approaches with in situ antibody recovery

4% PFA-fixed, 1mm-thick mouse cerebellum slices, 0.5μg anti-parvalbumin antibody (Invitrogen, PA1-933) and 0.5μg AlexaFluor 647-labelled Fab fragments of Donkey anti-Rabbit antibody (Jackson Immunoresearch 711-607-003) were used in this experiment to develop our method. Co-incubation of the secondary Fab fragment and primary antibody was utilized for 1-step immunostaining. All stainings were performed with an overnight immunostaining first stage at room temperature (unless specified otherwise) in various buffers, with subsequent recovery secondary stage at room temperature (unless specified otherwise) in various buffers, as detailed for each strategy below. The tissues were then washed in 1x PBSN, dehydrated with graded methanol, and cleared in BABB, before proceeding to imaging with confocal microscopy.

For the SDS/αCD system, immunostaining was performed in a solution consisting of 10mM sodium dodecylsulphate (SDS) in 1xPBS, while recovery was performed with a solution consisting of 10mM αCD in 1x PBS.

For the GnCl/GroEL+GroES system, immunostaining was performed in solution consisting of 6M guanidinium chloride in 1x PBS, while recovery was performed with GroEL+GroES refolding buffer, consisting of 0.5μM GroEL (MCLabs GEL-100), 1μM GroES (MCLabs GES-100), 2.5mM adenosine triphosphate (ATP), 20mM Tris base, 300mM NaCl, 10mM MgSO_4_, 10mM KCl, 1mM tris(2-carboxylethyl)phosphine hydrochloride (TCEP), 10% glycerol, with pH adjusted to 7.9 ^41^.

For the SDC/βCD system, immunostaining was performed in a solution consisting of 15mM sodium deoxycholate (SDC) with 240mM Tris base, 360mM CAPS (*N*-cyclohexyl-3-aminopropanesulfonic acid), with pH adjusted to 8, while recovery was performed with a solution consisting of 15mM βCD with 240mM Tris base, 360mM CAPS, with pH adjusted to 8.

For the Na_2_[B_12_H_12_]/γCD system, immunostaining was performed in a solution consisting of 0.1M Na_2_[B_12_H_12_] in 1x PBS, while recovery was performed in a solution consisting of 0.1M γCD in 1x PBS.

#### Benchmarking experiments

We designed a stringent benchmarking scheme for quantitative evaluation of antibody penetration depth and signal homogeneity across depth for comparison across existing deep immunostaining methods, based on our previously described principles (**Fig S1A**) ^6^. The benchmarking experiment is carried out in two parts, the first part using a whole mouse hemisphere stained in bulk with anti-Parvalbumin (PV) antibodies with excess AlexaFluor 647-conjugated secondary Fab fragments - termed bulk-staining - after which the tissue is cut coronally at defined locations using a brain matrix and re-stained with anti-PV antibodies and AlexaFluor 488-conjugated secondary Fab fragments - termed cut-staining (**Fig S1A**). Hence, signals from bulk-staining can be distinguished easily from cut-staining and reveal different penetration depths of the two-staged immunostaining. We tested different deep immunostaining methods in the bulk-staining stage of the experiments, while the cut-staining was performed in 1× PBS with 0.1% Tween-20 as a conventional immunostaining buffer.

All benchmarking samples were perfusion-fixed with 4% paraformaldehyde (PFA) in 1× PBS followed by post-fixation in 4% PFA overnight at 4°C, except for SHIELD and mELAST samples where the SHIELD protocol was used. In addition, the final RI matching where the benzyl alcohol/benzyl benzoate (BABB) clearing method was universally employed to standardize the changes in tissue volumes and hence penetration distance adjustments. The standardized optical clearing avoids the variability in fluorescent quenching and tissue shrinkage/ expansion introduced by different RI matching agents. For bulk-staining during our benchmarking experiment, we followed the published protocols except for eFLASH and mELAST due to the lack of specialized in-house equipment.

For eFLASH ^12^, we stained the SHIELDed and SDS-delipidated tissue in the alkaline sodium deoxycholate buffer (240mM Tris, 160mM CAPS, 20% w/v D-sorbitol, 0.9% w/v sodium deoxycholate) and titrated-in acid-adjusting booster buffer (20% w/v D-sorbitol and 60mM boric acid) hourly over 24 hours to achieve a -0.1 ± 0.1 pH/hr adjustment rate. The tissue was then washed with 1× PBSTN (1× PBS, 1% v/v Triton X-100 and 0.02% w/v NaN_3_) 2 times 3 hours each before imaging.

For mELAST ^7,13,14^, we stained the SHIELDed and SDS-delipidated tissue with the antibody and Fab fragments in 0.2×PBSNaCh (0.2× PBS, 5% w/v NaCh and 0.02% w/v NaN_3_, 5% v/v normal donkey serum) first for 1 day at 37°C without embedding the SHIELDed tissue in elastic gel nor compression/stretching, followed by adding Triton X-100 to a final concentration of ∼5% and incubated for 1 more day. The tissue was then washed with 1× PBSTN 2 times 3 hours each before imaging.

For CUBIC HistoVision ^8^ and iDISCO ^17^, the tissue was processed and stained as previously described ^9^.

For SHANEL ^42^, the tissue was first delipidated with CHAPS/NMDEA solution (10% w/v CHAPS detergent and 25% w/v *N*-methyldiethanolamine in water) for 1 week, then further delipidated with dichloromethane/methanol as in iDISCO, then treated with 0.5M acetic acid for 2 days, washed in water for 6 hours repeated 2 times, and then treated with guanidinium solution (PBS with 4M guanidinium chloride, 0.05 sodium acetate, 2% w/v Triton X-100) for 2 days, blocked in blocking buffer (1× PBS, 0.2% v/v Triton X-100, 10% v/v DMSO, 10% goat serum) for 1 day, and finally stained in antibody incubation buffer (1× PBS, 0.2% v/v Tween-20, 3% v/v DMSO, 3% v/v goat serum, 10 mg/L heparin sodium) for 1 week.

For quantification, PV-positive cells were identified using a Laplacian of Gaussian (LoG) filter, followed by intensity-based segmentation. These segmented masks allow the quantification of bulk- and cut-staining channel intensities, in addition to the distance transformation intensity, performed in MATLAB R2023a (MathWorks, US). For an ideal deep immunostaining, the bulk-immunostaining signals should be independent of the bulk-staining penetration distances computed with distance transform of the segmented tissue boundaries, and perfectly correlate with that of cut-immunostaining. This is often not the case, as “rimming” of bulk-staining signals inevitably occurs as a “shell” around the tissue due to more easily accessible antigens on the bulk-staining tissue surface (**Fig. S1B**). The rimming effect can be quantified by fitting a single-term exponential decay curve

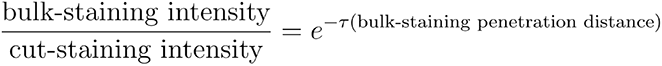

and evaluating the decay constant, tau (τ), across penetration depths, with τ → 0^+^ as we approach the ideal case.

#### Screening chemicals for INSIHGT

We first pre-screened the weakly coordinating anions by immunostaining for parvalbumin in 1mm^3^ of mouse cortex tissue cubes in the presence of 0.1M weakly coordinating superchaotropes, after 1 day of incubation at room temperature the staining solution was aspirated and 0.1M corresponding cyclodextrin was added and incubated overnight. The tissue was then washed in PBSN for 15 minutes 2 times and cleared with the benzyl alcohol/benzyl benzoate (BABB) method, and imaged. This procedure eliminated [B_12_Br_12_]^2-^, [B_12_I_12_]^2-^, and [PW_12_O_40_]^3-^ (as cesium or sodium salts) as they do not give the correct immunostaining pattern or lead to tissue destruction. We tested [Fe(C_5_H_5_)_2_]^+^ (as the hexafluorophosphate salt) for the sake of completion as a low-charge large-sized cation.

To benchmark the ability in achieving deep and homogeneous immunostaining, the above benchmarking procedure was used. Mouse hemibrains were fixed, washed and stained with 1ug rabbit anti-parvalbumin antibody with 1μg AlexaFluor 647-labeled donkey anti-rabbit secondary antibody Fab fragments in 0.1M of the weakly coordinating anions. The staining proceeded for 1 day after which the solution was replaced with 0.1M corresponding cyclodextrin (or its derivatives) and incubated overnight. The hemibrains were then washed in PBSN for 1 hour 2 times, cut in the middle coronally and re-stained for parvalbumin using AlexaFluor 488-labeled secondary Fab fragments. The tissue was then washed, cleared with the BABB method, and imaged on the cut face using a confocal microscope.

#### INSIHGT

A detailed step-by-step protocol used in this study has been given below. As a general overview, tissues were typically fixed using formalin or 4% paraformaldehyde, thoroughly washed in PBSN, and pre-incubated overnight at 37°C in INSIHGT buffer A. The tissues were then stained with a solution containing the desired antibodies, Fab fragments, lectins, and SBEβCD-complexed nucleic acid probes in INSIHGT buffer A, ensuring a final [B_12_H_12_]^2-^ concentration of 0.25 M. Staining duration varied from 6 hours to 10 days based on tissue size, antigen, and required homogeneity (please see the calculation of time *t* in the step-by-step protocol). Post-staining, the solution was aspirated and replaced with INSIHGT buffer B (0.25M 2-hydroxypropyl-γ-cyclodextrin in PBS) without prior washing, followed by a minimum 6-hour incubation with adequate shaking of the viscous buffer. After sufficient PBSN washing, tissues were ready for imaging or clearing. Over incubation for any steps up to 60 days was tolerable. After imaging, the antibodies can be eluted with 0.1M sodium sulphite in INSIHGT buffer A at 37°C overnight.

#### Screening antibodies compatible with INSIHGT

To test antibodies in a high-throughput manner, we compiled a list of antibodies, reviewed their tissue expression and staining patterns in the literature, then obtained the respective tissues known to have positive staining. These tissue blocks or entire organs were then washed, dehydrated, delipidated, rehydrated, washed, and infiltrated with INSIHGT solution A as described in the INSIHGT protocol. These INSIHGT-infiltrated tissues were then cut into ∼1mm^3^ tissue cubes and placed in a 96-well plate as indicated in the list, with each well containing 70μl of 1x INSIHGT solution A. About 0.5μg of the primary antibody to be tested was then added and 0.5μg of the corresponding AlexaFluor 647 or AlexaFluor 594-conjugated secondary antibody Fab fragment. The AlexaFluor 647 and 594 fluorophores were chosen for to minimize interference from any tissue autofluorescence on the result interpretation. For a total volume and antibodies added two each well, an equal volume of 2x INSIHGT solution A was then added to ensure the final concentration of 1x INSIHGT solution A. The plate was then sealed and the staining was allowed to proceed in the dark overnight at room temperature. The tissues were then washed in INSIHGT solution B for 2 hours, PBSN for 1 hour for two times, and then dehydrated with through 15 minutes-incubation of 50% methanol, 100% methanol and 100% methanol. The tissues were then cleared in BABB for 15 minutes and proceeded to imaging. The total fixed tissue-to-image time for the antibody compatibility test is < 36 hours.

#### Comparison between 2D histological staining of post-INSIHGT and control tissues

Mouse and human samples were pre-processed as described above. Tissues were divided into the post-INSIGHT treated group which underwent the INSIHGT protocol with 3 days of INSIHGT A incubation without the application of antibodies and 6 hours of INSIHGT B incubation, plus BABB clearing, and the control group which was immersed in PBSN for an equivalent period of time. Both groups were immersed in 70% ethanol, preceded by the immersion in 100% ethanol for the post-INSIHGT group (which were in BABB), and in 50% ethanol for the control group (which were in PBSN). Tissues were then immersed in 100% ethanol, xylene, and paraffin as in the standard paraffin embedding process. The embedded tissues were cut into 5 μm (human) or 10 μm (mouse) sections followed 2D histological staining with special stains. Following standard protocols, H&E staining was performed on human brain and kidney, PAS staining was performed on human kidney, Alcian blue staining was performed on mouse colon, and Masson trichrome staining was performed on mouse kidney samples.

#### Microscopy

Microscopic imaging was performed with either a Leica SP8 confocal microscope, a custom-built MesoSPIM v5.1 ^43^, or a two-photon tomography system as previously described ^9^.

#### RNA and DNA quality checking

Control and INSIHGT-treated samples following the 1mm^3^ treatment timeline were re-embedded in paraffin wax, and sent for nucleic acid integrity analysis services provided by the BGI Hongkong Tech Solution NGS Lab. RNA integrity number analysis was performed using the Qubit Fluorometer. Whole genome DNA quality analysis was performed using the Agilent 2100 Bioanalyzer system.

### Quantification and data analysis

#### Image processing

No penetration-related attenuation intensity adjustments were performed for all displayed images except for the 3D renderings (but not 2D cross-sectional views) in **Fig. 3** and **Movie S1** to provide the best visualization of internal signal. For samples imaged with two-photon tomography, we noticed a thin rim attributed to the heat produced during the gelatin embedding process (which we verified by repeating the staining and confirming its absence with light sheet microscopy). We employed an intensity transformation mask based on the exponent of the distance from the whole organ mask surface. Image segmentation was performed with Cellpose 2.0 ^26^ for cells implemented in MATLAB R2023b or Python, or with simple intensity thresholding. Affine and non-linear image registration was performed in MATLAB R2023a or manually in Adobe After Effects 2020 using the mesh warp effect and time remapping for *z*-plane adjustment. Image stitching was performed either with ImageJ BigStitcher plugin ^44^ or assisted manually with Adobe After Effects 2020 followed by tile allocation using custom written scripts in MATLAB R2023a.

3D image visualization and Movie rendering were performed with Bitplane Imaris 9.1, which were done as raw data with brightness and contrast adjustments, except for the whole mouse brain imaged with two-photon tomography. To remove their slicing artifacts, we resliced the volume into x-z slices, performed a z-direction Gaussian blur, followed by a 2D Fourier transform and filtered out non-central frequency peaks before inverting the transform. Finally, a Richardson-Lucy deconvolution was performed with a point-spread function elongated in the x-z direction, and resliced back into x-y slices.

#### Segmentation and analysis of podocyte-to-PEC microfilaments in mouse kidneys

Podocyte-to-PEC microfilaments of 14 mouse kidneys were manually traced via the SNT plugin in ImageJ ^45^. Path properties of the tracings were then exported for further analysis using custom codes in MATLAB R2023a. Distance transforms were performed under manually curated glomerulus and bowman space masks, such that each voxel value corresponds to the distance between that voxel and the nearest nonzero voxel of the Bowman space mask. Path displacement *d_fil_* was computed via Pythagoras theorem using the start and end coordinates of the filament . Minimal distance *d_min_* is defined as the voxel value difference between the start and end coordinates. Path length *d_path_* is directly measured via SNT. Tortuosity is defined as *d_path_* / *d_fil_*, skewness is defined as *d_fil_* / *d_min_*, and the angle of take-off is defined as the angle between the unit gradient vector of the distance transform and the unit path displacement vector. The geodesic distance *d_A_(p, q)* between voxels p, q ∈ A is defined as the minimal of length L of path(s) *P* = (*p*_1_, *p*_2_, …, *p_l_*) connecting *q, p* , where A is the set of all voxels constituting the surface of the glomerular mask ^46^:

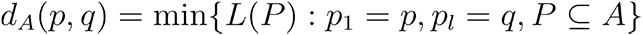

Correlation statistics were then performed via GraphPad Prism version 8 for Windows, GraphPad Software, Boston, Massachusetts USA, www.graphpad.com. Tracing and statistical analysis for the human cerebellar neurofilament inclusions were performed analogously.

#### Spatial orientation and fractional anisotropy visualization of human cerebellum neural and glial filaments

To visualize cerebellar neural and glial fibers in their preferred orientations, we performed structure tensor analysis with orientation-based color-coding in 3D.

In detail, let *G* : ℝ^3^ × ℝ_+_ → ℝ be a 3D Gaussian kernel with standard deviation σ :

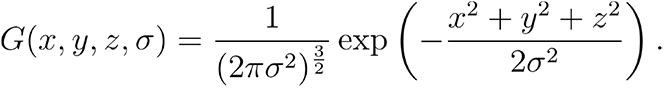

Define a 3D image as a function *I* : ℝ^3^ → ℝ which outputs the spatial voxel values. The gradient ▽*I* : ℝ^3^ → ℝ^3^ of *I* at each voxel is obtained by convolving *I* with the spatial derivatives of *G* :

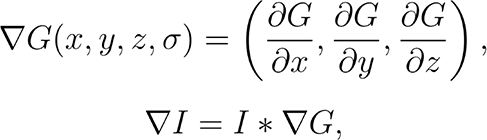

where * denotes the convolution operation.

Compute the structure tensor *T* : ℝ^3^ → ℝ^3×3^ as the outer product of ▽*I* with itself:

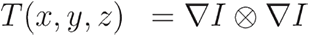

T is then smoothed over a neighbourhood *N* via convolution with *G* to give *T̄*:

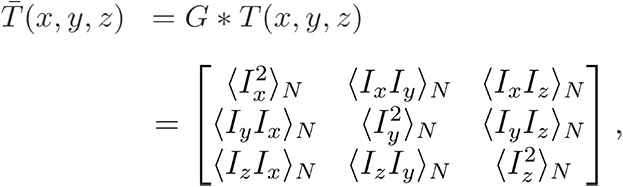

where 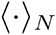 represents the Gaussian-weighted smoothing over *N* ^47,48^.

Eigendecomposition of *T̄* is then performed to define the shape (eigenvalues, λ) and the orientation (eigenvectors, *v_e_*) of the diffusion ellipsoid. The fractional anisotropy (*FA*) is then computed from λ:

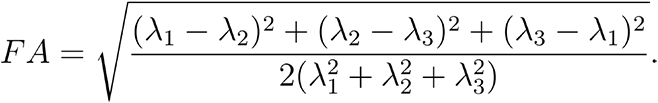

where FA ranges from 0 (complete isotropic diffusion) to 1 (complete anisotropic diffusion) ^49^.

The tertiary (least) eigenvalue-associated eigenvectors were then extracted for the 3-dimensional image volume, with the 4th dimension encoding the corresponding vector basis magnitudes. To visualize the orientation of fibers in the context of the image, the eigenvectors were intensity-modulated with both the fractional anisotropy and the original image voxel values, and represented as a 3D RGB stack for visualization in Imaris.

### Multi-round multiplexed 3D image processing and analysis

As the images were acquired across multiple rounds on a confocal microscope, we encountered the issues of misalignment and z-step glitching due to piezoelectric motor errors. Hence, the tiles of images can neither be directly stitched nor registered across multiple rounds. A custom MATLAB code was written to manually remove all the z-step glitching, followed by matching the z-steps across multiple rounds aiding by using the time-remapping function in Adobe After Effects, with linear interpolation for the transformed z-substacks. The resulting glitch-removed, z-matched tiles were then rigid registered using the image registration application in MATLAB, followed by non-rigid registration for local matching. Finally the registrated tiles were stitched for downstream processing.

Before segmentation, all non-vessel channels underwent background subtraction. They were then summed to capture the full morphology of stained cells, followed by segmentation using Cellpose 2.0 ^26^. Vessels were segmented based on their staining intensity, and a distance transform was used to obtain the distance from vessels for all voxels. The cell masks subsequently facilitated the acquisition of the statistics for all stained channels.

UMAP was performed in MATLAB R2023a using the UMAP 4.4 ^44,50^ package in a nested manner, incorporating the means and standard deviations of all immunostaining intensities, as well as the distance to the nearest blood vessel. An initial UMAP was applied to each image stack tile, followed by DBSCAN clustering to eliminate the largest cluster based on cell count. The remaining cells were subjected to a second UMAP, where another round of DBSCAN clustering yielded the final cell clusters for analysis.

Violin plots for each clustered cell type’s distance from neuropeptide Y-positive fibers were obtained by creating a distance transformation field from the segmented fibers. Segmented cell masks were used to compute the mean intensity value of the distance transformation field. The pairwise distances of the clustered cell types were obtained for the 30 nearest neighbors, followed by calculating the mean and SD for the coefficient of variation. The gramm package in MATLAB R2023a was used for plotting some of the graphs ^51^

### INSIHGT Protocol

This protocol describes the INSIHGT process for staining fixed or permeabilized tissues

#### Materials

|  |  |
| --- | --- |
| PBSN (1× PBS with 0.02% w/v $\text{NaN}_3$) | Sodium dodecahydrododecaborate ($\text{Na}_2\text{B}_{12}\text{H}_{12}$) |
| Methanol | 2-hydroxypropyl-γ-cyclodextrin (2HPγCD) |
| Dichloromethane | Nucleic acid probe (optional) |
| Sodium sulphite ($\text{Na}_2\text{SO}_3$, optional) | Sulfobutylether-β-cyclodextrin (SBEβCD, optional) |

#### Buffers/Solutions

INSIHGT buffer A (2×): 0.5M Na_2_B_12_H_12_ in PBSN INSIHGT buffer B (1×): 0.25M 2HPγCD in PBSN INSIHGT buffer C (1×): 0.25M SBEβCD in PBSN

#### Procedure overview

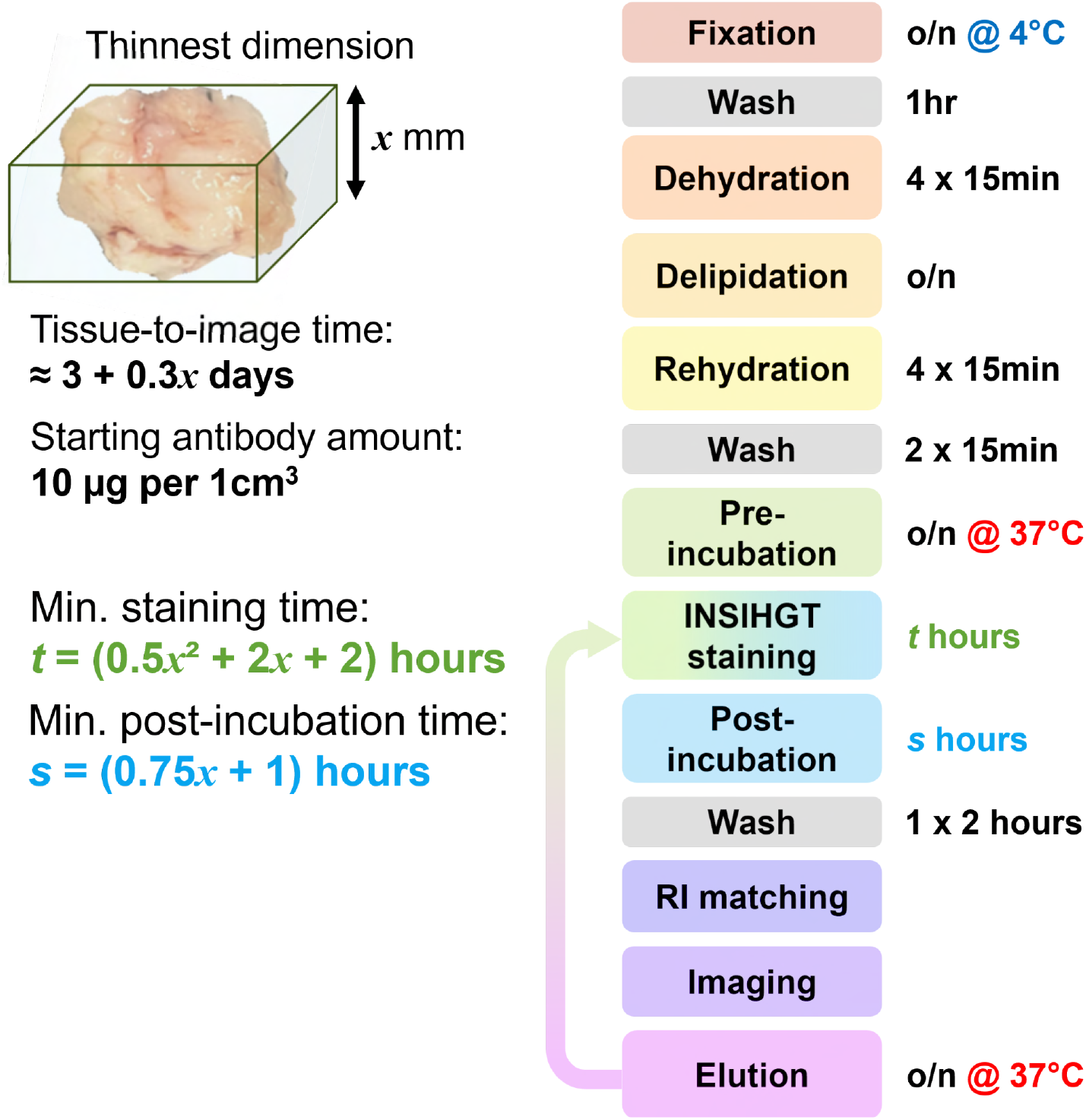

#### Procedure

The procedure below is illustrated for a 1 cm³ tissue sample but can be scaled as needed.

1. **Fixation:** Fix tissue overnight at 4°C with formaldehyde, glutaraldehyde or SHIELD-fixation. Wash in 5 ml PBSN for 1 hour at room temperature (r.t.) with shaking. For glutaraldehyde or SHIELD-fixation, quenching should be done appropriately.
2. **Dehydrate** by shaking incubation in 5 ml 50% v/v methanol/water, then 5 ml 100% methanol repeated 3 times, 15 min with shaking at r.t. for each step.
3. **Delipidate** by shaking incubation in 5 ml 2:1 v/v mixture of dichloromethane/methanol overnight at r.t. Use a glass or polypropylene containers (e.g. microcentrifuge tubes), in a well-ventilated hood with adequate shaking.
4. **Rehydration:** Repeat the dehydration steps in reverse order, then 2 × 15 min washes in PBSN.
5. **Pre-Incubation:** Incubate in 2 ml 1× INSIHGT buffer A at 37°C overnight with shaking.
6. **Prepare nucleic acid probe (optional):** Prepare 200 nmol of the probe by mixing it with INSIHGT buffer C in a 1:1 v/v ratio.
7. **INSIHGT Staining:**

a. Place the tissue in a container that fits its size well.
b. Add enough 2× INSIHGT buffer A to around half-cover the tissue.
c. Add the desired amount of primary antibodies (start with 10μg each).
d. Add the prepared nucleic acid probe in INSIHGT buffer C (if desired).
e. Add corresponding fluorophore-labeled secondary antibody Fab fragments or nanobodies (in the same μg amount as the primary antibody used), or lectins if desired.
f. Add PBSN to make the final concentration of INSIHGT buffer A to 1x. The volume required will be equal to the volume added in step 6a - (total volume added in step 6c, 6d, and 6e).
g. Top up with 1x INSIHGT buffer A to just cover the entire tissue.
h. Incubate at r.t. for 3 days (or *t* hours, see below) with shaking. Protect from light.
8. **Post-incubation:** Incubate the stained tissue in 3 ml INSIHGT buffer B at r.t. overnight (or *s* hours, see below) with shaking. Place the tube horizontally to ensure adequate mixing.
9. **Wash**: Wash in 5 ml PBSN three times for 2 hours each at r.t., shaked horizontally.
10. **Refractive index matching** (optional): Example for BABB clearing:

- Repeat the dehydration steps.
- Prepare BABB: 1:2 v/v mix of benzyl alcohol / benzyl benzoate.
- Place tissue in 5ml BABB at r.t. overnight with shaking. Refresh BABB until tissue is clear.
11. **Imaging**: Confocal, light-sheet and two-photon microscopies are all compatible.
12. **Stripping** (optional): Strip tissue in 2 ml 1× INSIHGT buffer A + 0.1 M Na_2_SO_3_ at 37°C overnight with shaking. Wash twice with 2 ml 1× INSIHGT buffer at 37°C for 2 hours each.
13. **Repeat staining** (optional): Proceed to step 6 for the next round of staining.

#### Notes

- All incubation periods can be extended up to one month without harm.
- Tissues delipidated with methods other than CH_2_Cl_2_/Methanol are acceptable, except if the buffer contains Triton X-100 or triethylamine. Tween-20 is compatible.
- Test pretreatment buffer compatibility with INSIHGT. Mi× 100μl of the buffer with 2× INSIHGT buffer A in a 1:1 ratio. If precipitation occurs, test each buffer component individually and exclude any that cause precipitation from your modified buffer.
- Avoid contact between 100% dichloromethane and plastics (except with polypropylene or PTFE tips or Eppendorf tubes). It dissolves plastics and makes the solution cloudy.
- The minimum staining time (*t* hours) in step 8 depends on the thinnest dimension (*x*, in mm) of the tissue. Calculate *t* as 0.5*x*² + 2*x* + 2.
- The minimum post-incubation time (*s* hours) in step 9 is *s* = 0.75*x* + 1.
- If staining is uniform, staining time can be reduced. Conversely, if penetration is inhomogeneous, extend the staining time by 1.5×. There is no need to adjust post-staining incubation and washing.
- Increasing antibody concentration in the staining solution by 1.5× can also enhance penetration homogeneity. Conversely, if staining is uniform and signal-to-noise is satisfactory, antibody concentration can be reduced.
- If using fluorophore-labeled whole IgG secondary antibodies or streptavidins, perform steps 6–8 with primary antibodies only, then repeat for secondary antibodies starting from step 4. There is no need to post-fix primary antibodies.
- To visualize endogenous fluorescence, staining them with anti-fluroescent protein antibodies is the best solution. Alternatively, one can attempt ethyl cinnamate (ECi) ^52,53^ with triethanolamine replacing triethylamine as pH adjusting agent, or dibenzyl ether clearing after tetrahydrofuran (THF) dehydration ^54^. SHIELD-based protection with aqueous-based refractive index matching can also be used after omitting Triton X-100 in all the buffers.
- After staining, the sample can be stored in PBSN at 4°C or in BABB at r.t. indefinitely.

#### Graphical summary of INSIHGT protocol

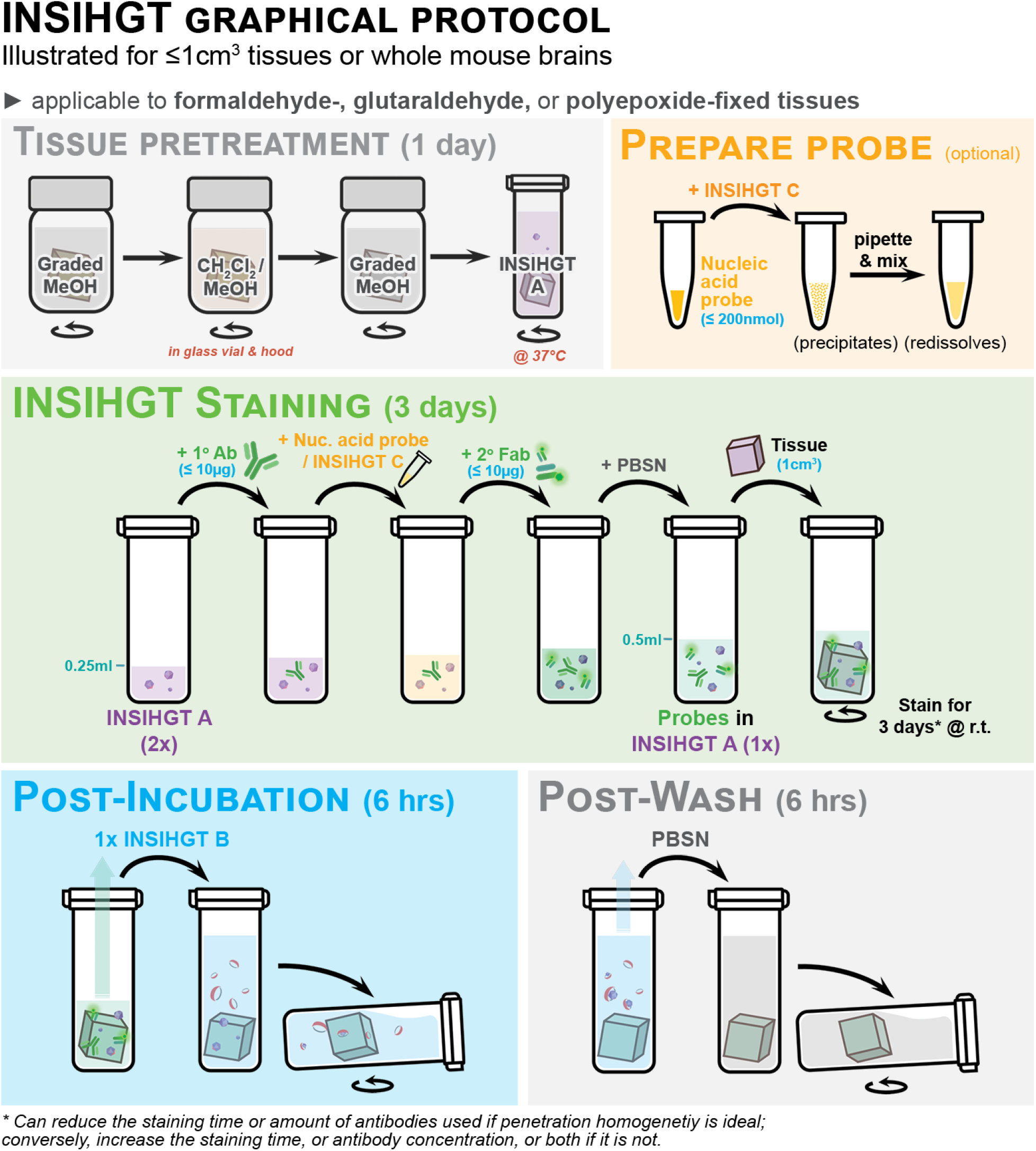

## Supplementary Text

### Weakly coordinating superchaotropes as non-denaturing protein solubilizers

For proteins, the significant role of intramolecular interactions in intermolecular interactions complicates the picture. Ideally, intermolecular interactions were selectively disturbed for solubilization, with minimal disturbance on intramolecular interactions to prevent denaturation. We reasoned that if the interactions between the native solvent (i.e., water) and proteins are promoted, the intramolecular forces (e.g., hydrophobic interactions) of proteins that evolved in an aqueous environment will help maintain their current folded state. This contrasts with agents that have significant affinities towards the proteins, which may compete against the intramolecular forces and disrupt their folding, such as with detergents ^55^ and strong hydrogen-bonding agents like urea ^56^. Hence, an ideal protein solubilizer is water soluble yet poorly coordinating with both water and proteins. The ideal weakly-coordinating superchaotropes have low charge-to-volume ratios and lack hydrophobic groups to avoid affinities towards protein hydrophobic surfaces. We thus screened the literature for counter anions used in the isolation of extremely electrophilic species for crystallography ^57^ or as conjugate bases of superacids, which includes perchlorate, perrhenate, and dodecaborates.

### Physicochemical properties of closo-dodecahydrododecaborate

The *closo*-dodecahydrododecaborate [B_12_H_12_]^2-^ was discovered in the 1950s and stands out as a remarkably stable member of boron cluster compounds and boron hydrides. It can withstand sustained dry heating, strong acids and alkalis, and diverse reactive organic or inorganic chemicals. Its chemical inertness has been attributed to the 3D aromaticity of the ion - a Hückel’s rule generalization in planar aromatic compounds. Apart from chemical inertness, [B_12_H_12_]^2-^ is also poorly coordinating, exemplified by the super-acidic character of its Brønsted acid H_2_B_12_H_12_ (p*K*_a_ = 28) ^58^ and perhaps by the stabilization of the conjugate base by delocalizing the two electrons over a large surface area. The combined weak ionic and poorly coordinating properties lead to easy solubilization of the [B_12_H_12_]^2-^ ion in diverse solvents, lending the proposal for it to be used for nuclear waste treatment by extracting charged ions into water-immiscible organic solvents. The chemical and coordinative inertness of [B_12_H_12_]^2-^ is also directly related to its biological inertness, with an oral lethal dose (LD_50_) for rats at > 7,500 mg/kg. Hence, the proposed biological applications for [B_12_H_12_]^2-^ and its ^10^B isotopic derivatives were largely related to boron neutron capture therapy (BNCT) *in vivo*. The utility of [B_12_H_12_]^2-^ *in vitro* has been scarcely explored. The available solution characteristics of [B_12_H_12_]^2-^ is summarized in **Table S4**.

### closo-Dodecahydrododecaborate hydration characteristics

**Table S4.** Basic physical chemistry characteristics of [B_12_H_12_]. ^2–59,60^

|  |  |
| --- | --- |
| Ion radius | 400 pm |
| $T\Delta S_{\text{struct,water}}$ | 14.8 kcal mol<sup>-1</sup> |
| $\Delta\text{HB}$ | -2.31 |
| $\Delta G_{\text{hyd}}$ | -140 kcal mol<sup>-1</sup> |
| $\Delta H_{\text{hyd}}$ | -145 |
| $T\Delta S_{\text{hyd}}$ | -5 kcal mol<sup>-1</sup> |
| Polarizability | 25.7 Å<sup>3</sup> (calculated) |
| Hydration $\Delta r$ | 9.6 pm (calculated) |
| Average number of H<sub>2</sub>O molecules in hydration shell | 1.8 (calculated) |
| $\Delta_{\text{hyd}}S$ | -17.9 cal K<sup>-1</sup> mol<sup>-1</sup> |
| $K_a$ with $\alpha\text{CD}$ in water | 100 L mol<sup>-1</sup> |
| $K_a$ with $\beta\text{CD}$ in water | 200 L mol<sup>-1</sup> |
| $K_a$ with $\gamma\text{CD}$ in water | 2,000 L mol<sup>-1</sup> |

### Interactions of [B_12_H_12_]^2-^ with other molecular species

The pair correlation function of [B_12_H_12_]^2-^ ion in water has been obtained by simulation. It showed concentric rings of water molecules on top of the hydrogen atoms, with an average number of water molecules forming ≈10 bridging O–H···H–B dihydrogen bonds (DHB) ^61^, with a simulated DHB length of 1.76 Å ^62^ with an estimated bond strength of 7.15 kcal mol^-1^ ^63^ (*ca.* O–H···O hydrogen bond length in water: 1.97 Å, bond strength 5.0 kcal mol^-1^). Simulated translational diffusion coefficients of the [B_12_H_12_]^2-^ ion in water is (1.8 ± 0.2) × 10^-9^ m^2^ s^-1^.

Apart from forming DHBs, [B_12_H_12_]^2-^ do have some coordinating activity as per evidence obtained in crystalline solid state ^64^ and has been shown to interact with adenine directly in aqueous solutions. ^60^

Interestingly, at concentrations up to 1.5mM, [B_12_H_12_]^2-^ does not disrupt the structure of bovine serum albumin (BSA) at all in 0.1M NaHCO_3_, does not show any significant interactions with BSA or affect the hydrodynamic radii of BSA even at near 1,000-fold molar excess. It also did not affect the thermal denaturing behavior of BSA. ^65^

## Supplementary Figures

**Supp Fig.1.**
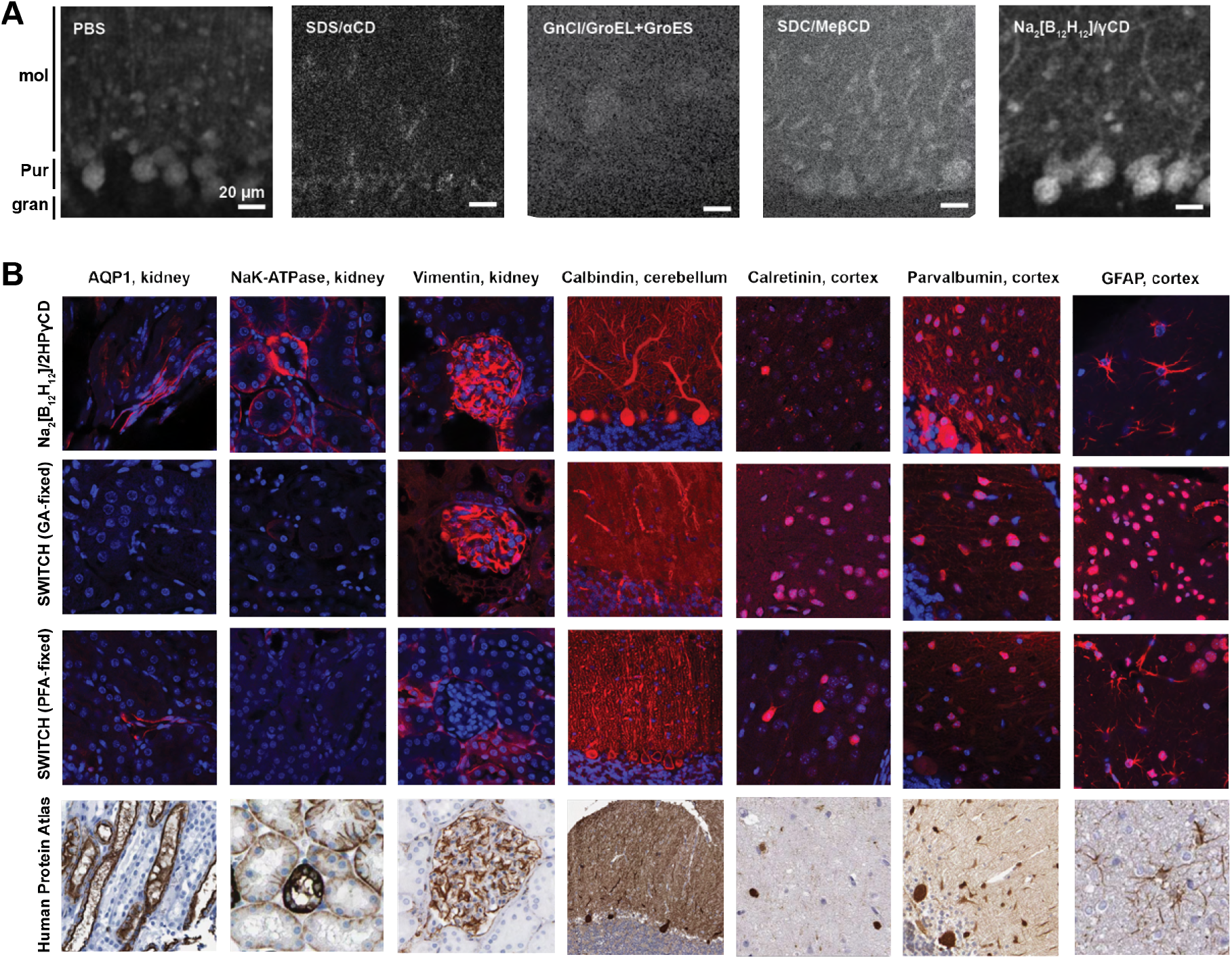
**(A)** Comparison of proposed antibody-antigen inhibition strategies and their recovery approaches. All stainings were performed in mouse cerebellum tissue for PVALB, with antibodies first co-incubated in a buffer containing an antibody-antigen interaction inhibitor followed by a measure to reinstate antibody-antigen binding. PBS: staining for PVALB done in PBS as a control. SDS/αCD: inhibition with 10mM SDS, followed by its complexation with alpha-cyclodextrin (αCD) for recovery, GnCl/GroEL+GroES: inhibition with 6M guanidinium chloride (GnCl) followed by in situ refolding with GroEL and GroES molecular chaperones, SDC/MeβCD: inhibition with sodium deoxycholate (SDC) followed by its complexation with random methylated beta-cyclodextrin (MeβCD) for recovery, Na_2_[B_12_H_12_]-γCD: inhibition with 0.125M sodium dodecahydro-closo-dodecaborate (Na_2_[B_12_H_12_]) followed by its complexation with gamma-cyclodextrin (γCD) for recovery. **(B)** Broader antibody compatibility and more specific staining was observed with the use of Na_2_[B_12_H_12_]-2HPγCD system with conventional paraformaldehyde fixation (upper row), compared with the SWITCH-labelling method reported by Murray *et al* ^14^, regardless of whether the tissue was fixed as described using glutaraldehyde (GA-fixed) or paraformaldehyde (PFA-fixed, i.e., no SWITCH-fixation employed). Lower row: Human tissue immunostaining patterns from the Human Protein Atlas for comparison.

**Supp Fig.2.**
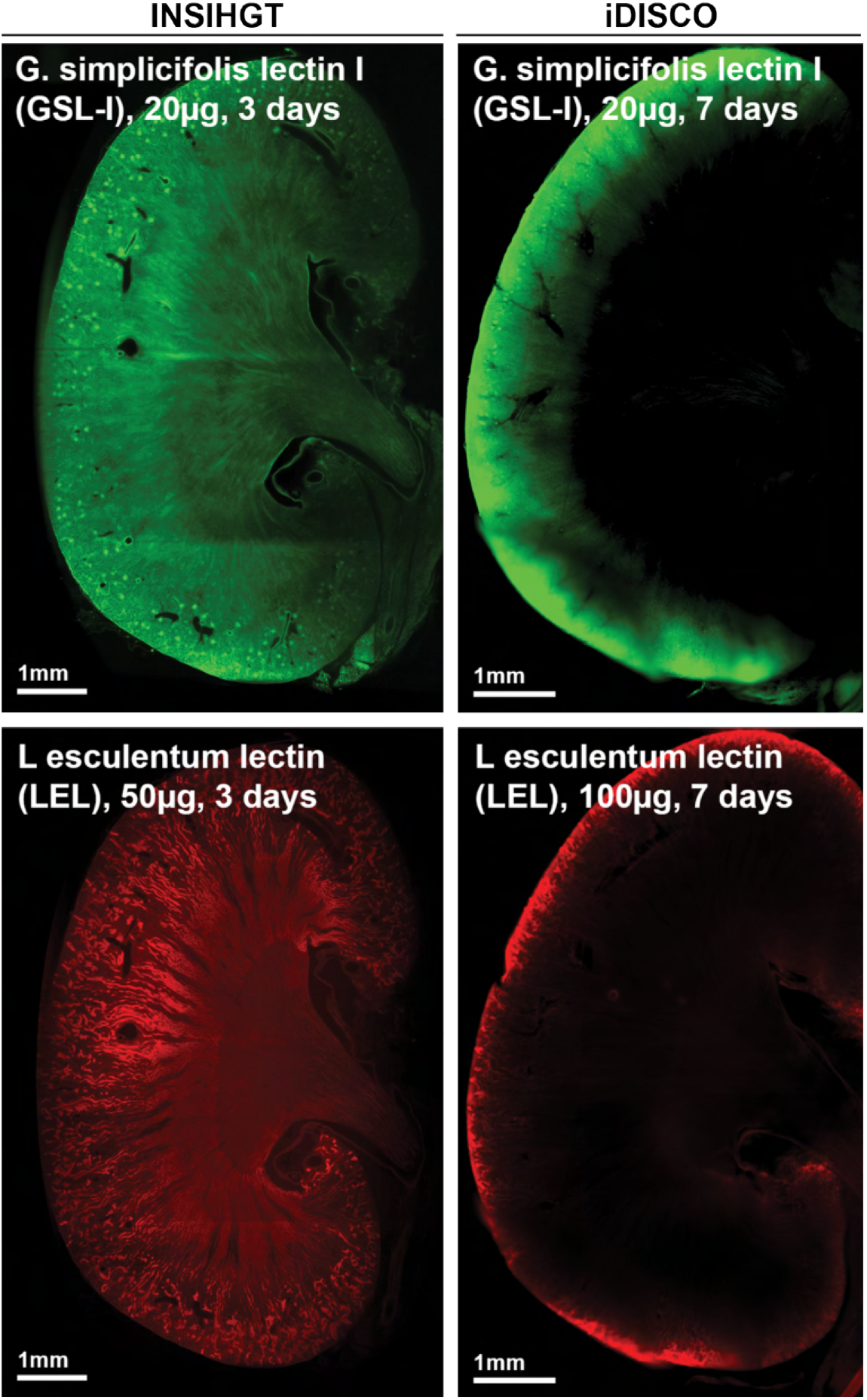
Comparison of lectin staining results between INSIHGT and iDISCO. Lectin staining often results in a significant rimming pattern where only superficial layers are stained, as demonstrated via iDISCO staining of mouse kidney with GSL-I and LEL.

**Supp Fig.3.**
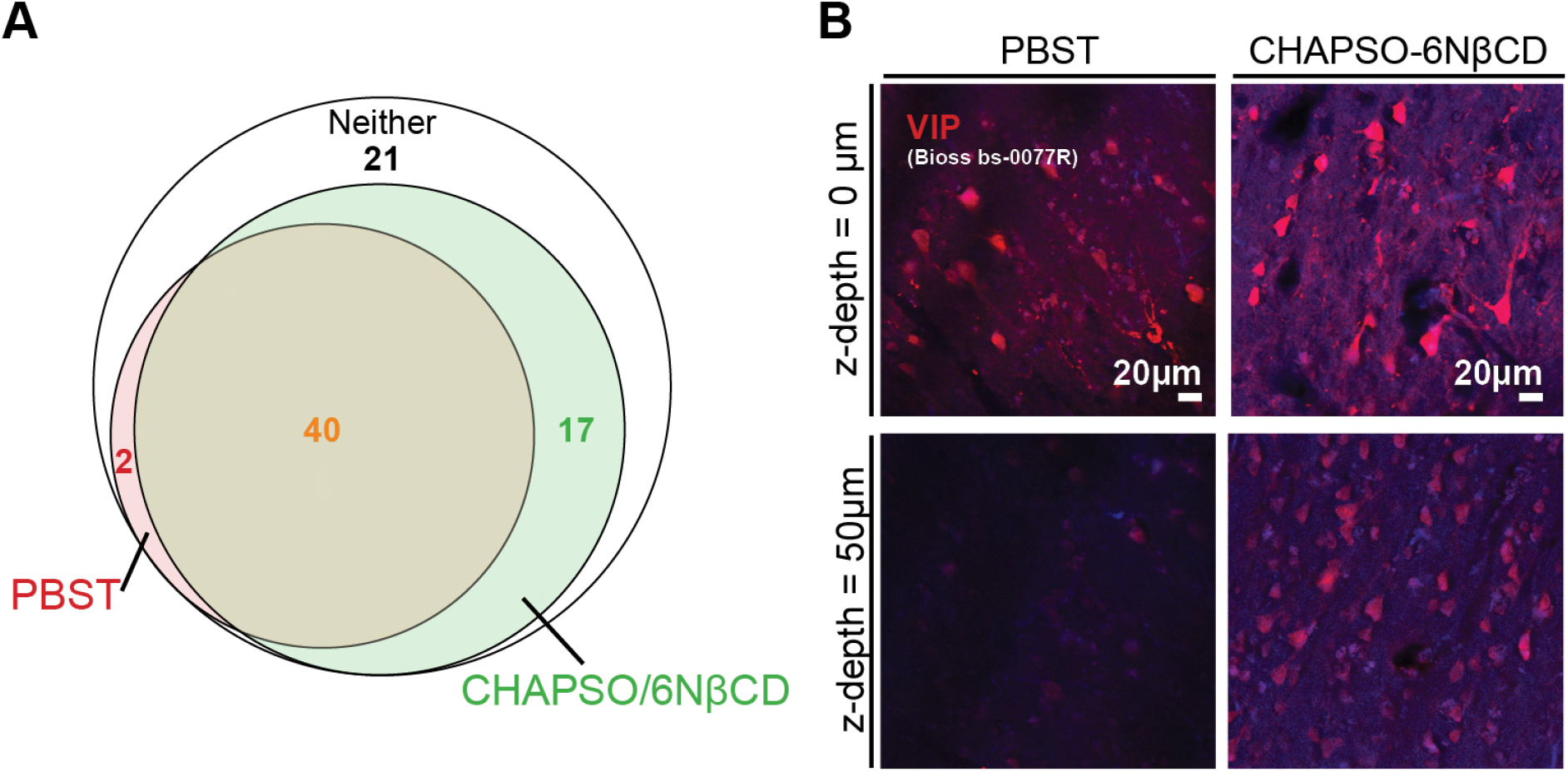
**(A)** The use of an optimized artificial chaperone host-guest system, CHAPSO/6NβCD, is compatible with 57 out of 80 validated commercial primary antibodies on human cancer tissue, human brain tissue, mouse kidney tissue and mouse brain tissue, compared to the use of PBS with 1% Triton X-100 (PBST) as the staining buffer. **(B)** Examples in the head-to-head staining penetration comparison using human brain tissue. DAPI signals were colored in blue, while antibody signals were colored in red.

**Supp Fig.4.**
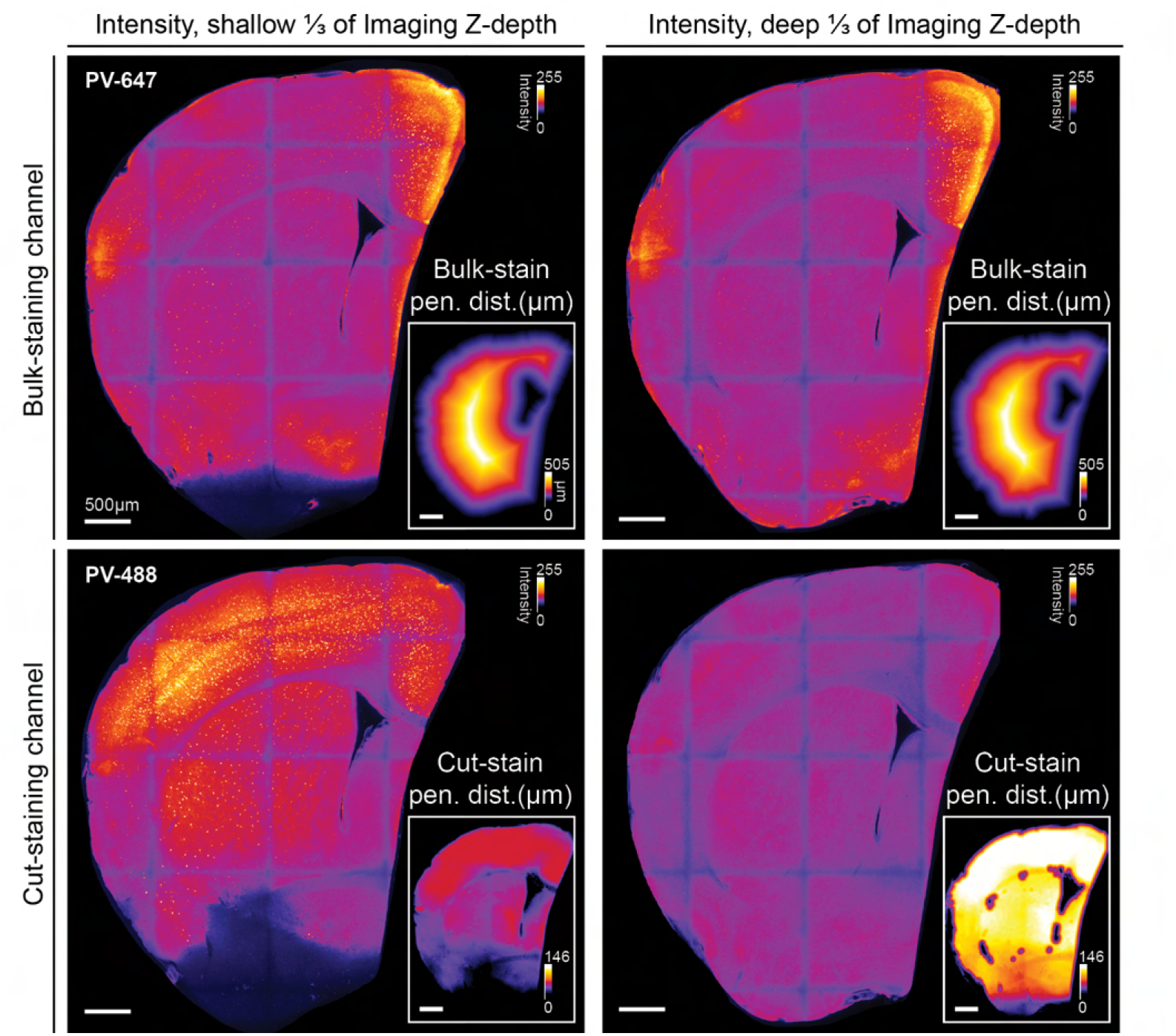
Benchmarking Experiment Design and Principle Illustration. Typical imaging results from a sample in the benchmarking experiment. The bulk-staining was conducted before tissue cutting, so the bulk-staining intensities (upper row) and their corresponding calculated penetration lengths (insets of upper row) are relatively invariant at the same *z*-depth of imaging (arranged in columns). Conversely, cut-staining was performed after tissue cutting, and antibodies could penetrate through the cut surface, which corresponds to the imaging plane, displayed in the *xy*-plane here (lower row). The penetration distance was irregular and inhomogeneous due to minor tissue curvatures and irregular tissue texture, and areas of lower penetration distances due to vasculatures. These factors were taken into account during the quantification of cut-staining penetration depth (insets of lower row).

**Supp Fig.5.**
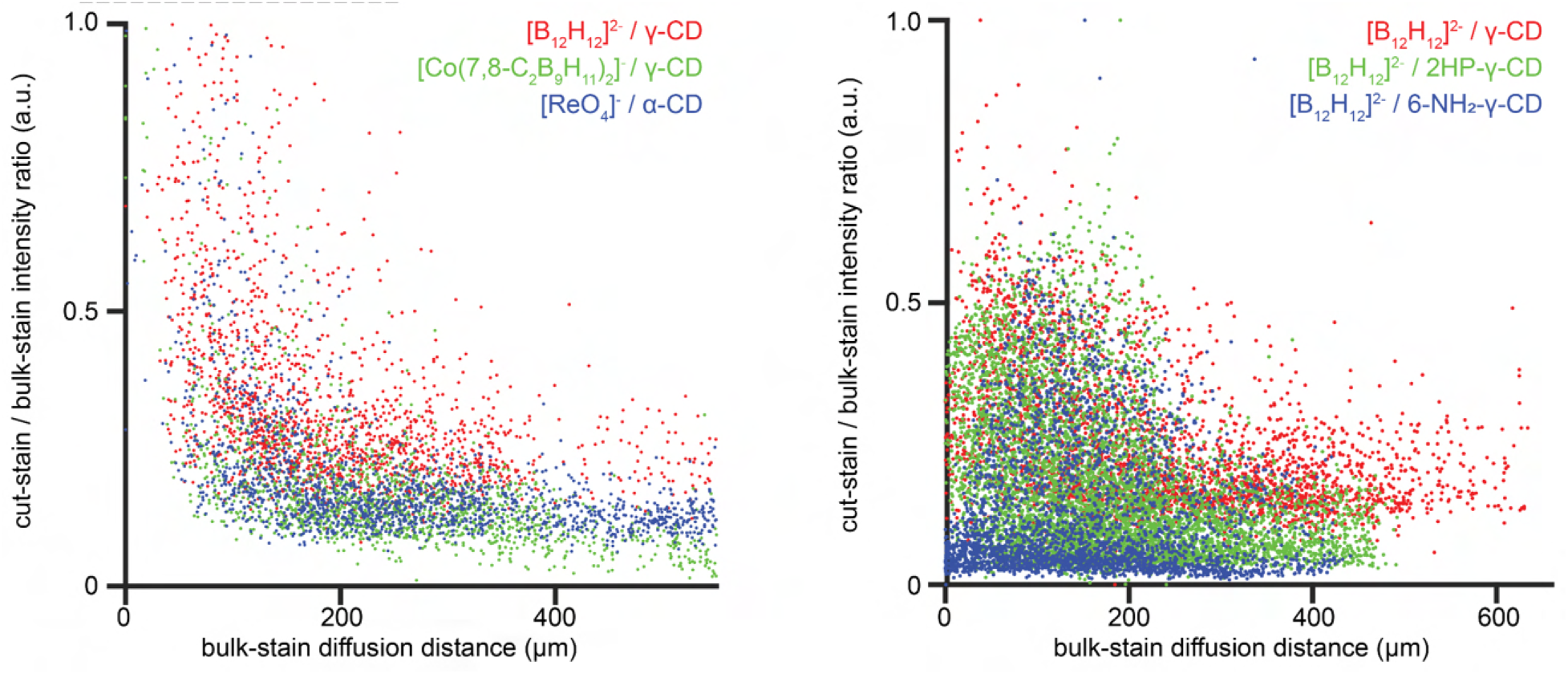
Evaluation of different weakly coordinating anions (left panel) and γ-cyclodextrin derivatives (right panel) both at 0.1M concentration using the benchmarking experimental pipeline as described previously ^6,9^.

**Supp Fig.6.**
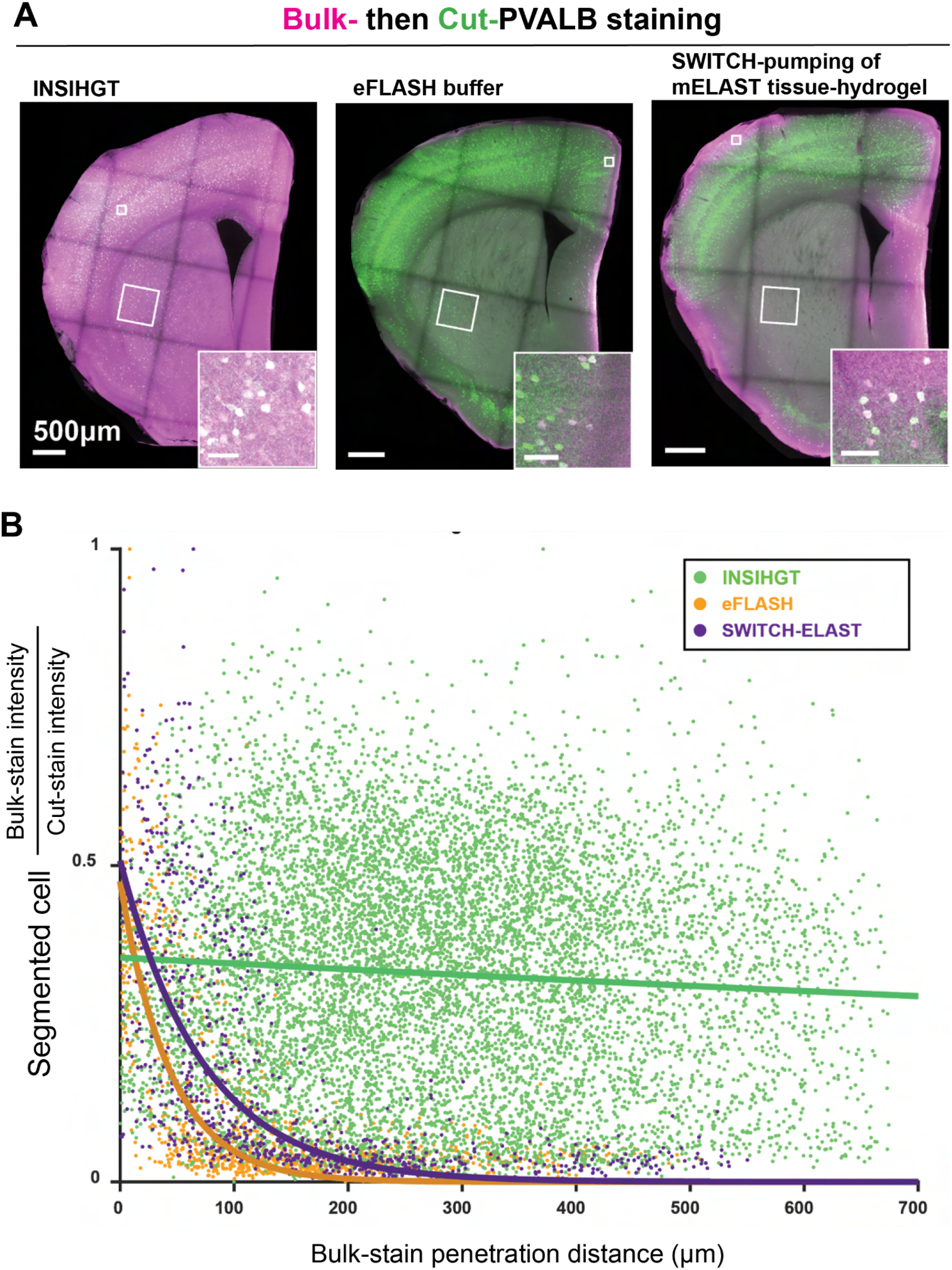
**(A)** Comparison of INSIHGT against binding kinetics modulatory buffers in eFLASH and SWITCH-pumping of mELAST tissue hydrogel buffer. Panel of INSIHGT is the same as **Fig 2B**. **(B)** Quantification of bulk:cut-staining signal ratio against penetration distance for segmented cells in the same procedures described in Fig. 2B.

**Supp Fig.7.**
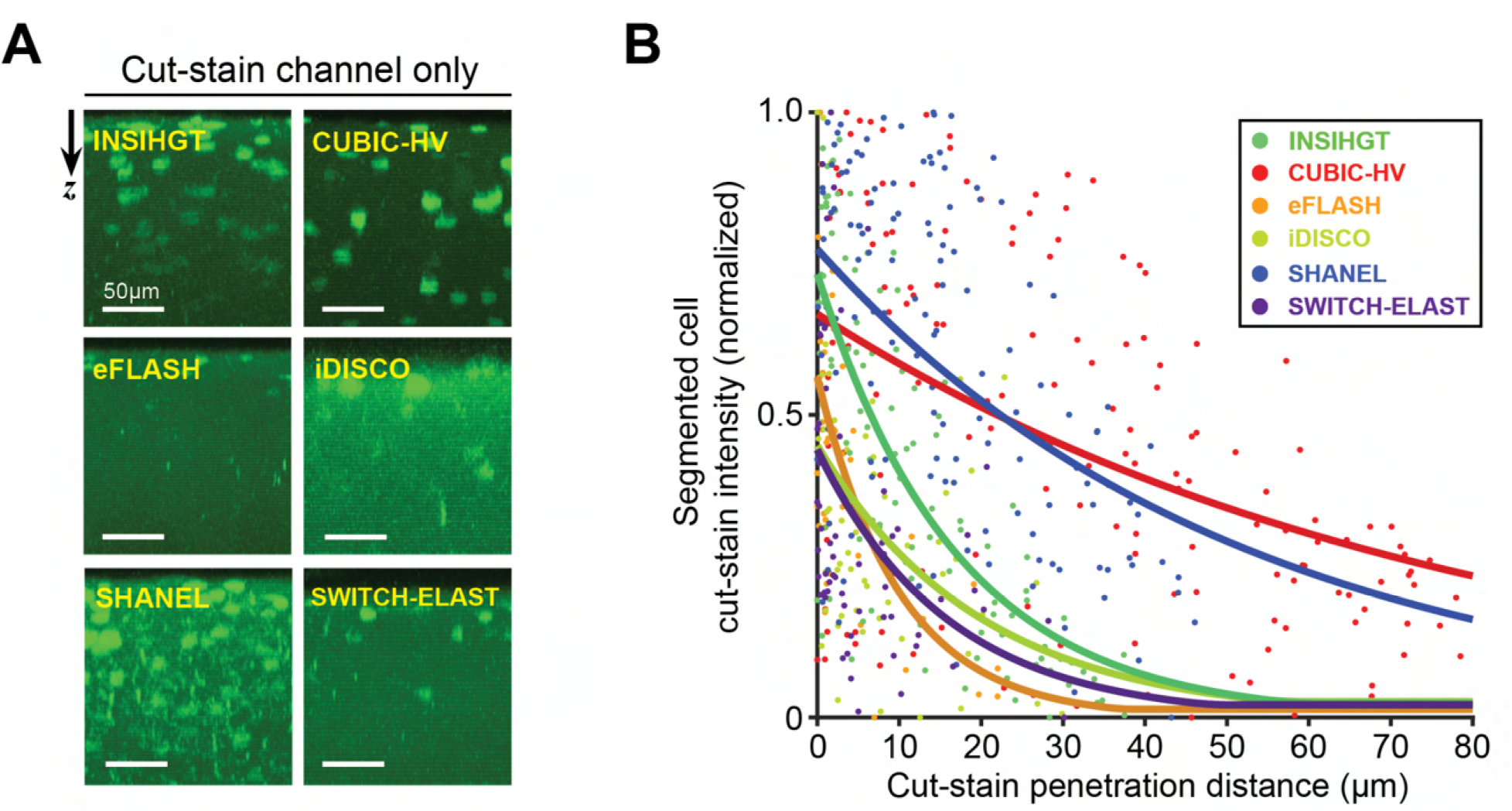
(**A**) *x-z* projection of larger white boxed areas from Figure 2B **and S6A** at deep bulk-staining penetration depths, with only the cut-staining channels (without the use of any deep immunostaining methods) shown. The deeper the penetration, the more permeabilized the tissue. (**B**) Quantitative comparison of tissue permeabilization by correlating cut-staining penetration depth and cut-staining intensity.

**Supp Fig.8.**
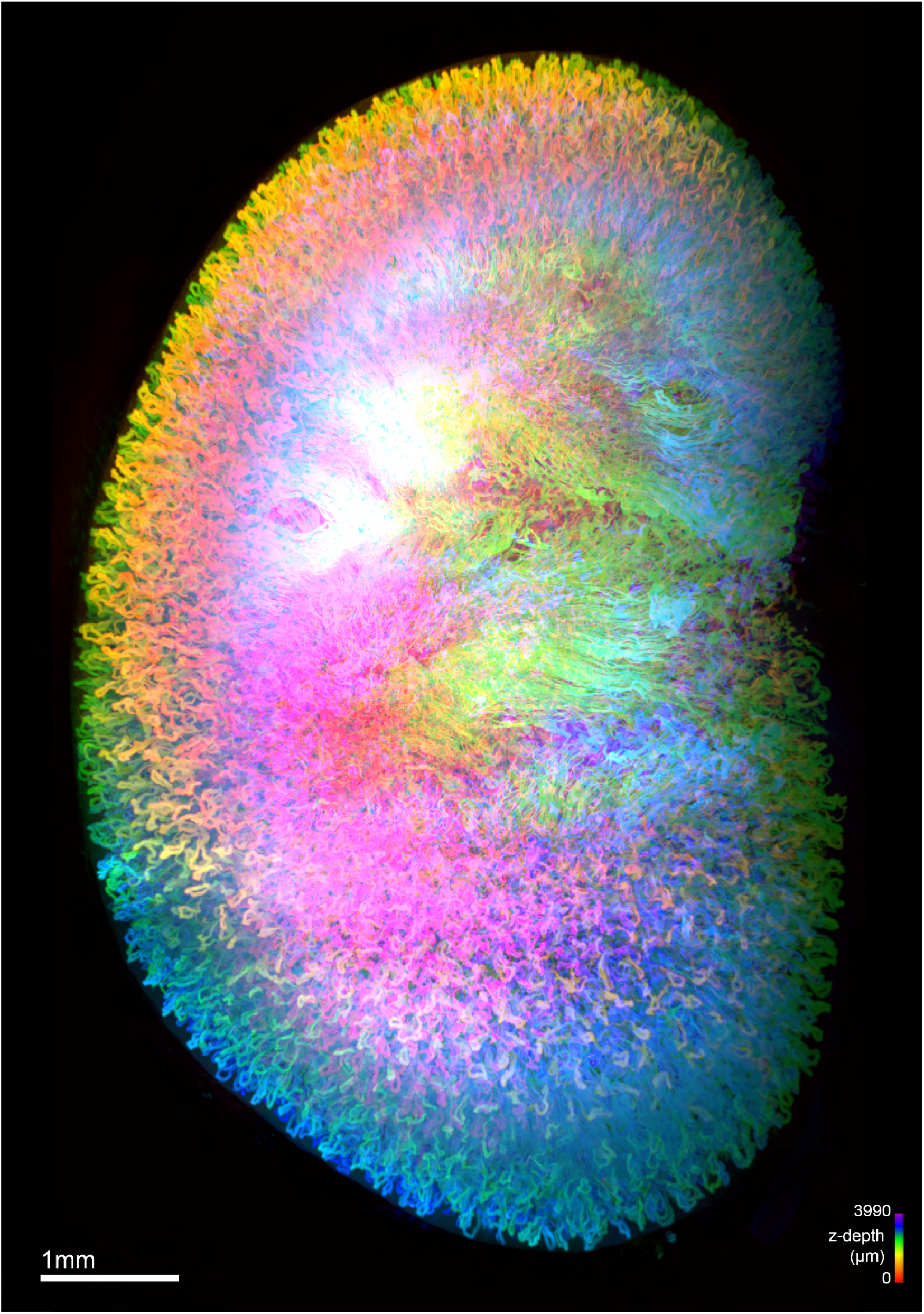
Whole mouse kidney stained with Lycopersicon esculentum (Tomato) Lectin. 3D rendering, colour-coding: z-depth.

**Supp Fig.9.**
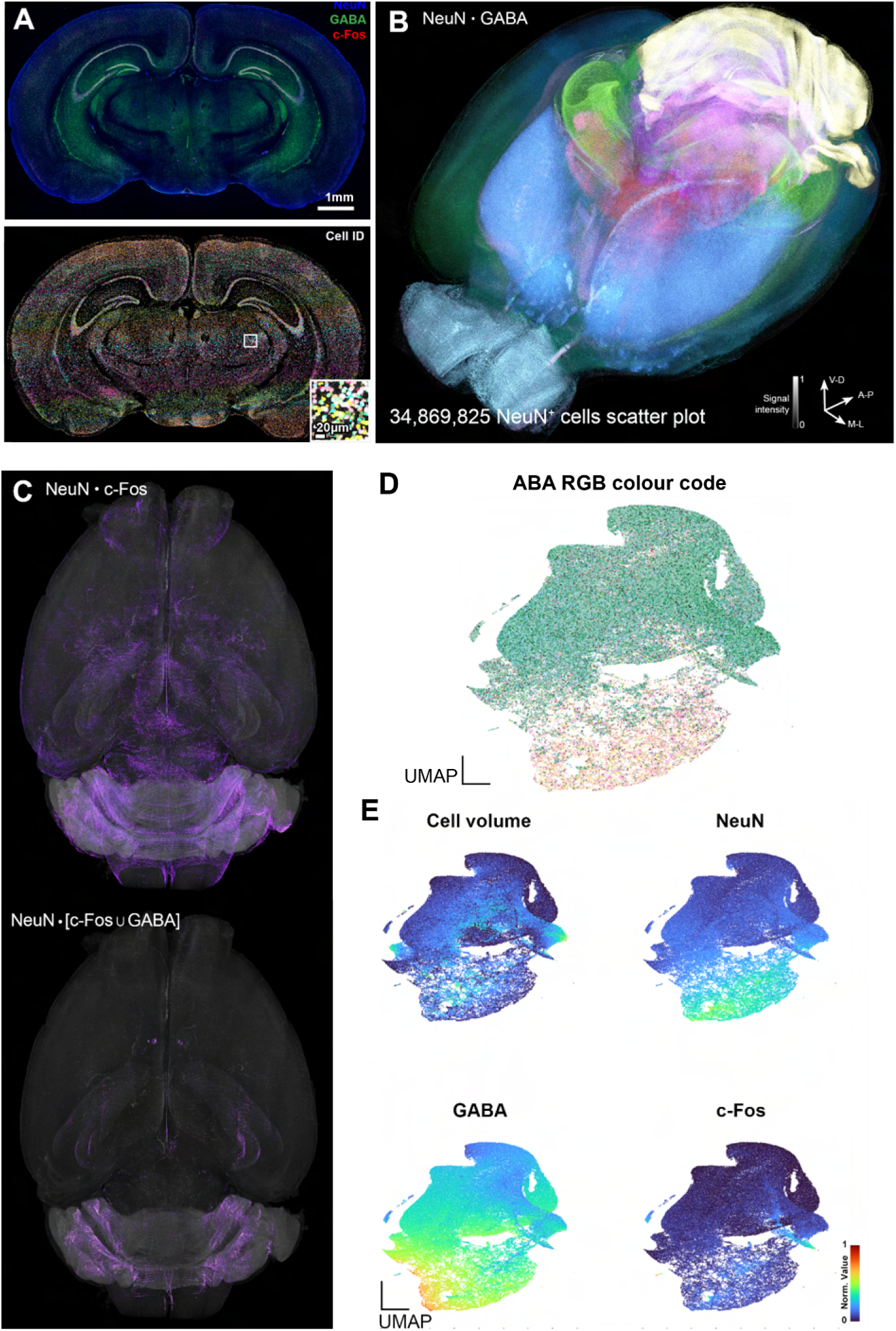
INSIHGT enables whole brain structural molecular functional quantification. **(A)** Whole mouse brain multiplexed mapping of over 34 million neurons by molecular (NeuN), neurotransmitter type (GABA), and functional (c-Fos) mapping. An example coronal slice and its corresponding 3D segmentation results are shown in the upper and lower panels, respectively. Inset in the lower panel shows the cell masks in the white boxed area. **(B)** Rendered view of all segmented cells, color-coded according to brain regions with intensity modulated by a combination of marker levels. The RGB color codes for the brain regions are based on that of the Allen Brain Atlas CCFv3. **(C)** Whole-brain active neurons and GABAergic neurons inferred by double positivity in c-Fos and NeuN (upper panel) and triple positivity in c-Fos, NeuN and GABA (lower panel). **(D)** UMAP embedding of 0.35M (1%) segmented NeuN+ cells in **(B)** with spatial regional classification. UMAP embedding was based on cell volume, NeuN and c-Fos expression levels and GABA levels detected with INSIHGT. These normalized values were color-coded in **(E)**.

**Supp Fig.10.**
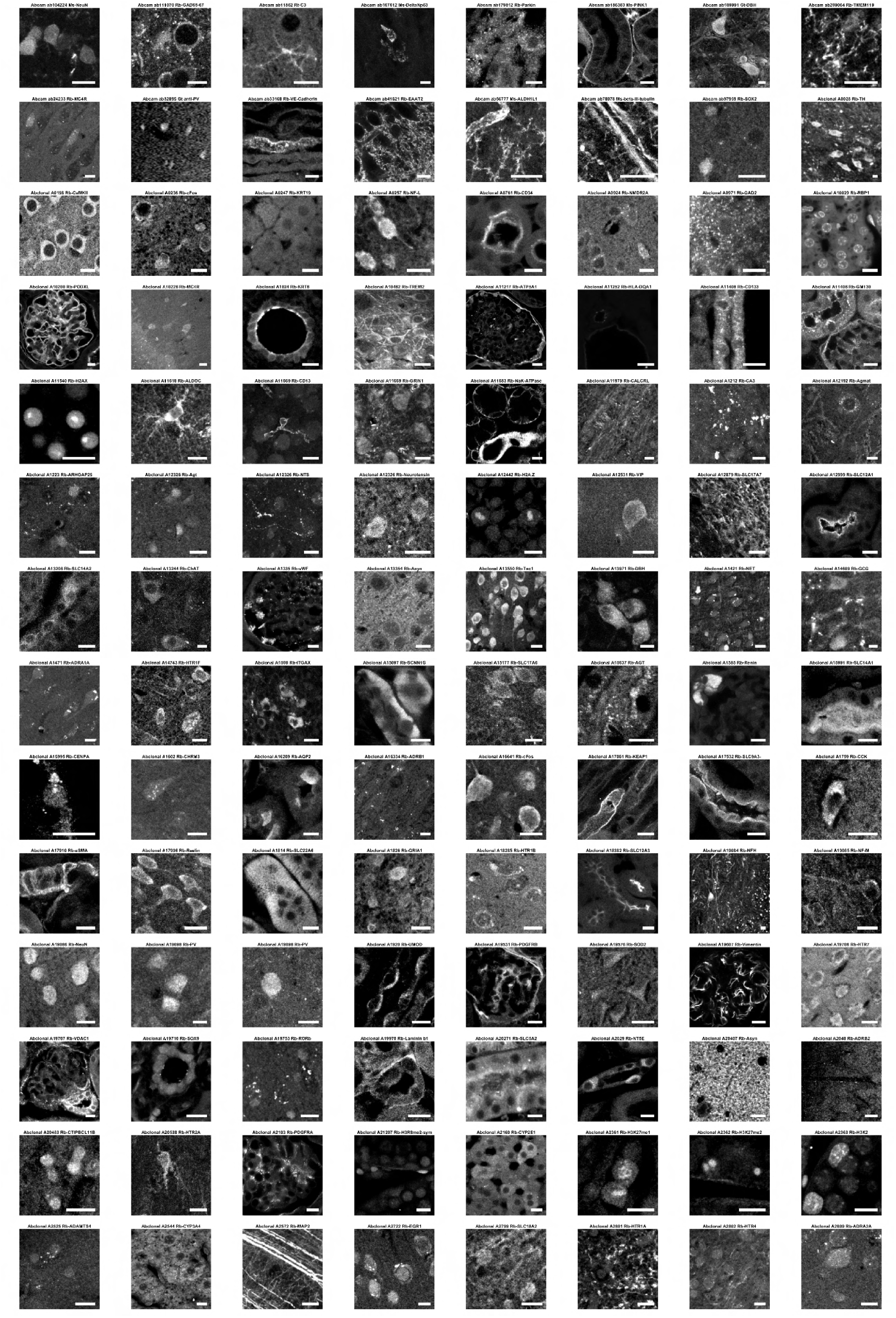
Representative images from INSIHGT-compatible antibodies, Part 1. Scale bars are set to 10 μm.

**Supp Fig.11.**
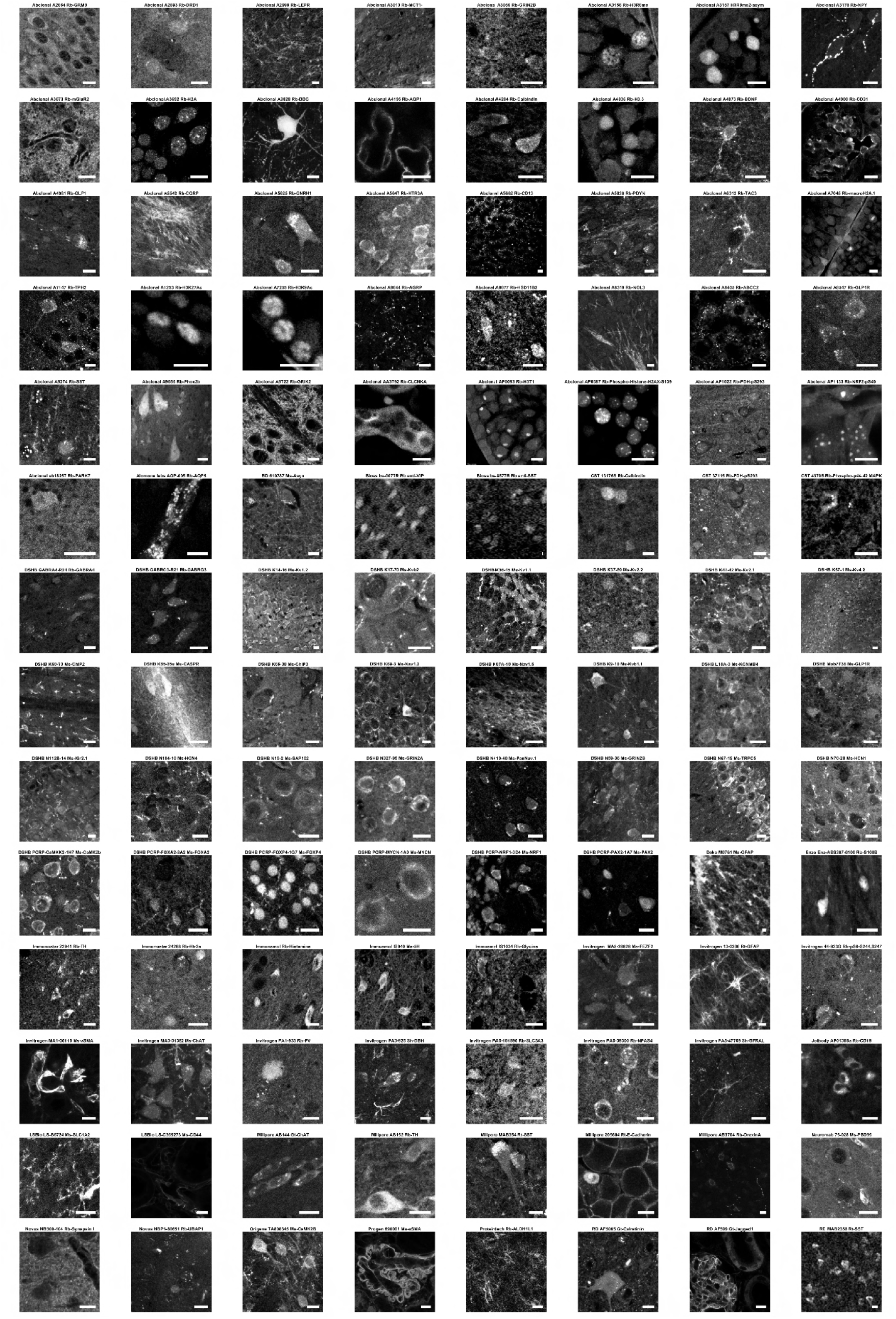
Representative images from INSIHGT-compatible antibodies, Part 2. Scale bars are set to 10 μm.

**Supp Fig.12.**
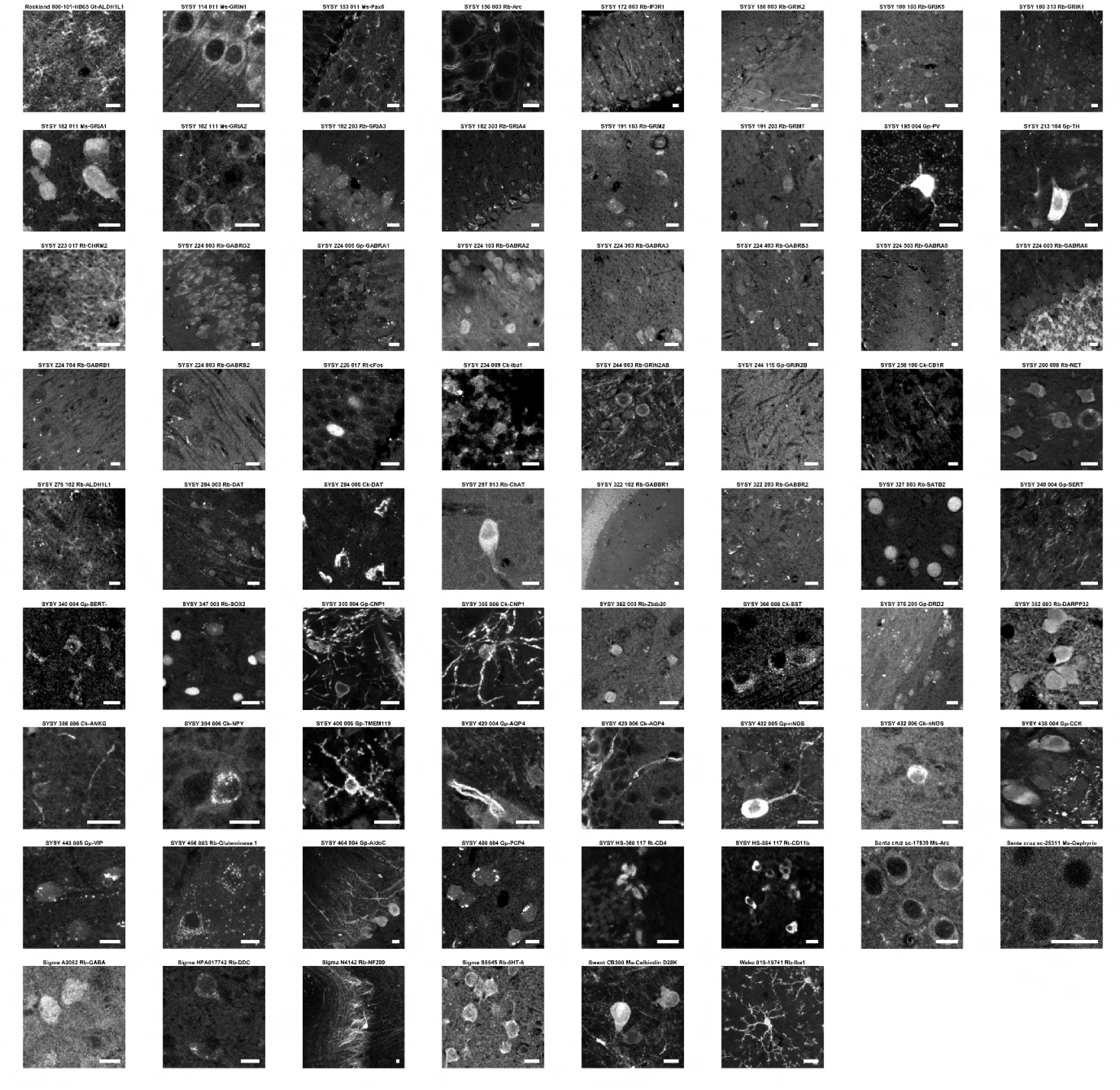
Representative images from INSIHGT-compatible antibodies, Part 3. Scale bars are set to 10 μm.

**Supp Fig.13.**
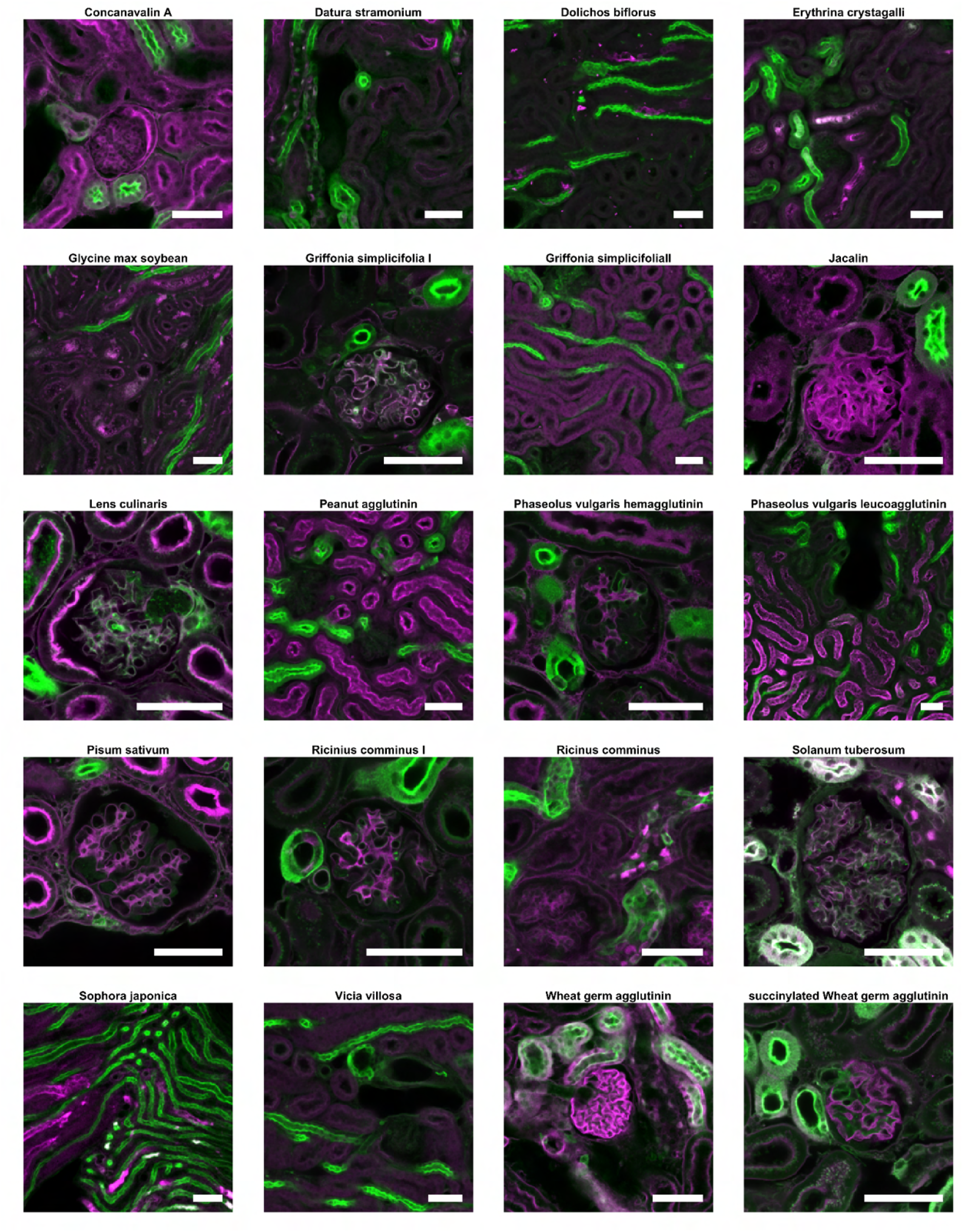
Representative images of lectins histochemistry compatible with INSIHGT performed on mouse kidney tissue. Shown in magenta are the titled lectin’s signals. The signals from *Lycopersicon esculentum* lectin are shown in green for orientation. Scale bars are set to 50 μm.

**Supp Fig.14.**
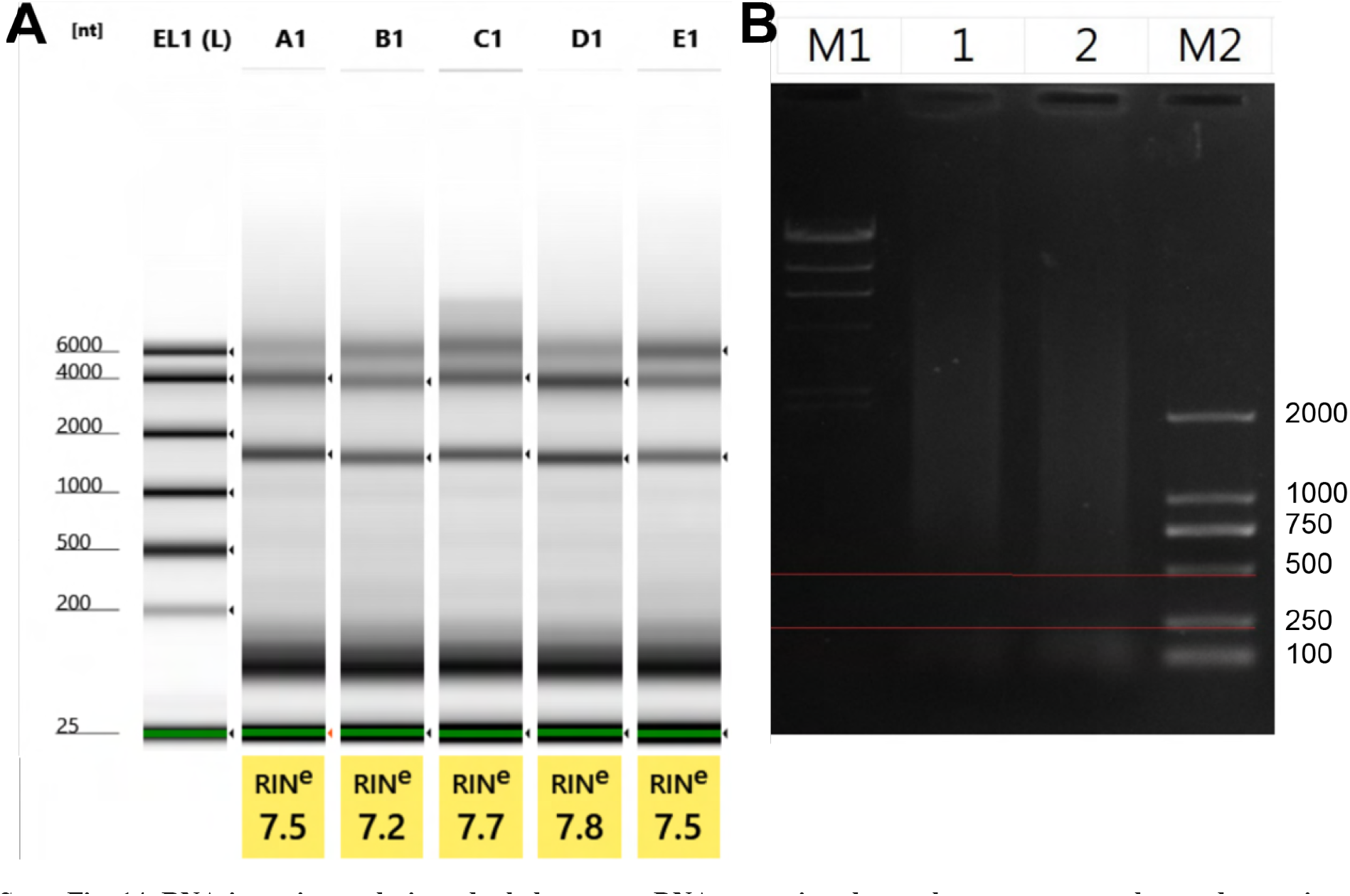
RNA integrity analysis and whole genome DNA extraction electrophoretogram reveals non-destructiveness of INSIHGT. **(A)** RNA integrity number (RIN) analysis of INSIHGT-retrieved tissues. The wells contain samples that are (A1) control, (B1) with delipidation, (C1) with INSIHGT solution A treatment, (D1) with INSIHGT solution B treatment, (E1) with BABB clearing. **(B)** Whole genome DNA extraction electrophoretogram. (M1) λ-Hind Ⅲ digest DNA (Takara), (M2) D2000 (Tiangen). (1) Control, (2) INSIHGT-treated sample following our described full pipeline with 3 days of INSIHGT solution A incubation. DNA total mass for control (14.588 μg) and INSIHGT-treated sample (10.12 μg), and their DNA length (DNAfragment ≥ 500bp) are comparable.

**Supp Fig.15.**
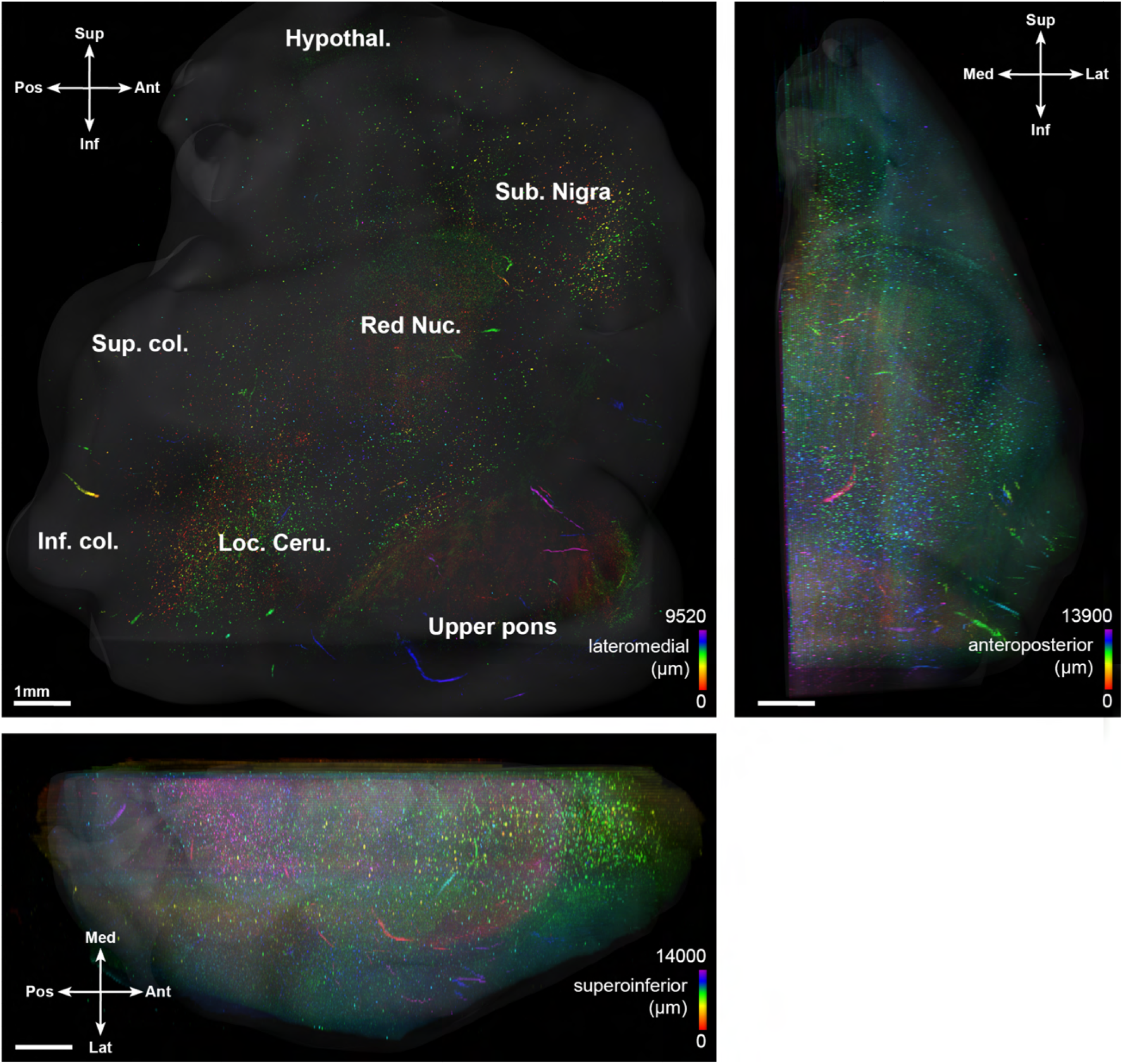
Overview of the human brainstem stained for alpha-synuclein phosphorylated at serine 129 shown in Fig. 5G-I. The INSIHGT immunostaining signal was color-coded according to the dimension going in and out of the paper plane. The surface rendering provides a rough view of the tissue contour. Hypothal.: hypothalamus, Loc. Ceru.: locus ceruleus, Inf. col.: inferior colliculus, Sub. Nigra: substantia nigra, Sup. col., superior colliculus.

**Supp Fig.16.**
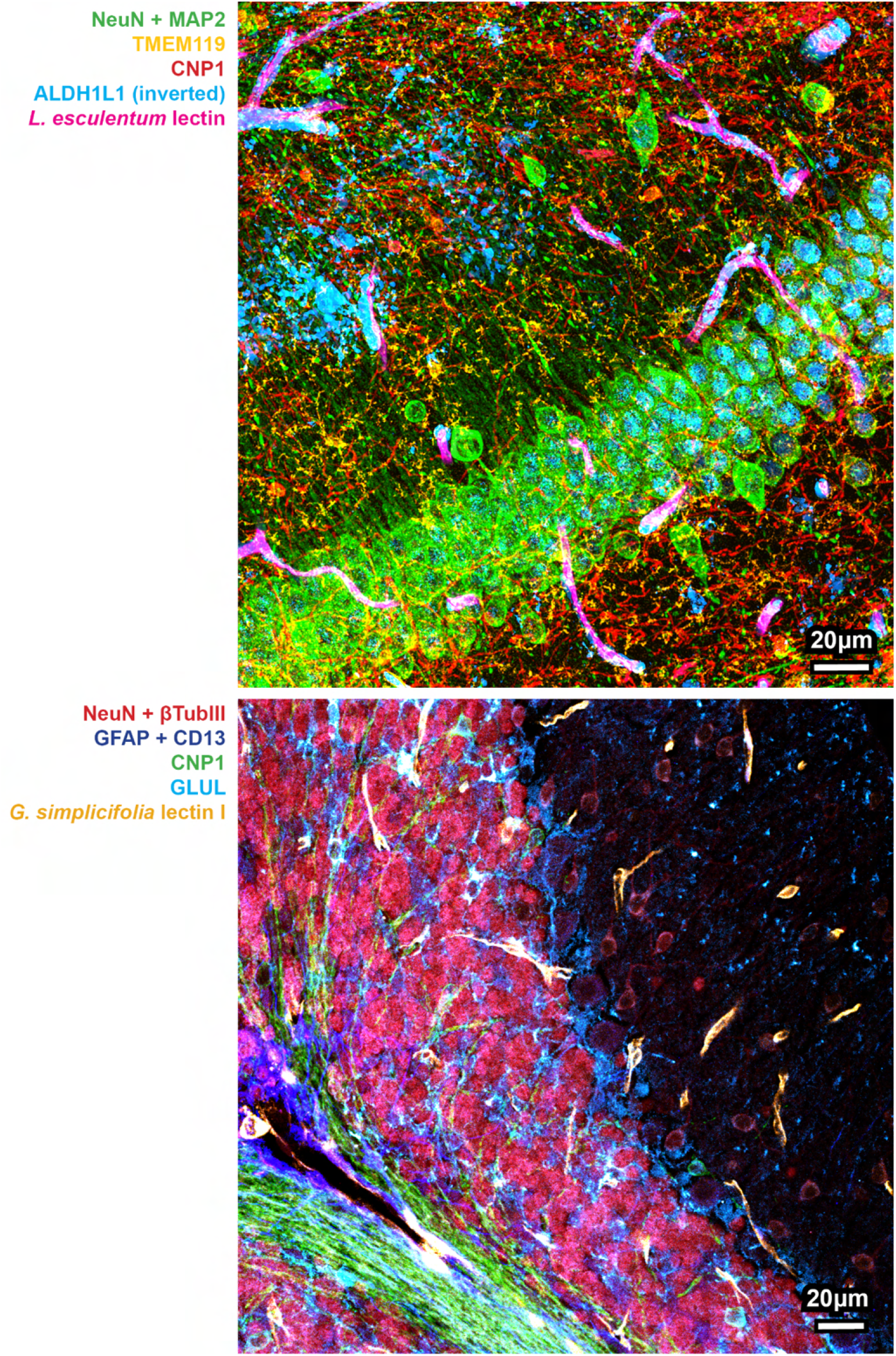
Multiplexing with INSIHGT with high concentrations of antibodies. Upper image: mouse hippocampus stained with 6 protein probes, lower image: mouse cerebellum stained with 7 protein probes. The signal of ALDH1L1 was inverted due to the ubiquitous homogeneous staining of the original signal.

**Supp Fig.17.**
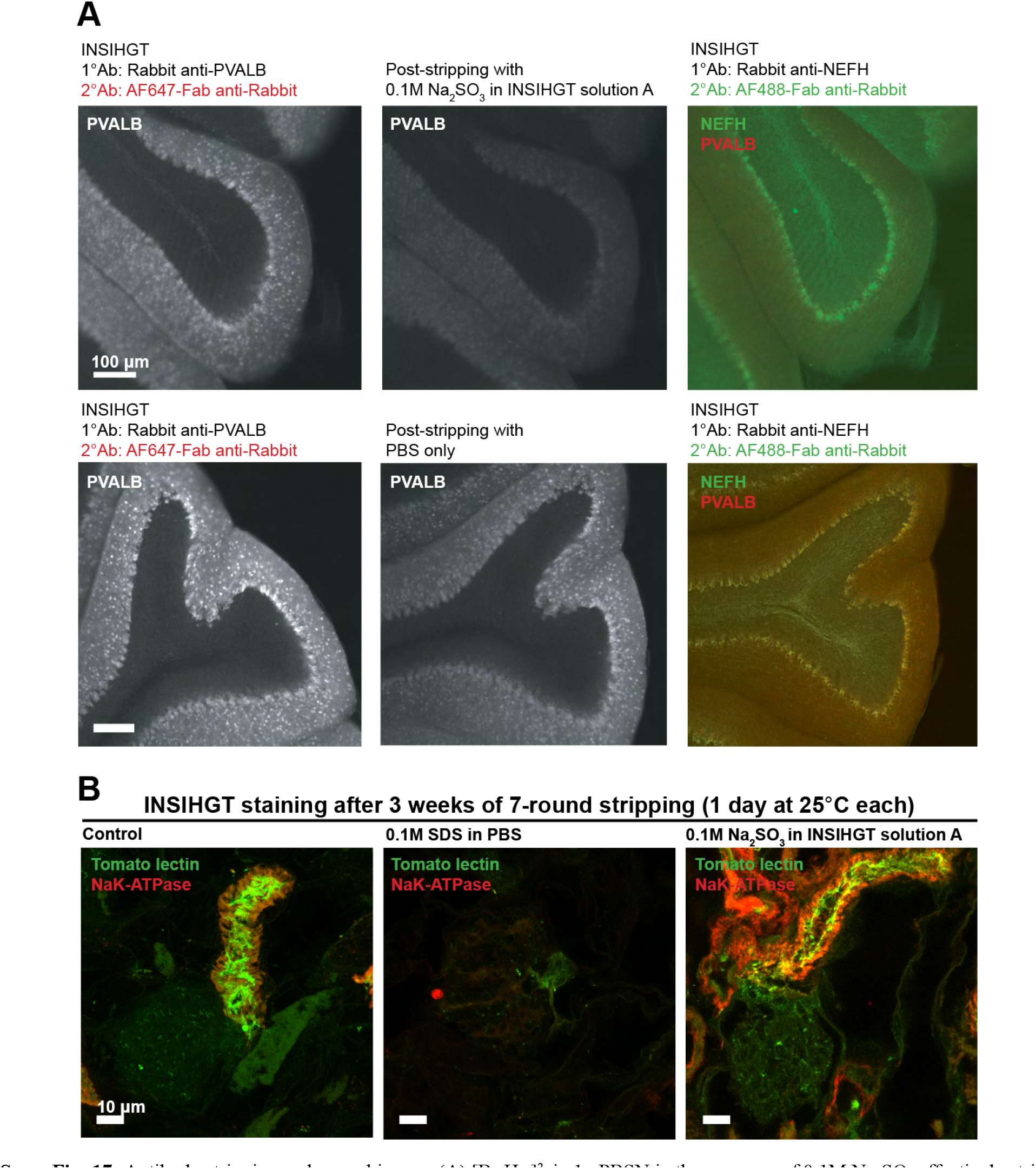
Antibody stripping and re-probing on (**A**) [B_12_H_12_]^2-^ in 1x PBSN in the presence of 0.1M Na_2_SO_3_ effectively strips away the rabbit antibody staining from INSIHGT-stained slices (upper row), while PBS alone would not lead to stripping of antibodies (lower row). After stripping, a second round of INSIHGT-based rabbit anti-NEFH staining was applied to demonstrate the striped rabbit IgGs within the tissue. (**B**) 4% PFA-fixed mouse kidney undergone 7-weeks of stripping with SDS versus Na_2_SO_3_. The total incubation time was 3 weeks. The samples were then re-stained with INSIHGT protocol using tomato lectin (in green) and rabbit anti-NaK-ATPase antibodies (in red).

**Supp Fig.18.**
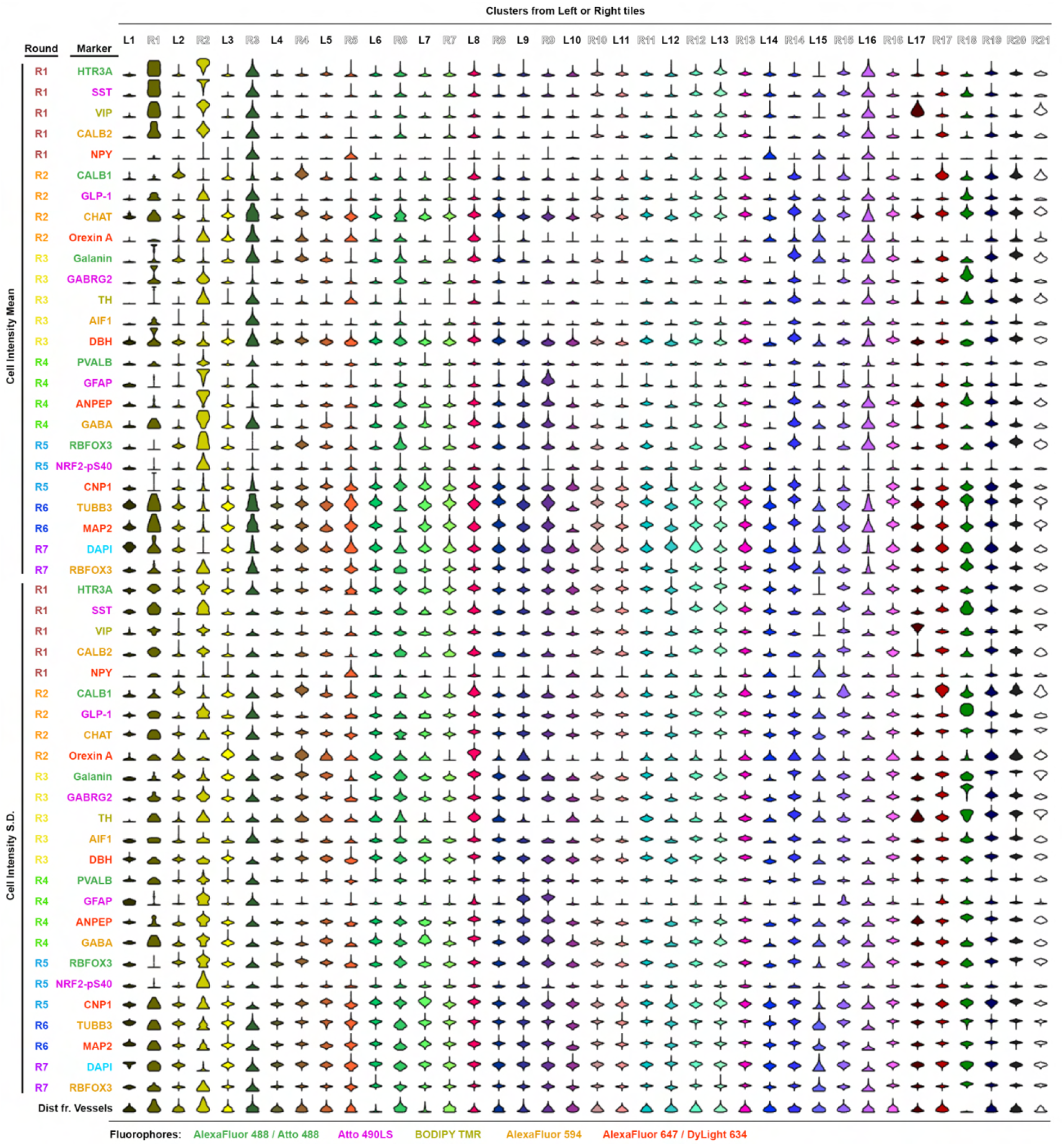
Violin Plots of Each Marker. These plots represent immunostaining mean intensities and their standard deviations (SDs), separately, and the distance from vessels for each cluster obtained by nested UMAP analysis in Figure 6. The text is color-coded based on the fluorophore used in the imaging experiment.

**Supp Fig.19.**
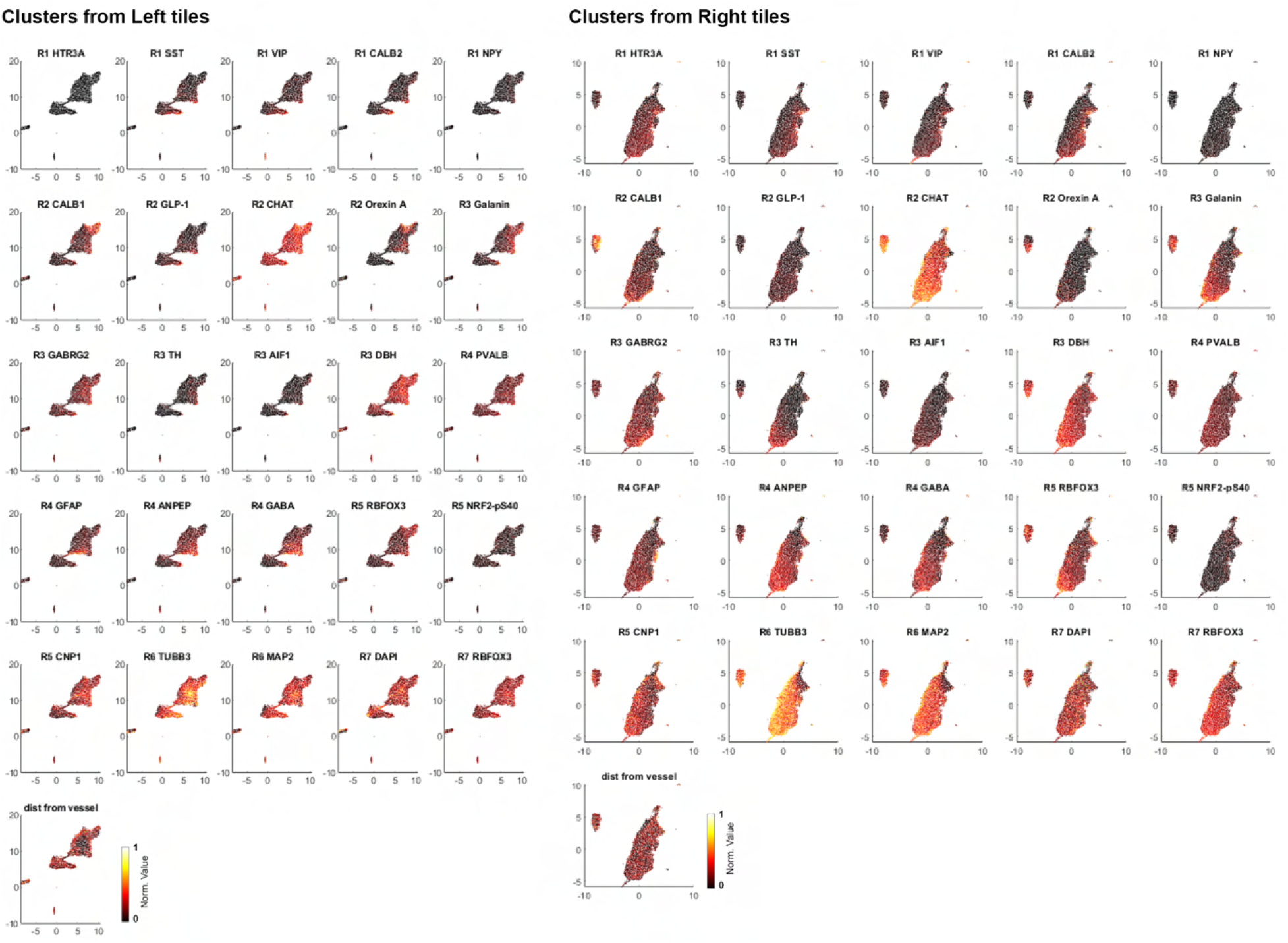
The mean immunostaining intensities of each marker in a 2D-embedded UMAP space.

**Supp Fig.20.**
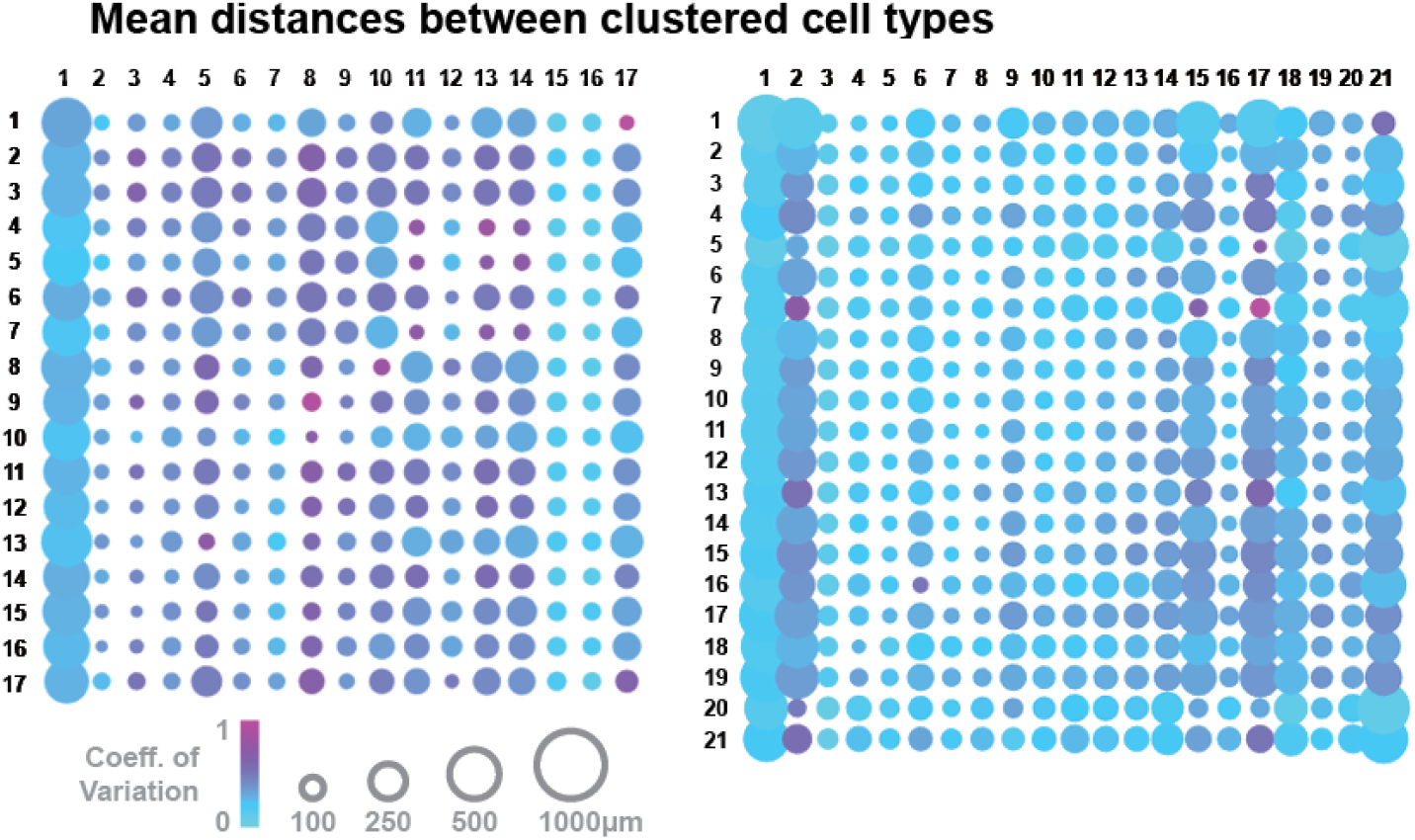
Mean nearest distance matrix between each cell type from each cluster. The size of the dot plot represents the coefficient of variation.

**Supp Fig.21.**
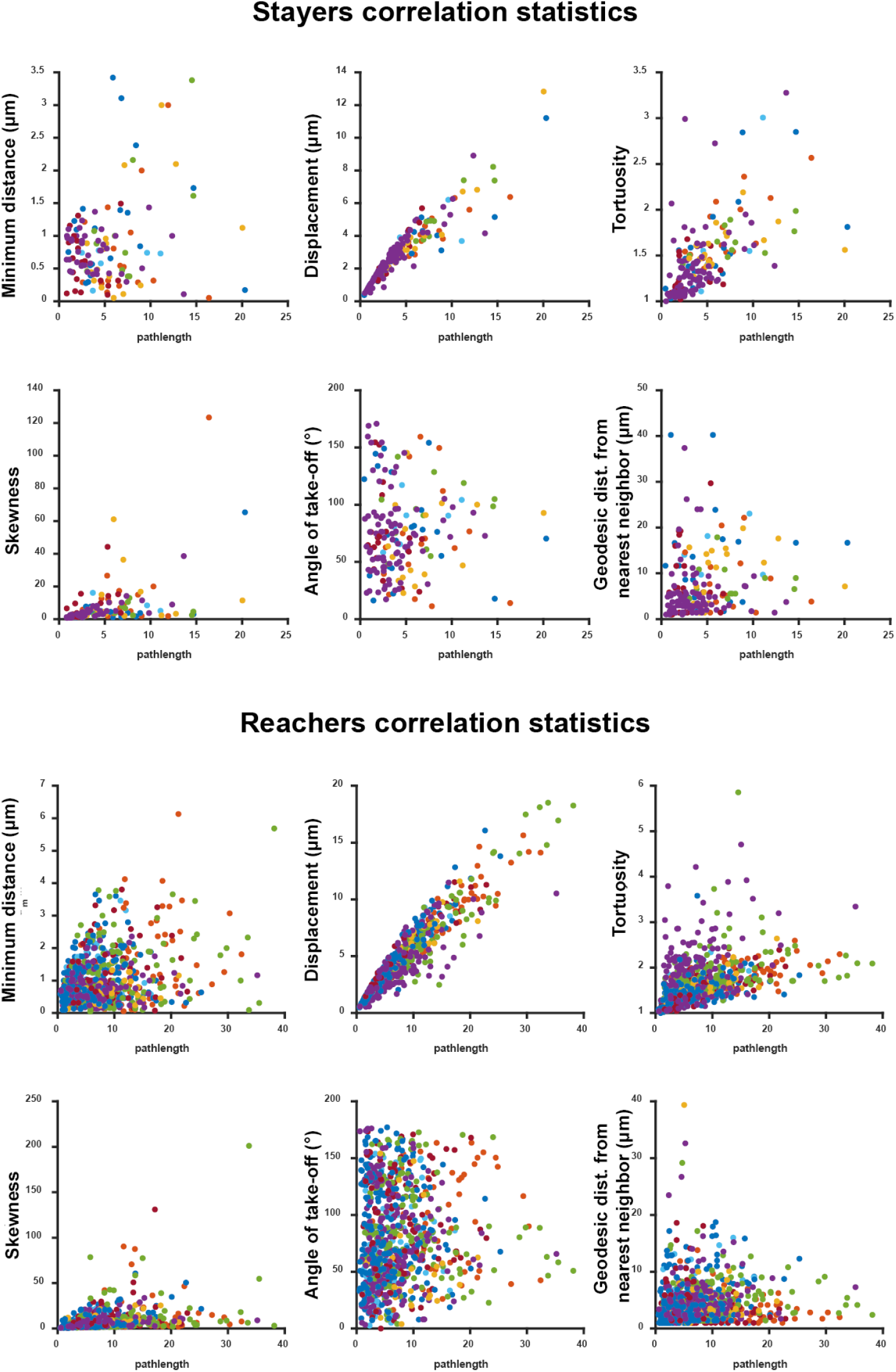
Correlation statistics of physical parameters obtained from 3D images of mouse kidneys, for reacher and stayer clusters respectively. (see also Figure 7).

**Supp Fig.22.**
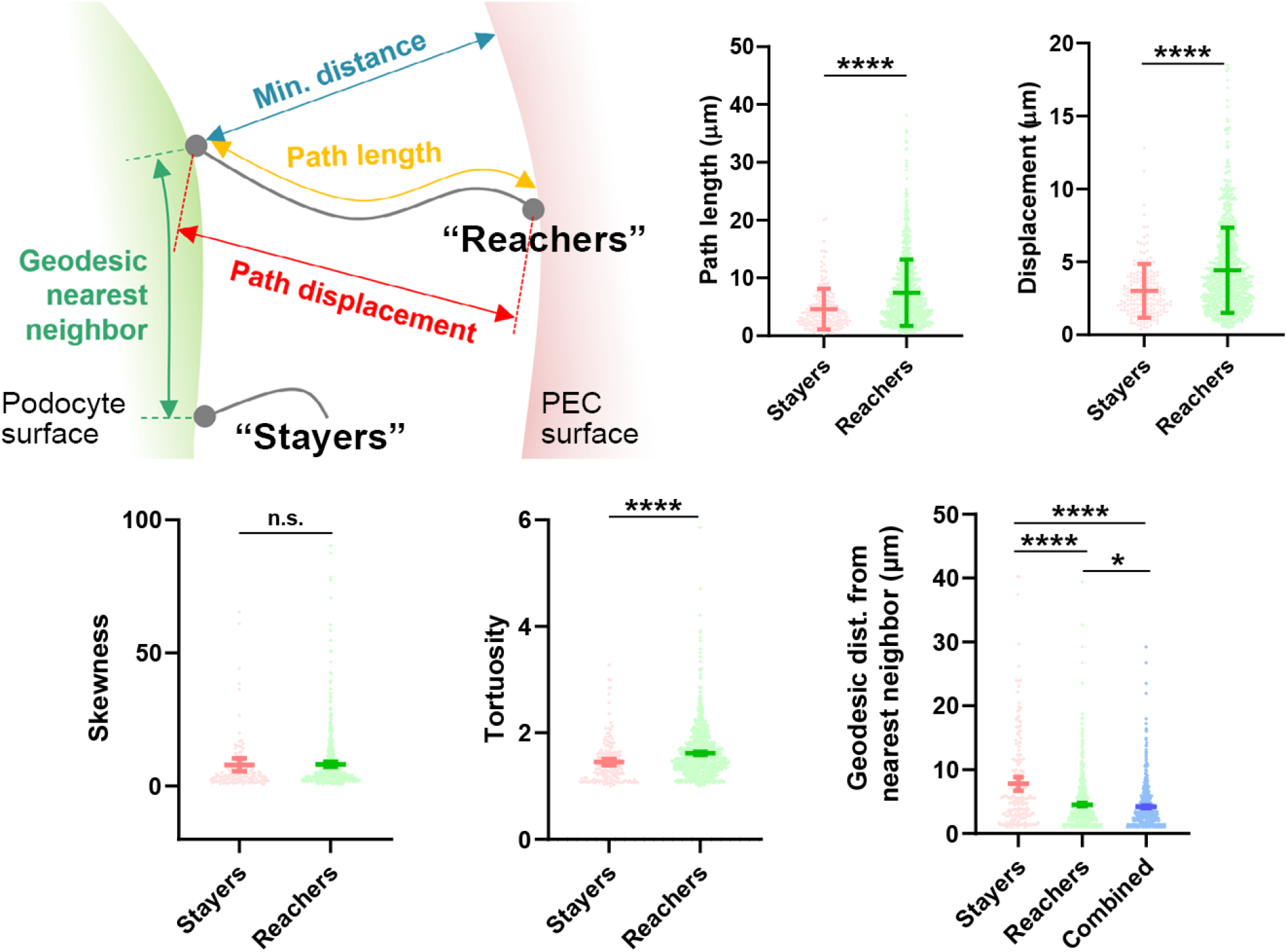
Comparison of morphological and spatial parameters between stayers and reachers clusters obtained from 3D images of mouse kidneys (****: p < 0.0001).

**Supp Fig.23.**
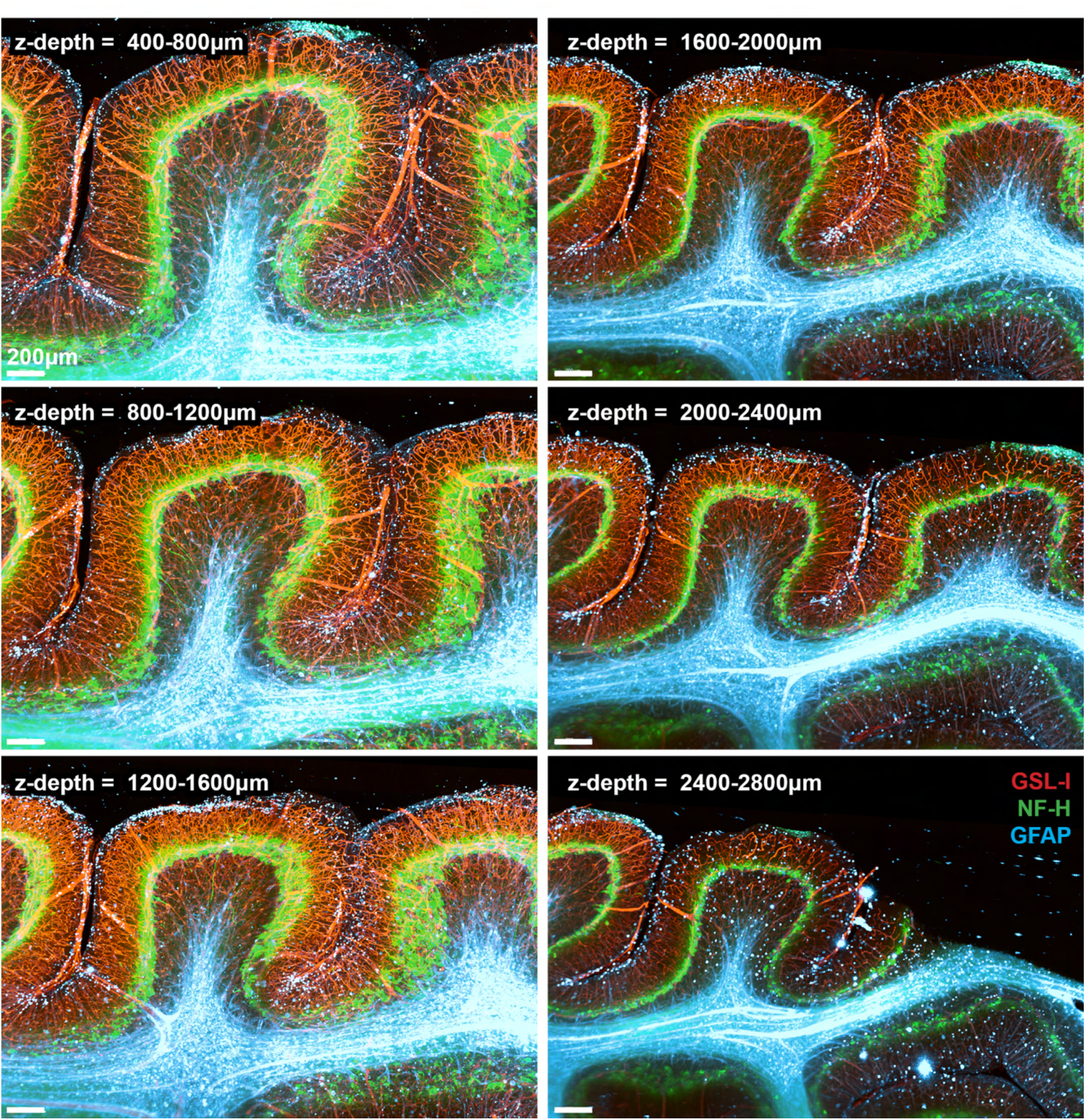
Series of Z-projections of the 4mm × 8mm × 3mm human cerebellum triplex stained for vessels (with *Griffonia simplicifolia* lectin I, GSL-I, in red), neurofilaments (NF-H, in green), and astrocytes (GFAP, in blue).

**Supp Fig.24.**
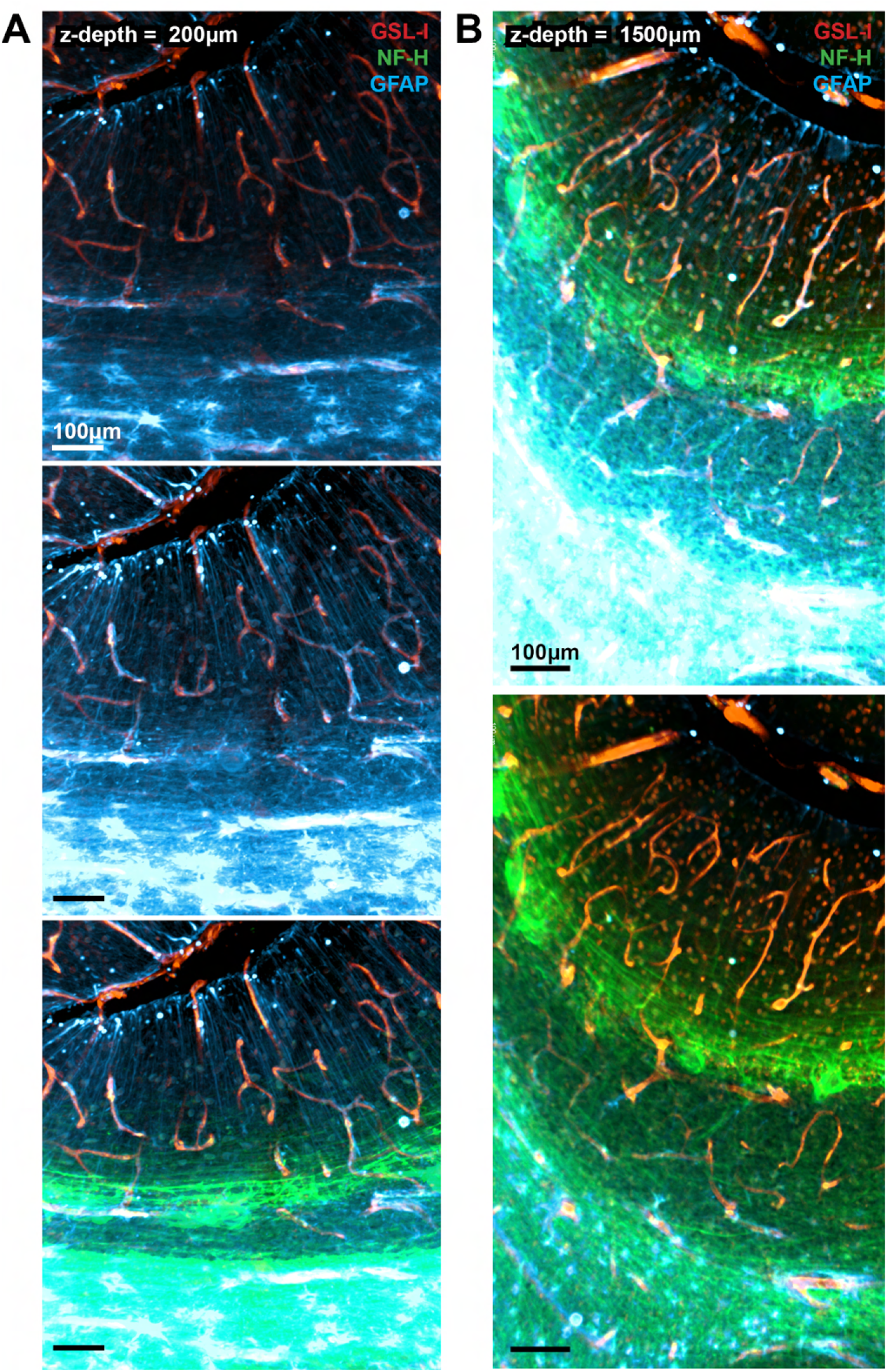
Zoomed-in view of the light-sheet microscopy image in Figure 8. Showing details such as astrocytic endfeets around blood vessels, Bergmann glia processes, and neurofilament fibres in all 3 layers at different *z*-depths. The images were rendered in different dynamic ranges to facilitate visualization of one layer over the other.

**Supp Fig.25.**
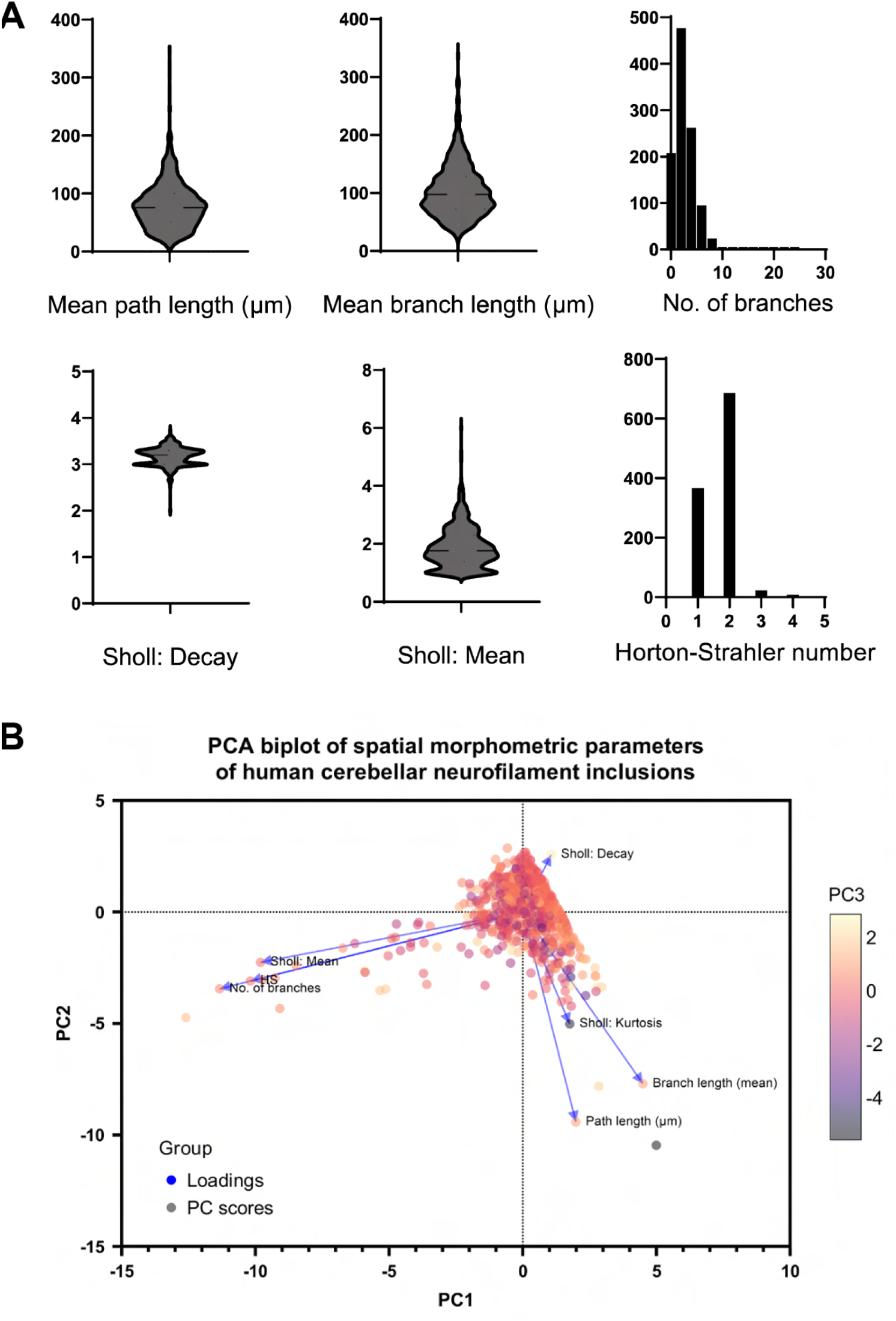
Descriptive statistics of physical parameters obtained from 3D images of human cerebellar neurofilament inclusions, with subsequent principle component analysis with selected parameters. (see also Figure 8).

## Supplementary Movies

**Movie S1.** 3D visualization of the whole mouse brain stained for calcium-binding proteins in Fig. 3J-L.

**Movie S2.** Demonstration of a zoom-in from a centimeter to micrometer scale on a mesoscale-imaged human hemi-brainstem at micrometer resolution of immunostaining signals on phosphorylated alpha-synuclein at serine 129. This is the same specimen detailed in **Fig. 5G-I** and **Fig. S15**.

**Movie S3.** 3D visualization of the 3 mm-thick, 3-plex immunostained human cerebellum in **Fig. 8** and **Fig. S23-25**.

---

**Table S1.** Antibody penetration exponential decay constant of benchmarking experiment using mouse hemi-brains as an estimate of signal inhomogeneity.

| | Inhomogeneity, $\tau \times 10^3$ (confidence interval) |
| --- | --- |
| **INSIHGT** | 2.7 (1.7, 3.7) |
| **CUBIC-HV** | 24.7 (23.3, 26.1) |
| **eFLASH** | 227.8 (210.7, 244.9) |
| **iDISCO** | 48.6 (44.8, 52.4) |
| **SHANEL** | 5.5 (5.2, 5.7) |
| **mELAST** | 136.6 (122.8, 150.4) |

**Table S2.**
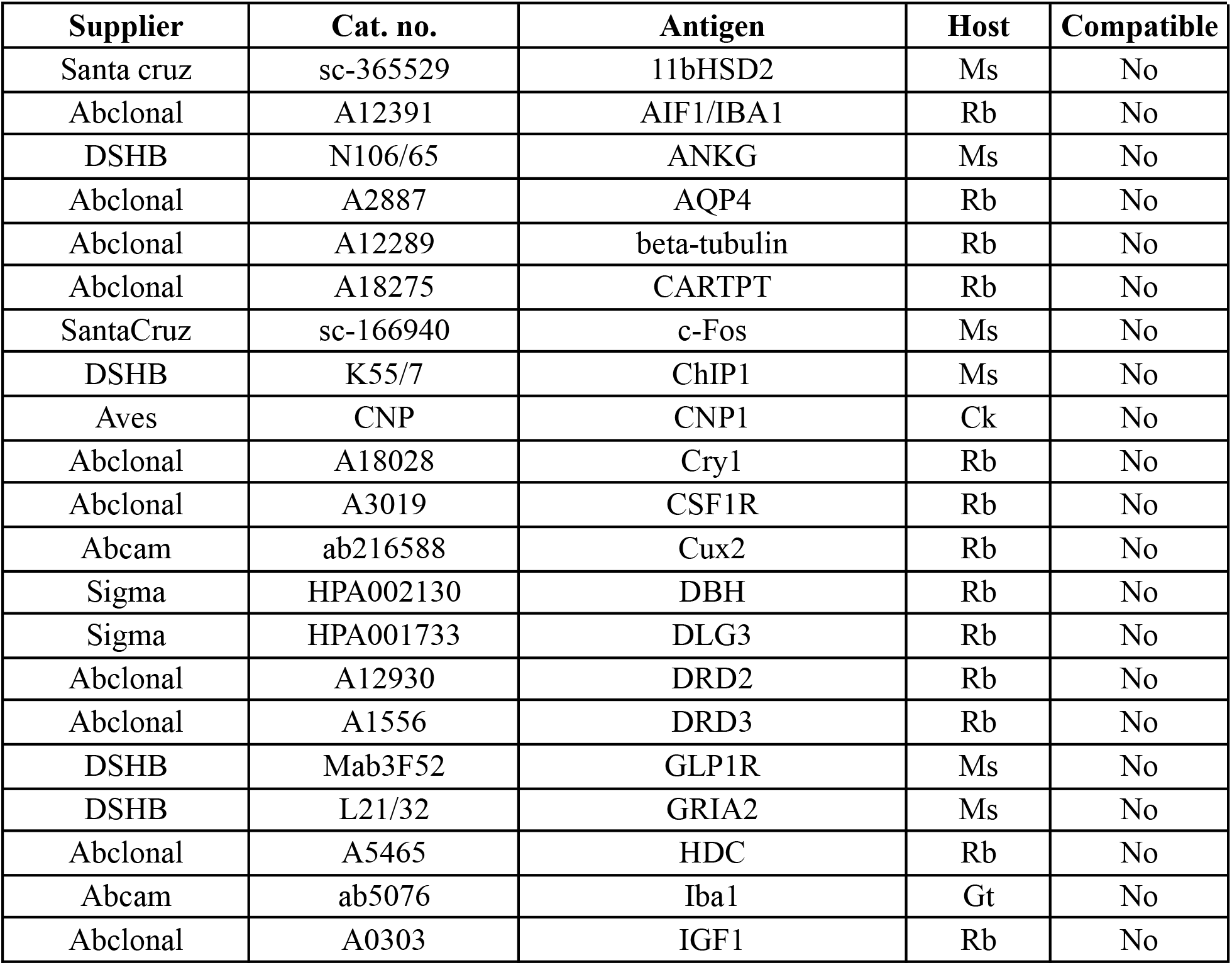
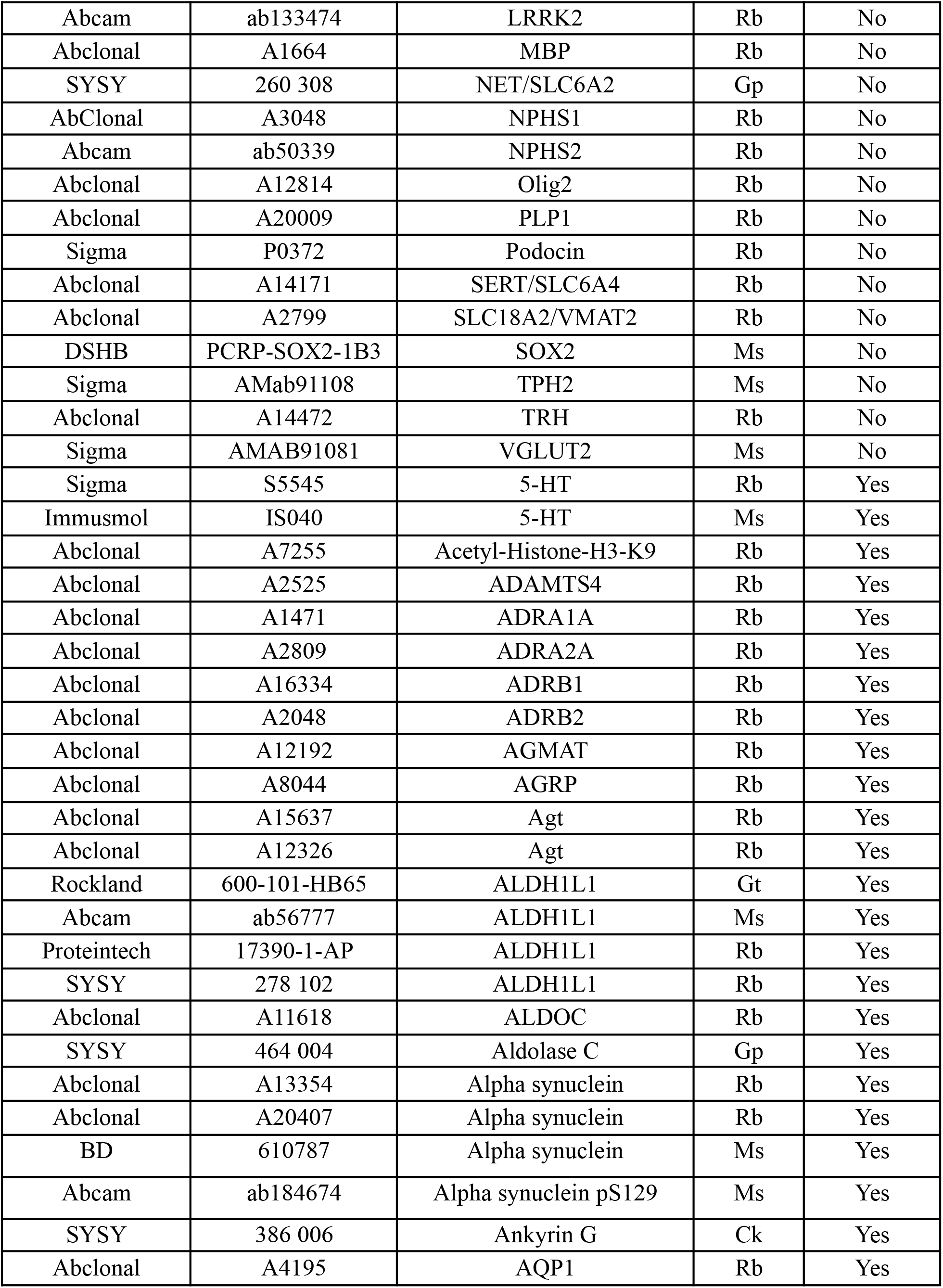
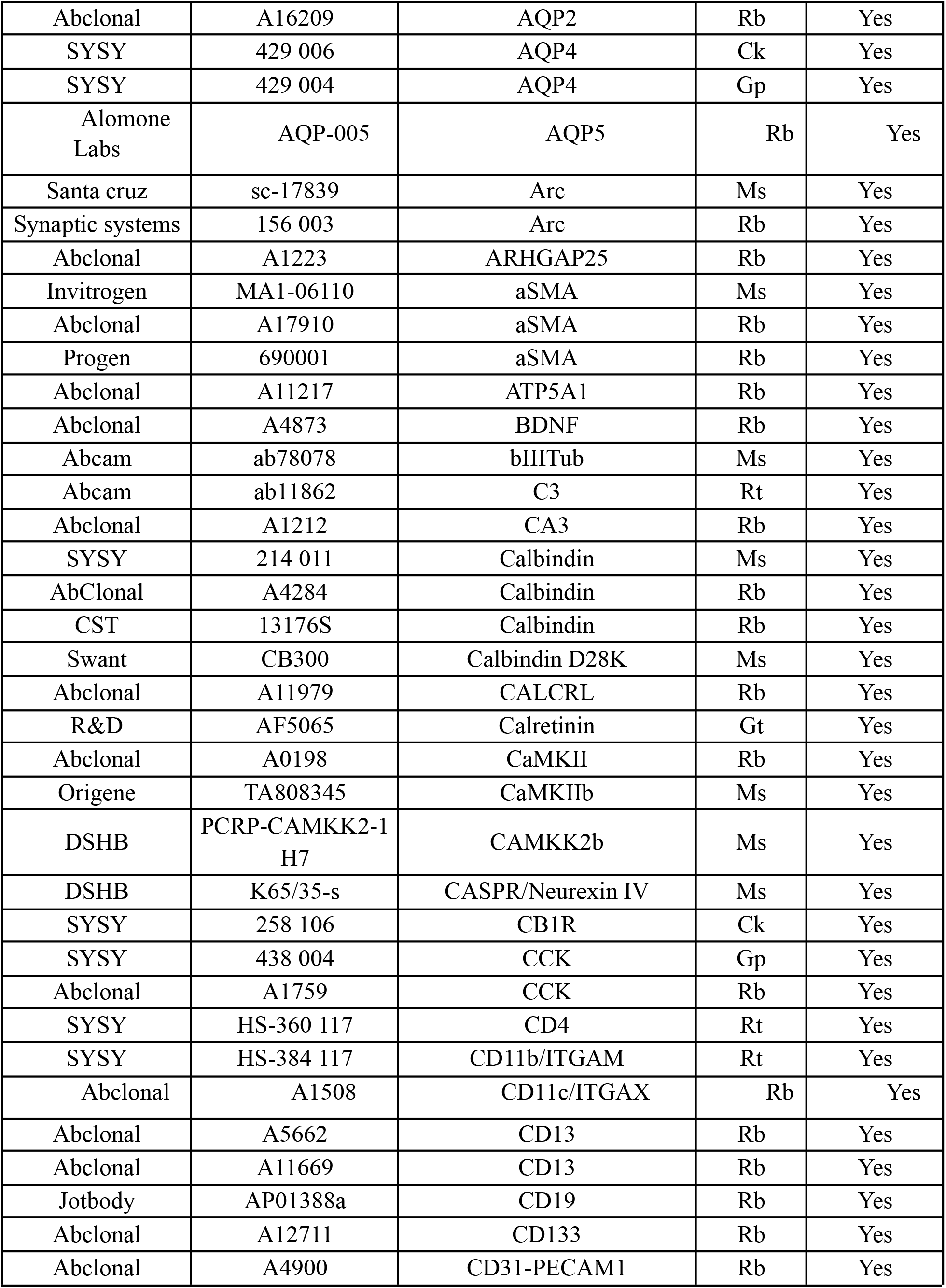
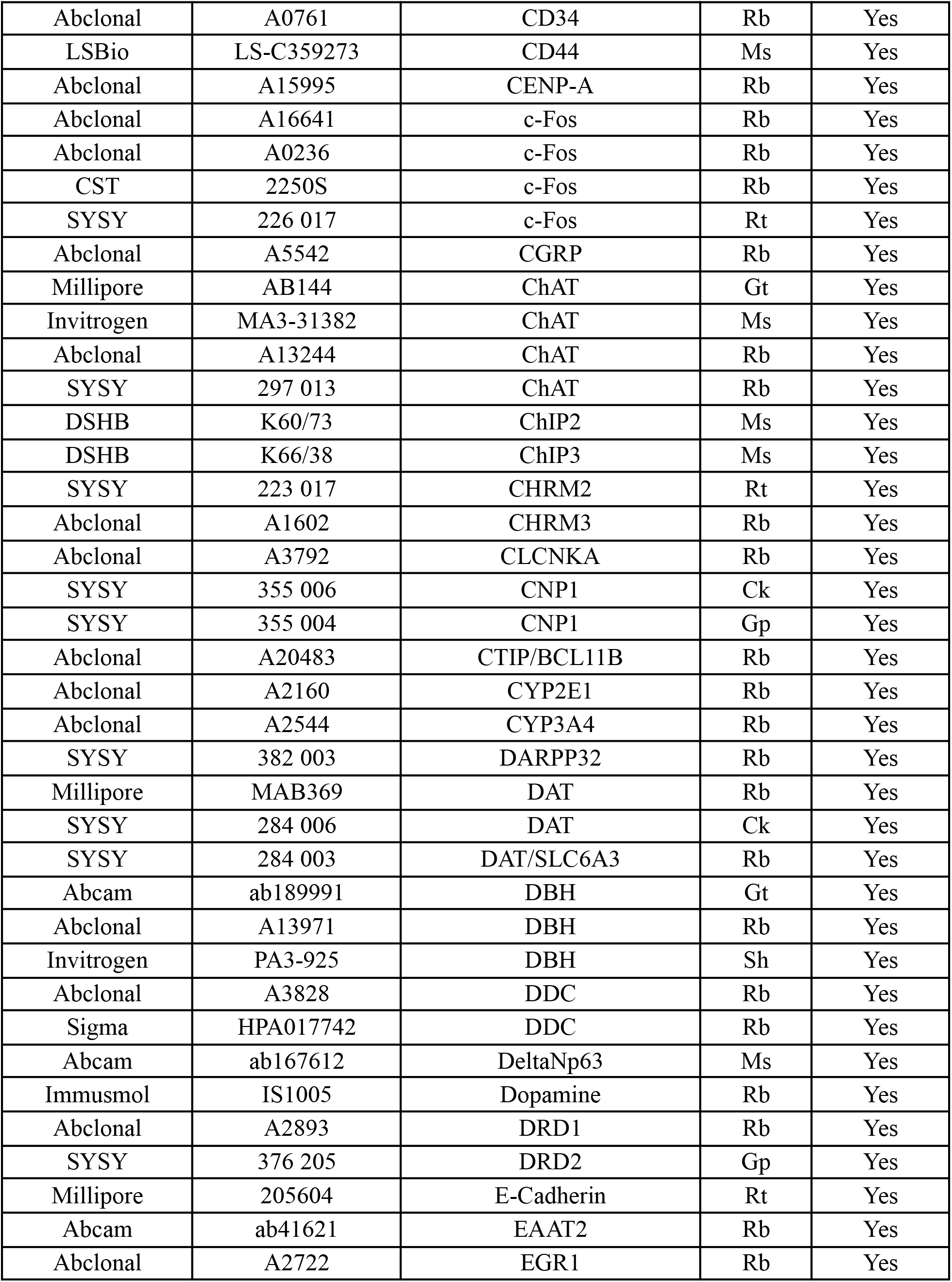
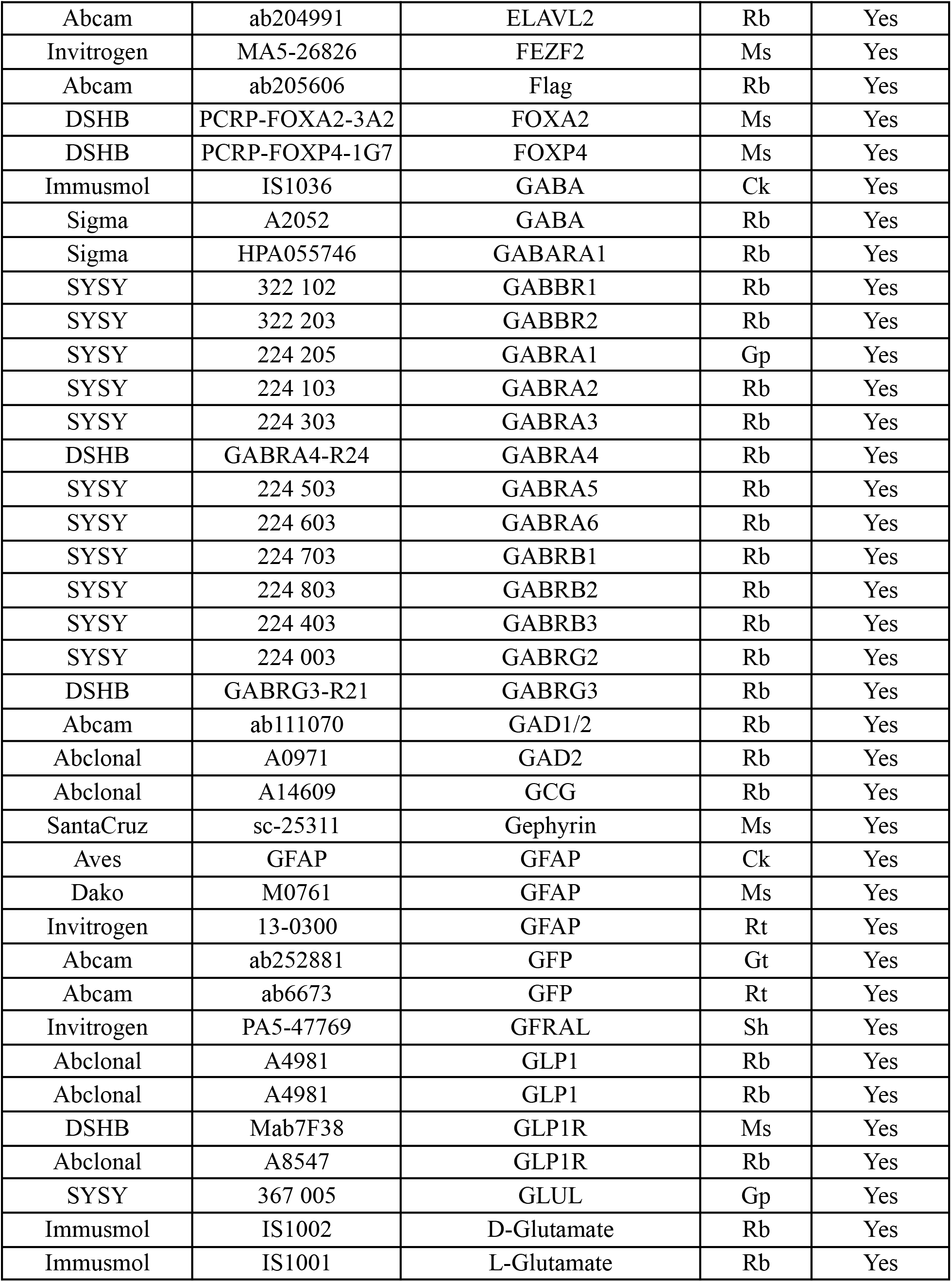
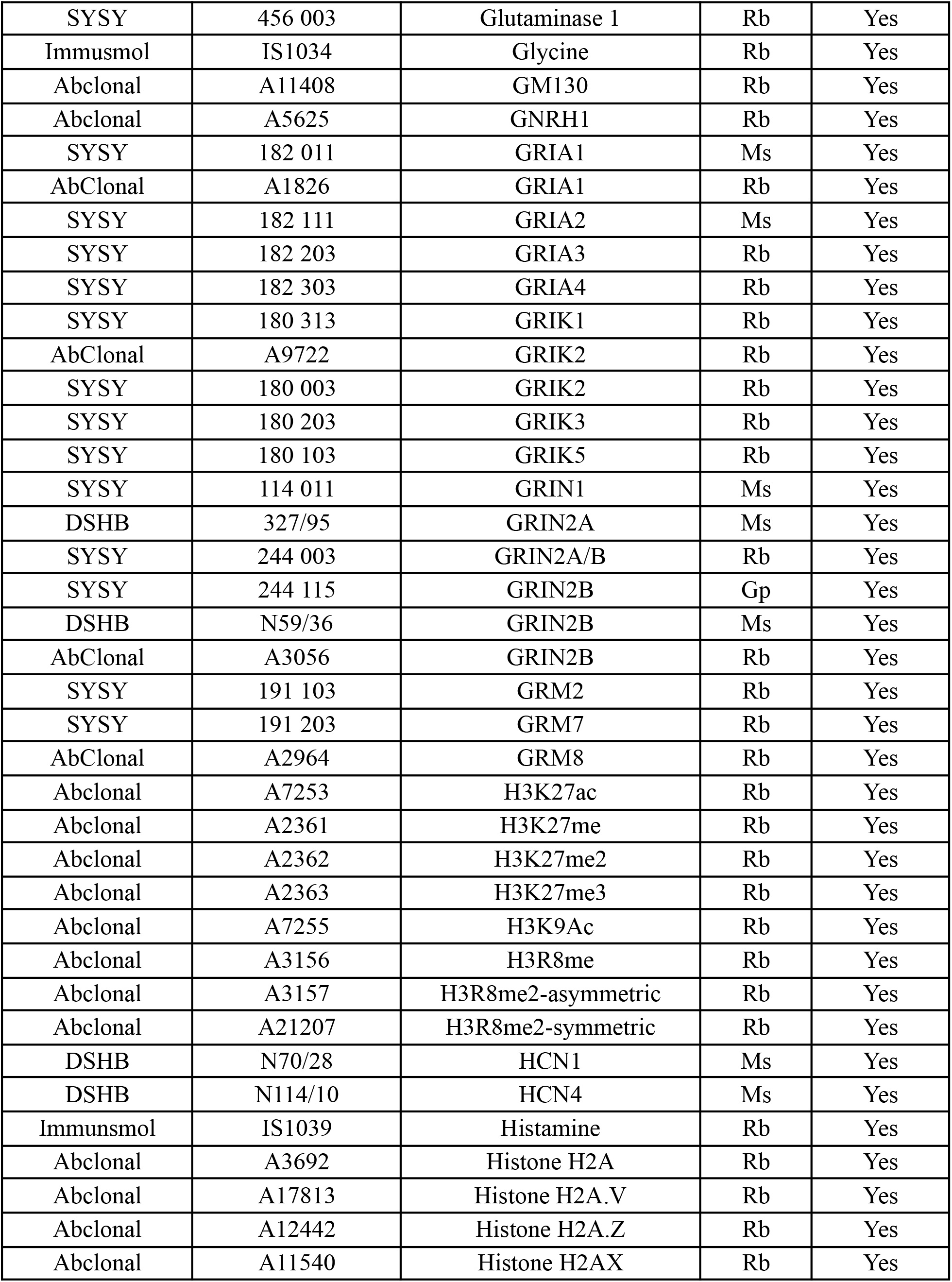
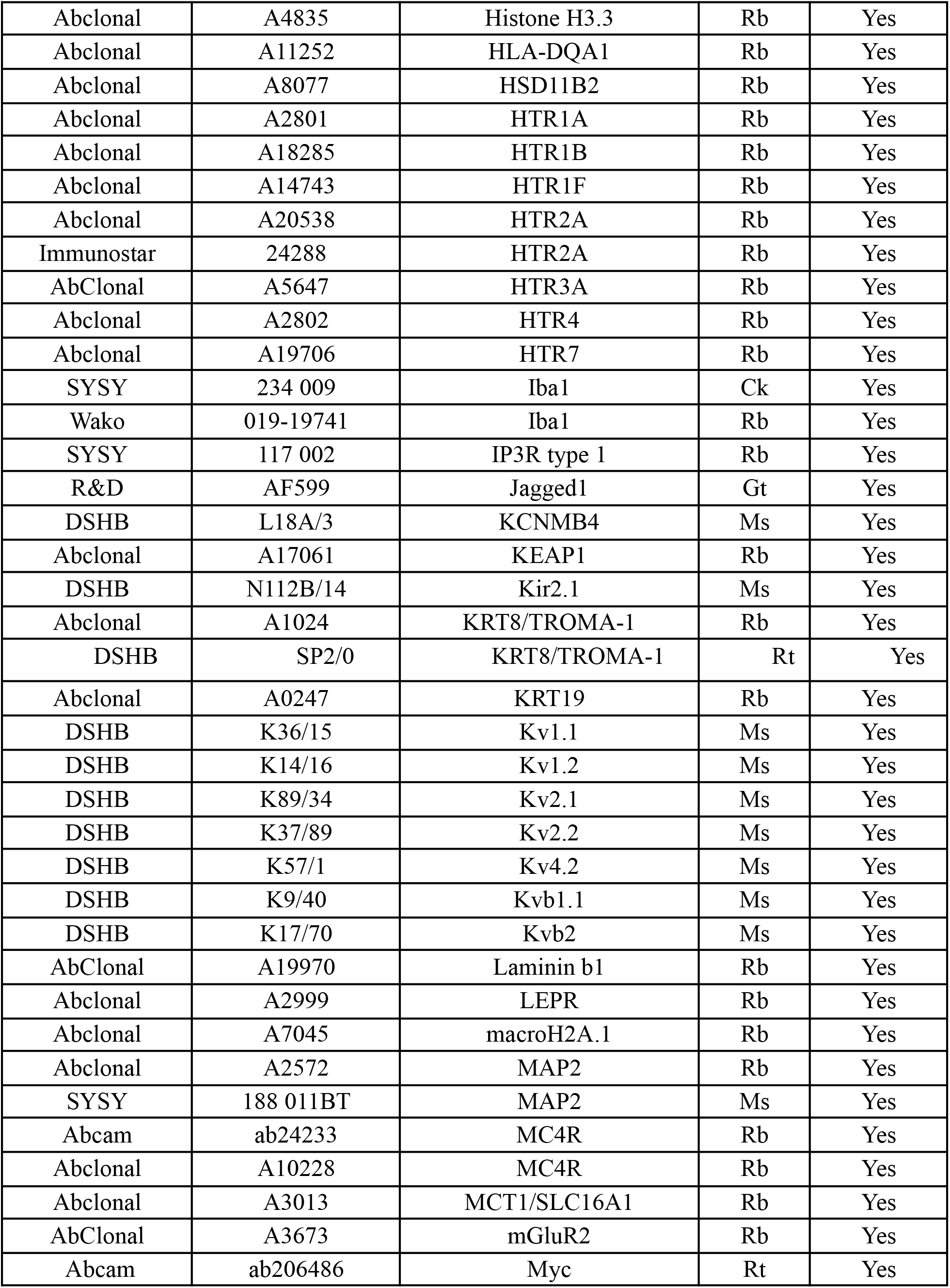
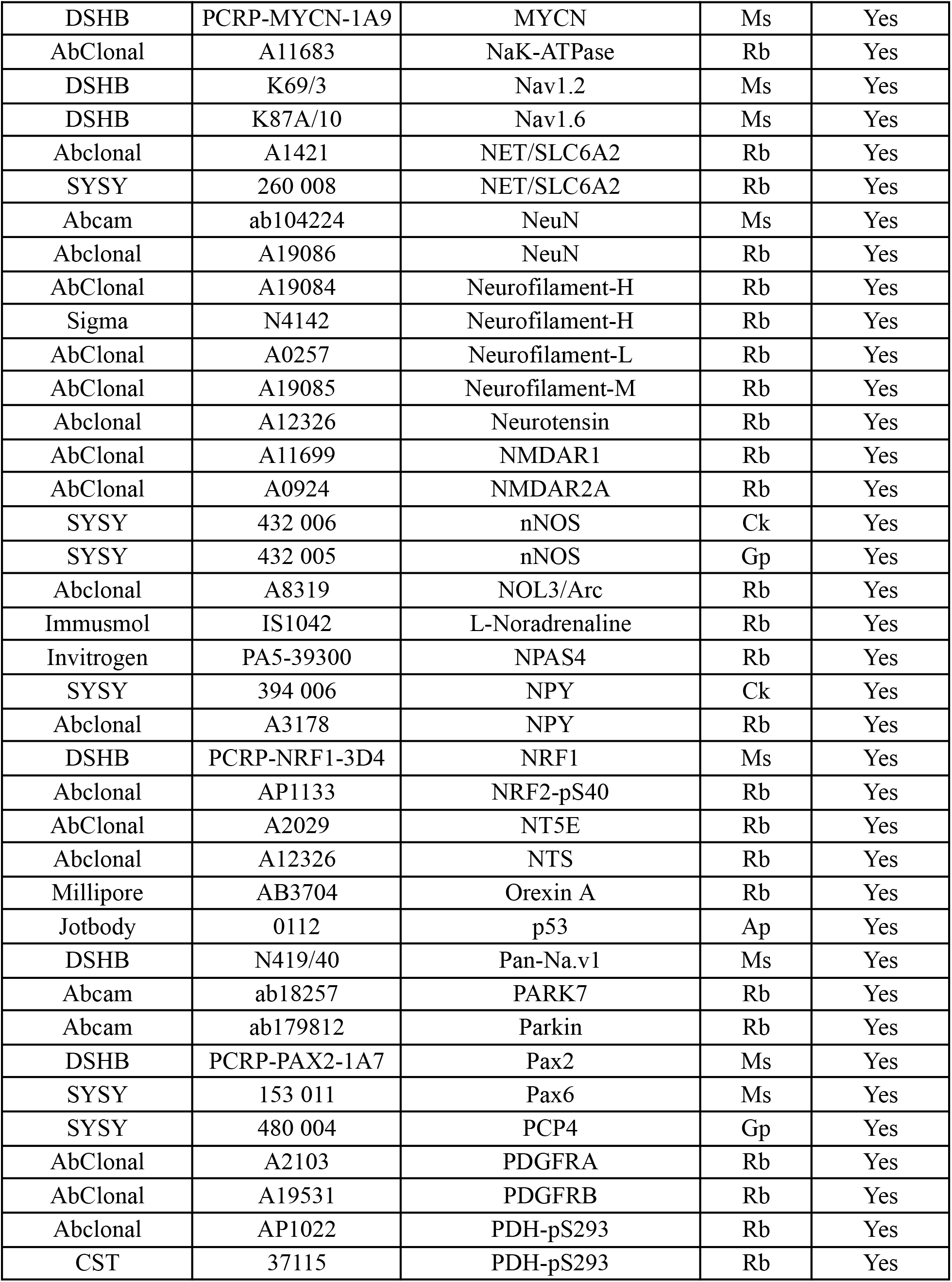
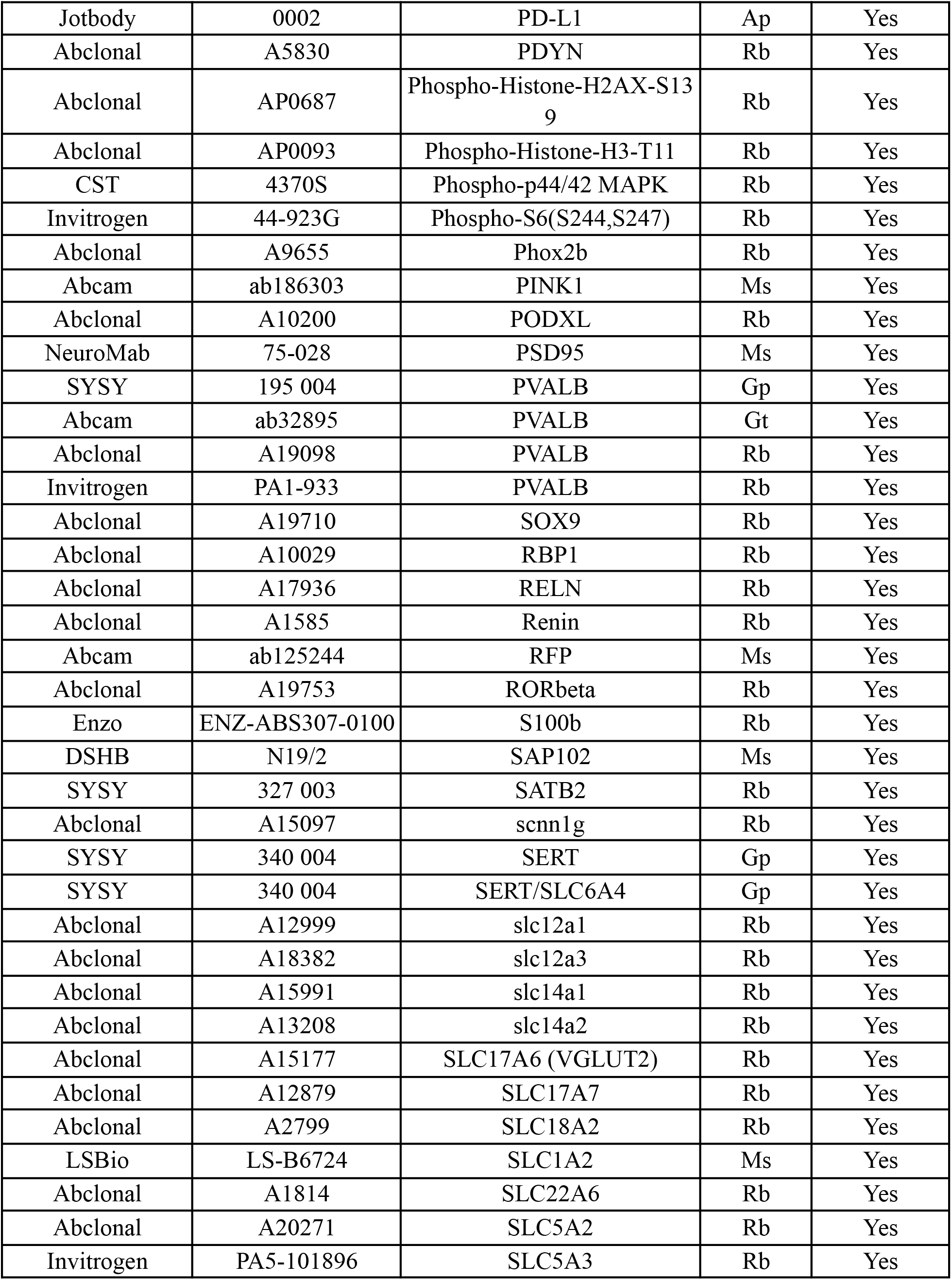
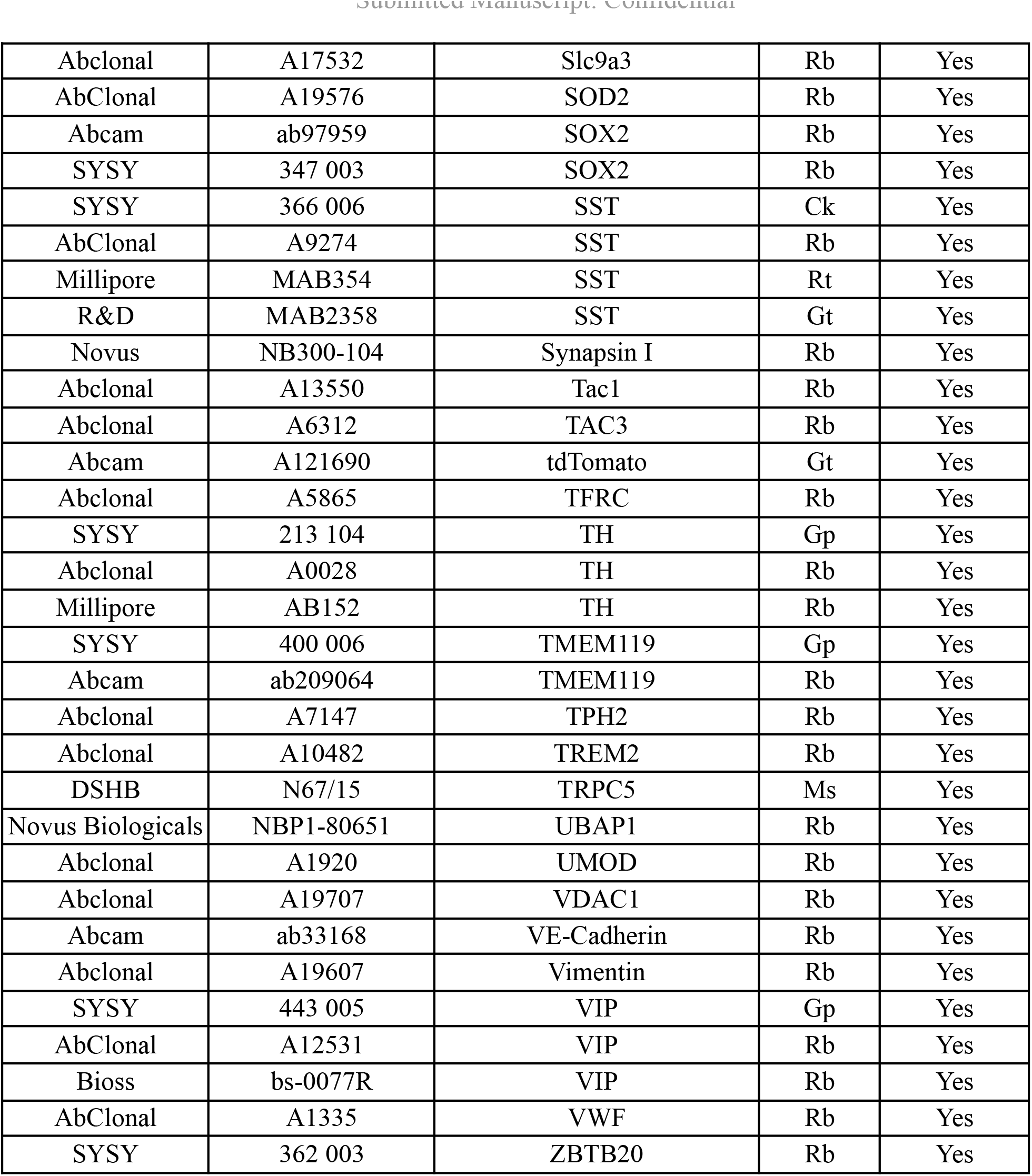
List of antibodies tested with INSIHGT.

## References and Notes

1. Moses, L. & Pachter, L. Museum of spatial transcriptomics. Nat. Methods 19, 534–546 (2022).

2. Hickey, J. W. et al. Spatial mapping of protein composition and tissue organization: a primer for multiplexed antibody-based imaging. Nat. Methods 19, 284–295 (2021).

3. Cho, W., Kim, S. & Park, Y.-G. Towards multiplexed immunofluorescence of 3D tissues. Mol. Brain 16, 37 (2023).

4. Liu, J. T. C., Glaser, A. K., Poudel, C. & Vaughan, J. C. Nondestructive 3D Pathology with Light-Sheet Fluorescence Microscopy for Translational Research and Clinical Assays. Annu. Rev. Anal. Chem. 16, 231–252 (2023).

5. Richardson, D. S. et al. Tissue clearing. Nature Reviews Methods Primers 1, 1–24 (2021).

6. Yau, C. N. et al. Principles of deep immunohistochemistry for 3D histology. Cell Rep Methods 3, 100458 (2023).

7. Ku, T. et al. Elasticizing tissues for reversible shape transformation and accelerated molecular labeling. Nat. Methods 17, 609–613 (2020).

8. Susaki, E. A. et al. Versatile whole-organ/body staining and imaging based on electrolyte-gel properties of biological tissues. Nat. Commun. 11, 1982 (2020).

9. Lai, H. M. et al. Antibody stabilization for thermally accelerated deep immunostaining. Nat. Methods 19, 1137–1146 (2022).

10. Mai, H. et al. Whole-body cellular mapping in mouse using standard IgG antibodies. Nat. Biotechnol. (2023) doi:10.1038/s41587-023-01846-0.

11. Kim, S.-Y. et al. Stochastic electrotransport selectively enhances the transport of highly electromobile molecules. Proceedings of the National Academy of Sciences 112, E6274–E6283 (2015).

12. Yun, D. H., et al. Ultrafast immunostaining of organ-scale tissues for scalable proteomic phenotyping. bioRxiv 660373 (2019) doi:10.1101/660373.

13. Park, J., et al. Integrated platform for multi-scale molecular imaging and phenotyping of the human brain. bioRxiv 2022.03.13.484171 (2023) doi:10.1101/2022.03.13.484171.

14. Murray, E. et al. Simple, Scalable Proteomic Imaging for High-Dimensional Profiling of Intact Systems. Cell 163, 1500–1514 (2015).

15. Park, Y.-G. et al. Protection of tissue physicochemical properties using polyfunctional crosslinkers. Nat. Biotechnol. 37, 73–83 (2018).

16. Pitochelli, A. R. & Hawthorne, F. M. THE ISOLATION OF THE ICOSAHEDRAL B_12_H_12_^-2^ION. J. Am. Chem. Soc. 82, 3228–3229 (1960).

17. Renier, N. et al. iDISCO: A Simple, Rapid Method to Immunolabel Large Tissue Samples for Volume Imaging. Cell 159, 896–910 (2014).

18. Scardigli, M. et al. Comparison of Different Tissue Clearing Methods for Three-Dimensional Reconstruction of Human Brain Cellular Anatomy Using Advanced Imaging Techniques. Front. Neuroanat. 15, 752234 (2021).

19. Darche, M. et al. Light sheet fluorescence microscopy of cleared human eyes. Communications Biology 6, 1–7 (2023).

20. Cartmell, S. C. et al. Multimodal characterization of the human nucleus accumbens. Neuroimage 198, 137–149 (2019).

21. Lichtenegger, A. et al. Assessment of pathological features in Alzheimer’s disease brain tissue with a large field-of-view visible-light optical coherence microscope. Neurophotonics 5, 035002 (2018).

22. Gail Canter, R., et al. 3D mapping reveals network-specific amyloid progression and subcortical susceptibility in mice. Commun Biol 2, 360 (2019).

23. Salvi, G., De Los Rios, P. & Vendruscolo, M. Effective interactions between chaotropic agents and proteins. Proteins 61, 492–499 (2005).

24. Proc, J. L. et al. A quantitative study of the effects of chaotropic agents, surfactants, and solvents on the digestion efficiency of human plasma proteins by trypsin. J. Proteome Res. 9, 5422–5437 (2010).

25. Dugger, B. N. et al. Neuropathological analysis of brainstem cholinergic and catecholaminergic nuclei in relation to rapid eye movement (REM) sleep behaviour disorder. Neuropathol. Appl. Neurobiol. 38, 142–152 (2012).

26. Pachitariu, M. & Stringer, C. Cellpose 2.0: how to train your own model. Nat. Methods 19, 1634–1641 (2022).

27. Shankland, S. J., Smeets, B., Pippin, J. W. & Moeller, M. J. The emergence of the glomerular parietal epithelial cell. Nat. Rev. Nephrol. 10, 158–173 (2014).

28. Li, Z.-H. et al. The Role of Parietal Epithelial Cells in the Pathogenesis of Podocytopathy. Front. Physiol. 13, 832772 (2022).

29. Succar, L., Boadle, R. A., Harris, D. C. & Rangan, G. K. Formation of tight junctions between neighboring podocytes is an early ultrastructural feature in experimental crescentic glomerulonephritis. Int. J. Nephrol. Renovasc. Dis. 9, 297–312 (2016).

30. Neal, C. R. et al. Novel hemodynamic structures in the human glomerulus. Am. J. Physiol. Renal Physiol. 315, F1370–F1384 (2018).

31. Ohse, T. et al. The enigmatic parietal epithelial cell is finally getting noticed: a review. Kidney Int. 76, 1225–1238 (2009).

32. Potter, S. S. Single-cell RNA sequencing for the study of development, physiology and disease. Nat. Rev. Nephrol. 14, 479–492 (2018).

33. Piwecka, M., Rajewsky, N. & Rybak-Wolf, A. Single-cell and spatial transcriptomics: deciphering brain complexity in health and disease. Nat. Rev. Neurol. 19, 346–362 (2023).

34. Tanaka, N. et al. Whole-tissue biopsy phenotyping of three-dimensional tumours reveals patterns of cancer heterogeneity. Nature Biomedical Engineering 1, 796–806 (2017).

35. Xie, W. et al. Prostate Cancer Risk Stratification via Nondestructive 3D Pathology with Deep Learning-Assisted Gland Analysis. Cancer Res. 82, 334–345 (2022).

36. Liu, J. T. C. et al. Harnessing non-destructive 3D pathology. Nat Biomed Eng 5, 203–218 (2021).

37. Seo, J. et al. PICASSO allows ultra-multiplexed fluorescence imaging of spatially overlapping proteins without reference spectra measurements. Nat. Commun. 13, 1–17 (2022).

38. Bai, B. et al. Deep learning-enabled virtual histological staining of biological samples. Light Sci Appl 12, 57 (2023).

39. Suhling, K., French, P. M. W. & Phillips, D. Time-resolved fluorescence microscopy. Photochem. Photobiol. Sci. 4, 13–22 (2005).

40. Buchwalow, I., Samoilova, V., Boecker, W. & Tiemann, M. Multiple immunolabeling with antibodies from the same host species in combination with tyramide signal amplification. Acta Histochem. 120, 405–411 (2018).

41. Dahiya, V. & Chaudhuri, T. K. Chaperones GroEL/GroES accelerate the refolding of a multidomain protein through modulating on-pathway intermediates. J. Biol. Chem. 289, 286–298 (2014).

42. Mai, H. et al. Scalable tissue labeling and clearing of intact human organs. Nat. Protoc. 17, 2188–2215 (2022).

43. Voigt, F. F. et al. The mesoSPIM initiative: open-source light-sheet microscopes for imaging cleared tissue. Nat. Methods 16, 1105–1108 (2019).

44. Hörl, D. et al. BigStitcher: reconstructing high-resolution image datasets of cleared and expanded samples. Nat. Methods 16, 870–874 (2019).

45. Arshadi, C., Günther, U., Eddison, M., Harrington, K. I. S. & Ferreira, T. A. SNT: a unifying toolbox for quantification of neuronal anatomy. Nat. Methods 18, 374–377 (2021).

46. Soille, P. Morphological Image Analysis. (Springer Berlin Heidelberg).

47. Bigun, J. Optimal Orientation Detection of Linear Symmetry. 433–438 (1987).

48. Khan, A. R. et al. 3D structure tensor analysis of light microscopy data for validating diffusion MRI. Neuroimage 111, 192–203 (2015).

49. Bigun, J., Bigun, T. & Nilsson, K. Recognition by symmetry derivatives and the generalized structure tensor. IEEE Trans. Pattern Anal. Mach. Intell. 26, 1590–1605 (2004).

50. Website. Connor Meehan, Jonathan Ebrahimian, Wayne Moore, and Stephen Meehan (2022). Uniform Manifold Approximation and Projection (UMAP) (https://www.mathworks.com/matlabcentral/fileexchange/71902), MATLAB Central File Exchange.

51. Morel, P. Gramm: grammar of graphics plotting in Matlab. Journal of Open Source Software 3, 568 (2018).

52. Masselink, W. et al. Broad applicability of a streamlined ethyl cinnamate-based clearing procedure. Development 146, dev166884 (2019).

53. Klingberg, A. et al. Fully automated evaluation of total glomerular number and capillary tuft size in nephritic kidneys using lightsheet microscopy. J. Am. Soc. Nephrol. 28, 452–459 (2017).

54. Becker, K., Jährling, N., Saghafi, S., Weiler, R. & Dodt, H.-U. Chemical Clearing and Dehydration of GFP Expressing Mouse Brains. PLoS One 7, e33916 (2012).

55. Jacso, T. et al. The Mechanism of Denaturation and the Unfolded State of the α-Helical Membrane-Associated Protein Mistic. (2013) doi:10.1021/ja408644f.

56. Ganguly, P., Boserman, P., van der Vegt, N. F. A. & Shea, J.-E. Trimethylamine N-oxide Counteracts Urea Denaturation by Inhibiting Protein–Urea Preferential Interaction. (2017) doi:10.1021/jacs.7b11695.

57. Riddlestone, I. M., Kraft, A., Schaefer, J. & Krossing, I. Taming the Cationic Beast: Novel Developments in the Synthesis and Application of Weakly Coordinating Anions. Angew. Chem. Int. Ed. 57, 13982–14024 (2018).

58. Lipping, L. et al. Superacidity of closo-Dodecaborate-Based Brønsted Acids: a DFT Study. (2015) doi:10.1021/jp506485x.

59. Assaf, K. I. & Nau, W. M. The Chaotropic Effect as an Assembly Motif in Chemistry. Angew. Chem. Int. Ed Engl. 57, 13968–13981 (2018).

60. Assaf, K. I., et al. Water Structure Recovery in Chaotropic Anion Recognition: High-Affinity Binding of Dodecaborate Clusters to γ-Cyclodextrin. Angew. Chem. Int. Ed. 54, 6852–6856 (2015).

61. Karki, K., Gabel, D. & Roccatano, D. Structure and dynamics of dodecaborate clusters in water. Inorg. Chem. 51, 4894–4896 (2012).

62. Jiang, Y. et al. Highly Structured Water Networks in Microhydrated Dodecaborate Clusters. J. Phys. Chem. Lett. 13, 11787–11794 (2022).

63. Jiang, Y. et al. Unraveling hydridic-to-protonic dihydrogen bond predominance in monohydrated dodecaborate clusters. Chem. Sci. 13, 9855–9860 (2022).

64. Bukovsky, E. V. et al. Comparison of the Coordination of B12F122–, B12Cl122–, and B12H122– to Na+ in the Solid State: Crystal Structures and Thermal Behavior of Na2(B12F12), Na2(H2O)4(B12F12), Na2(B12Cl12), and Na2(H2O)6(B12Cl12). (2017) doi:10.1021/acs.inorgchem.6b02920.

65. Goszczyński, T. M., Fink, K., Kowalski, K., Leśnikowski, Z. J. & Boratyński, J. Interactions of Boron Clusters and their Derivatives with Serum Albumin. Sci. Rep. 7, 9800 (2017).

